# LAYERED GENOMIC COMPOSITION AND LINGUISTIC CONTINUITY IN GRECÌA SALENTINA

**DOI:** 10.64898/2026.09.16.751136

**Authors:** Giulia Cangialosi, Martina Gulì, Francesca Menato, Gerardo Pepe, Ester Mercuri, Francesco Zangaro, Carla Jodice, Fulvio Cruciani, Beniamino Trombetta, Manuela Helmer-Citterich, Nejat Akar, Cristina Guardiano, Giuseppe Longobardi, Valeria Specchia, Ornella Semino, Eugenia D’Atanasio, Andrea Novelletto

## Abstract

Grecìa Salentina (GS) is the label assigned to a geographic area in Southern Italy where a Greek dialect has historically been spoken. The origins of this Greek-speaking community, whether dating from ancient or Medieval settlement, are a controversial and essentially unsolved issue.

To evaluate which historical-linguistic scenario is best supported on genomic grounds, we used whole genome sequencing of 27 GS natives and 21 controls, embedding them in a 799 modern samples dataset in which Greece and Anatolia were evenly represented, an uncommon feature in the genomic literature on the area. We considered different aspects of the genetic structuring, at different spatial and temporal scales. Analyses also included 4021 ancient samples from the literature.

All analyses pointed to the same conclusion: the GS genomic profile displays the same characteristic features of the rest of Southern Italy, with no definite evidence for a single source population from outside Italy. Correspondingly, we detected strong genetic connectivity among GS municipalities and between GS and the rest of Italy. Only a subset of our analyses suggested a slightly increased affinity of GS with Greece, or with the Anatolian peninsula. These results indicate that GS has been all but a genetically closed community, and that the Greek language did not act as a barrier to gene flow. By endorsing a “transient mobility model” we support the most recent linguistic hypothesis: the prolonged resilience of the Greek language in GS, maintained *in situ* and periodically rejuvenated through recurrent external inputs from the Greek and Byzantine worlds.

## Introduction

Salento, the extreme southern part of Apulia, is a peninsula at the south-eastern edge of Italy, which separates the Adriatic Sea (to the East) from the Gulf of Taranto (to the West). ‘Grecìa Salentina’ (henceforth GS) is the label assigned to a geographic area in Salento inhabited by an officially recognized (https://libertaciviliimmigrazione.dlci.interno.gov.it/minoranze) and constitutionally protected linguistic minority. In GS a Greek dialect (Griko) is being spoken (although it is now less widespread than in the past). GS coincides with 12 municipalities (https://www.unionegreciasalentina.le.it), 10 of which are considered Greek-speaking (ellenofoni, see Supplemental Text). These are adjacent to each other, with their main centres separated by some tens of kilometres in an essentially flat territory. The overall census population reaches approximately 43,000. The origins of this community and of its linguistic peculiarity are poorly known.

Anecdotal and historical accounts of population events impacting the territory of today’s GS are scanty. This contrasts sharply with the precise reconstructions of the settlement, zones of influence and rivalries of Greek colonies elsewhere in Southern Italy before the imposition of the Roman rule. After Romanization, the cities of Brindisi and Otranto were the leading seaports for out- and in-bound contacts with the Eastern Mediterranean. Finally, after the collapse of the Roman Empire, Apulia in general, though especially its northern part, endured a very complex succession of dominations, but to what extent these affected the composition of GS population is unclear (see Supplemental Text).

The coinheritance of genes and languages through generations has long inspired human population genetics research (Cavalli-Sforza 2001; Colonna et al. 2010; Longobardi et al. 2015; Lazaridis et al. 2025) and has been detected in a long list of cases. When applying this approach to the Greek-speaking communities of Southern Italy (another one is located in Southern Calabria), one could naturally posit that immigration movements brought together new genes and Greek as a new language in a linguistic landscape characterized by other idioms or, later, by Latin/Romance. Linguists have long and rather inconclusively debated whether the Greek language in Salento was introduced by the ancient western colonization (Rohlfs 1924) or by Medieval immigrations (Morosi 1870; Parlangeli 1953). Recently, a third, more nuanced account was proposed by (Fanciullo 1996; Fanciullo 2001) and has since been developed. With reference to Salento, this view envisages a complex history of continuity, renewed diffusion, and prolonged Greek–Romance contact from Antiquity onward.

In such a framework, tracing the genetic heritage of present-day inhabitants could be the most straightforward method to discriminate among the hypotheses. The advent of genome-wide analyses of diversity and the possibility of genotyping a subject’s DNA at millions of variants, coupled with methods of analysis scaled to the corresponding amounts of data (Moorjani and Hellenthal 2023), have led to an explosion of studies aimed at reconstructing how the gene pools of current populations formed. The characterization of DNA recovered from archaeological contexts has further empowered these analyses, by providing time-stratified representations of the diversity and how it spatially spread from ancient source populations. In the genomic landscape of Italy, the contribution of early hunter-gatherer populations emerges, overlayered by genetic components traceable to Neolithic and Bronze Age expansions (Pagani et al. 2016; Mathieson et al. 2018; Aneli et al. 2021). A first component is traced back to the Anatolian Neolithic (Hofmanová et al. 2016; Kilinç et al. 2016; Lazaridis et al. 2016; Omrak et al. 2016). A component related to Caucasus hunter-gatherers is more relevant in Southern than Northern Italy and has been conveyed by post-Neolithic movements (Raveane et al. 2019). Also, a component related to steppe people has been detected, as part of a gradient spanning the entire European continent (Haak et al. 2015; Saupe et al. 2021). These three components contribute to the diversity gradient along the Italian peninsula (Fiorito et al. 2016; Raveane et al. 2019).

Due to its position, Apulia is a segment of the so-called “Mediterranean genetic continuum” (Sarno et al. 2014; Haber et al. 2017; Lazaridis et al. 2017; Sarno et al. 2017; Feldman et al. 2019), i.e. a nearly continuous cline of variation stretching longitudinally from the Middle East to Iberia. As the core of this had settled since Neolithic times (Lazaridis et al. 2022a), the superimposition of multiple migratory east-to-west movements continued throughout the Bronze and Iron Ages, antiquity and the CE (Antonio et al. 2019; Lazaridis et al. 2022b; Antonio et al. 2024; Ravasini et al. 2024; Ravasini et al. 2025), further smoothing the pre-existing discontinuities. These processes involved individuals, families and entire communities from Greek antiquity to the end of the Byzantine Empire and beyond. Thus, when considering Southern Italy, and GS in particular, as landing points, in most cases the incomers (if any) may have come from areas where different varieties of Greek have been spoken for at least two millennia. Hence, to understand the geographic origin(s) of these people, considering also a temporal dimension becomes essential.

In this context, the eastern side of Southern Italy has been the focus of genomic studies aimed at understanding the contacts with the Balkan side, leveraging precisely dated archaeological specimens (aDNA). A close similarity emerged between Southern Italians and peoples from the Eastern Peloponnese (Raveane et al. 2022), in some case higher than ancient Peloponnesians and extant Greeks, a pattern attributed to elevated connectivity during the Iron Age. On the other hand, Iron Age inputs seem to have weakly impacted the Daunians (Aneli et al. 2022), a population of North-Eastern Apulia (see Supplemental Text) for which an autochthonous origin remains the most parsimonious scenario, in light of their sharing of a Western Hunter-Gatherers (WHG) signature with pre-existing Italic groups.

With these concepts in mind, we aimed at answering the following research questions: 1) Are there components of the GS genome pool that make it distinguishable from the rest of Southern Italy? And subordinately: 2) Are these components geographically rooted outside Italy? 3) Is GS characterized more strongly than the rest of Southern Italy by a genetic contribution attributable to the First Greek Colonization of the 8^th^ century BCE? 4) Are additional contributions, potentially attributable to the Byzantine rule, detectable and relevant?

Based on uniparental markers (Menato et al. 2025), we previously found few clear instances of GS-specific lineages that distinguished this community from the current surrounding genetic landscape of Southern Italy, with most likely provenances as diverse as the Middle East and the Balkans. We concluded that this minor component detected in the GS gene pool is the result of past immigrations, with the contribution of both sexes to the build-up of the entire community.

Here we report on the autosomal diversity of 27 natives of GS (Table S1) as obtained from whole genome sequencing and contextualized with different datasets of whole genomes and high-density SNP arrays from extant populations, covering the North Mediterranean, from the Iberian Peninsula and Central Europe to the Caucasus and the Middle East. We also considered different sets of genetic markers, each capable of revealing different aspects of the genetic structuring, at different spatial and temporal scales. Additionally, we put the same genotyping data in the context of variation in aDNA data from temporal strata from the Paleolithic to the Middle Age.

By analyzing the overall features of the GS gene pool in comparison with other North Mediterranean populations, as well as the patterns of affinity among GS municipalities, we support the most recent hypothesis formulated on linguistic grounds, which then becomes quite compelling: a prolonged and reinforced resilience of the Greek language in GS, which persisted *in situ* amid non-Greek-speaking surrounding peoples despite their abundant genetic contributions.

## Results

### GS in the contemporary context

We used whole genome sequencing of 27 GS natives and 21 controls (Table S1). In order to represent the genomic landscape of the continental areas that may have contributed to the GS gene pool, we included our original data into dataset of 503,578 (500K dataset) strictly biallelic SNPs in non-repetitive DNA regions, LD-pruned, with MAF >= 0.03 and less than 2% missing genotypes among 799 samples grouped into 34 population_ID’s, from the Iberian Peninsula to the Caucasus and the Middle East (Table S2). We obtained such a dataset by combining data from different sources (Mallick et al. 2016; Gilly et al. 2018; Bergström et al. 2020; Kars et al. 2021; Byrska-Bishop et al. 2022), with the aim of having nearly even representation (50-100) of key populations across this geographic range and in particular Italians, Greeks and Turks. In many cases the same national population was represented by multiple ID’s (Table S2) or multiple samples within the same population_ID derived from different sources and were sequenced on different platforms. This enabled a powerful control against batch effects potentially generated at any step leading to the final individual genotype description.

Principal component analysis performed on the 500K dataset (Figs. 1A, S1) was able to highlight the distinctness of some of the population_ID’s in the space of PC1 (0.58% of variance) and PC2 (0.29%). The positioning of samples on the PC1 axis tightly followed their longitude (horizontally flipped), with Middle Easterners at one extreme and Iberians and GBR at the opposite one. Sardinians and the populations from the Caucasus mapped at opposite extremes on PC2. At the center of the plot the Greeks and Italians formed two partially overlapping compact clouds, with the Turkish shifted to the upper-left, towards the Caucasus samples. A gradient within Italy could be captured, with samples from Bergamo (Northern Italy) mapping towards the Iberians and French, and the Tuscans in the middle of the Italian cluster. One Italian sample, sequenced in the frame of the MinE project (all samples lacking precise provenance) mapped within the Sardinian cluster, further confirming the fact that cluster affiliation as inferred by the position in the PC’s space, was not obscured or confounded by the particular sequencing platform. We note here that tight correlation between geography and PC’s, prominent here, is not an invariant feature across studies of the same area, an issue discussed in [(Lazaridis et al. 2014), Fig. S10.5].

**Fig. 1.**
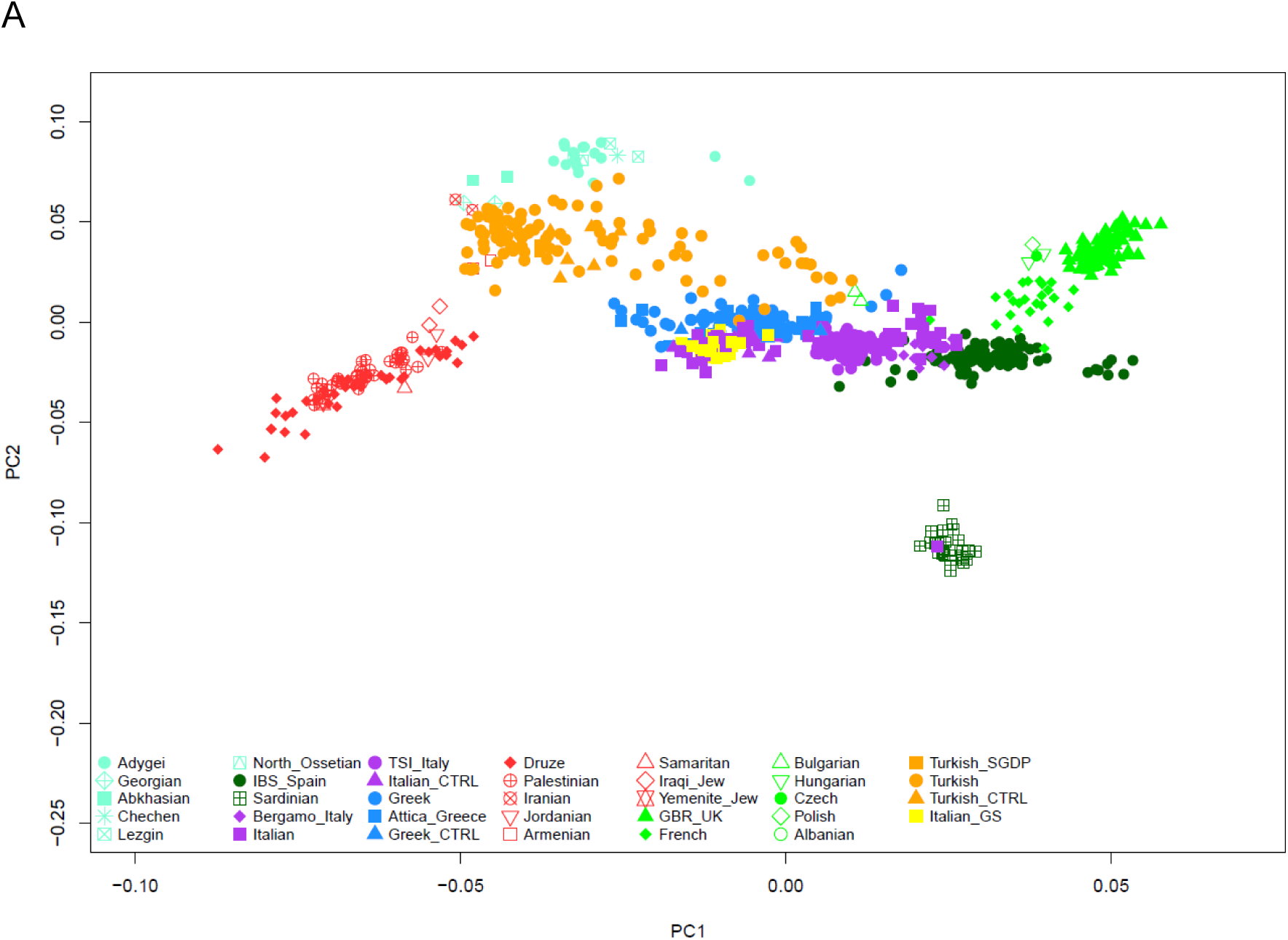
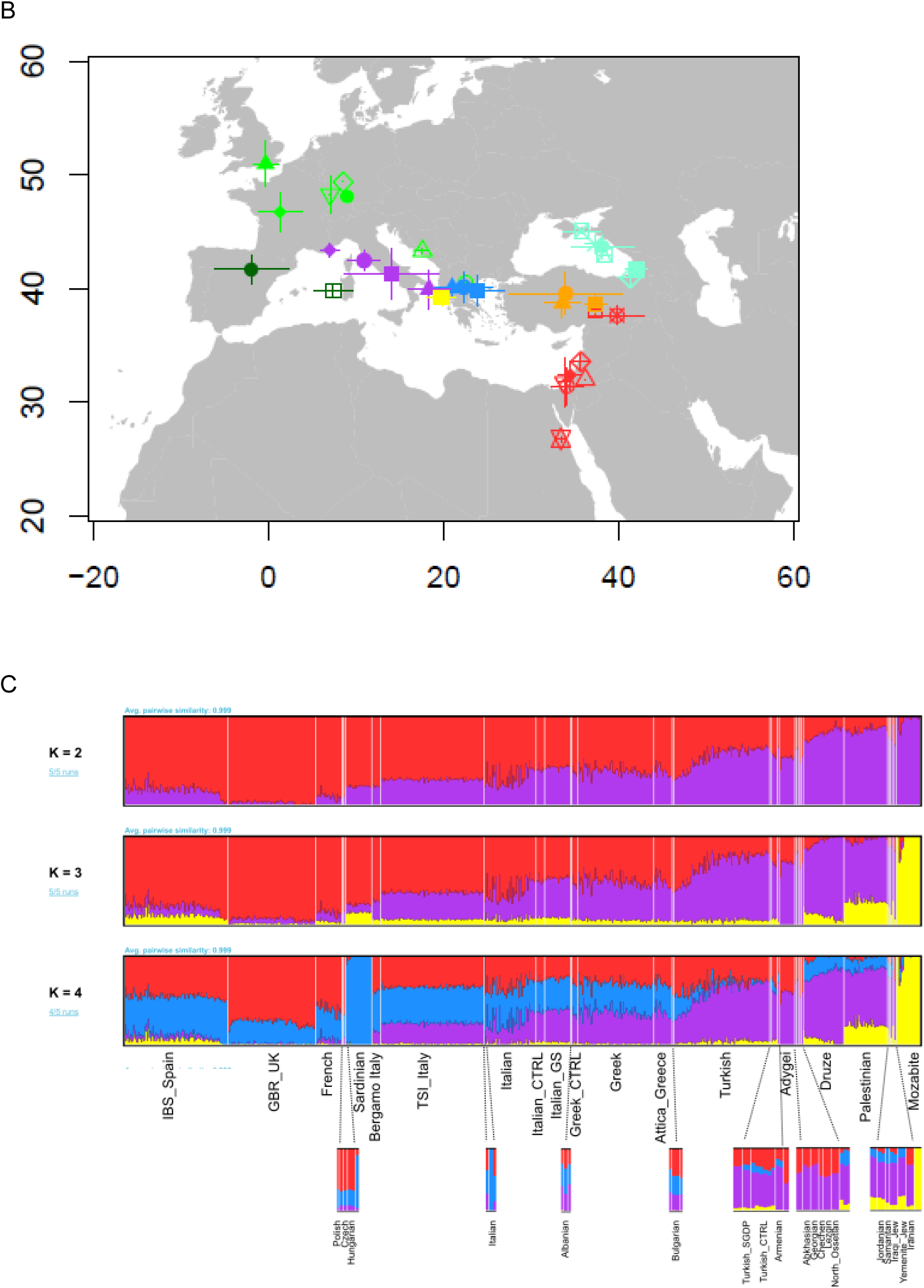
A) Plot of the 799 samples of the 500K dataset in the space of PC1 vs. PC2. The plots of PC1 vs the following PCs are reported as supplemental figures in a separate file. B) Plot of the 34 population_ID’s of the 500K dataset on their average “Expected Sampling Location” (see Supplemental Text for definition). Symbols and colours are as in the original PCA plot. Whiskers represent +/- 1 s.d. C) Admixture analysis of the 500K dataset for K 2 to 4 (complete figure in Supplemental Text, Fig. S4). Each plot is based on 5 runs, summarized with pong. The number of runs producing coherent partitions in K components is reported in light blue to the left. Population_ID for each vertical belt at K=4 are resolved at bottom, with small ID’s magnified. Also a section of the Italians is magnified, to show the single sample with a “Sardinian” pattern (see PCA).

The GS samples mapped to the left extreme of the Italian cluster, together with our Southern Italian controls, and slightly below the Greek cluster, showing that PC2 could potentially summarize the diversity across the two sides of the Adriatic Sea.

We then considered other PC’s, down to the 20^th^ (all significant at p<5E-50, Tracy-Widom statistics), obtaining the plots reported as supplemental figures. In all plots, samples from the same population mapped coherently, irrespective of their source (sequencing platform and variant calling), indicating that at this level of data filtering the 500K dataset was devoid of relevant batch effects. PC3 neatly separated the Druze from the rest of Middle Eastern samples, possibly highlighting a segment of a well-documented Eastern cline (Lazaridis et al. 2014). In some plots there were areas sparsely populated by samples, indicating lines of discontinuity of variation. For example, when focusing on PC2 and PC6 (Fig. S2) the group of GS samples was shifted from the cluster of TSI, Italians, and Southern Italian controls, sharing low PC6 values with some of the Greek samples and the single Albanian.

We then wanted to extract the maximum of geographic information from genomic data, by considering that, beyond PC1, other PC’s may convey information on latitudinal and/or longitudinal clines. The expected sampling locations conditional on the overall composition of the dataset (Figs. 1B, S3) revealed a remarkable predictive power, with only few, less numerous, population_ID’s mapping outside their true position (e.g. Bergamo_Italy, Armenia). In this plot, the GS group mapped right at the tip of Apulia, with a very light shift toward Greece.

In the unsupervised admixture analysis performed on the same data (Figs. 1C, S4), GS displayed patterns very similar to our Southern Italian controls. At K=3, a component (violet) which increased along the west-to-east axis and peaked among Druze and Palestinians was represented in GS (45%) more than the other Italians ID’s (20.9, 30.8, 33.0 and 41.9% for Bergamo, TSI, Italians and Italian_CTRL, respectively), resembling the Greek samples (40.8, 44.2 and 44.5% for Greek_CTRL, Greeks and Attica_Greece, respectively). Also, a minor component related to the Mozabites (yellow) and detected in the analyses by [(Raveane et al. 2019), Fig. S2C] and [(D’Atanasio et al. 2023), Fig. S1] appeared in GS. At K=4 the violet component characterized more clearly Italy (14.0, 23.4, 25.5 and 34.2% for Bergamo, TSI, Italians and Italian_CTRL, respectively) and Population_IDs further East, with the GS average (36.9%) shifted toward the Greek value (33.1, 36.2 and 36.3% for Greek_CTRL, Greeks and Attica_Greece, respectively).

Though accounting for few percents, the yellow North African/Levantine component was overrepresented in GS (3.2%) and Italian_CTRL (3.6%) than the other Italians [1.1, 1.2, and 2.3% for Bergamo, TSI and Italians, respectively, in agreement with (Moorjani et al. 2011)] and Greek ID’s (1.9, 1.2 and 0.6% for Greek_CTRL, Greeks and Attica_Greece, respectively) and was shared with the Iberians (5.8%) and part of Turkish samples (3.6, 4.1 and 6.0% for Turkish, Turkish_SGDP and Turkish_CTRL, respectively) [see also Fig. S7 in (Kars et al. 2021)].

These results show that information in the GS genomes can lead to the detection of subtle differences as compared to the rest of Italians represented in the 500K dataset.

When examining ancient admixture (Figs. 2, S12) with the f3 statistics (Patterson et al. 2006) using a distant outgroup in the forms f3(GS;A,Mbuti) and f3(Southern Italian controls;A,Mbuti) we obtained a superimposable order of populations, with Sardinians producing the lowest and Middle Eastern populations the highest values, respectively, in both cases. With D(GS, Southern Italian controls; A, Mbuti), we obtained a barely significant (p<0.05) negative value for the single Czech sample. Significantly positive values were obtained only for Greek_CTRL and Turkish_CTRL controls, indicating a closer affiliation of GS to these groups than Southern Italians. Interestingly, the top positive (though not significant) values were produced by Middle Eastern populations, suggesting that the similarity of GS to these latter tends to exceed that of Southern Italians. These results indicate that the GS pool fulfils the characteristics of the Mediterranean continuum (Raveane et al. 2019), but most likely with a stronger Middle Eastern than Caucasus contribution.

**Fig. 2a.**
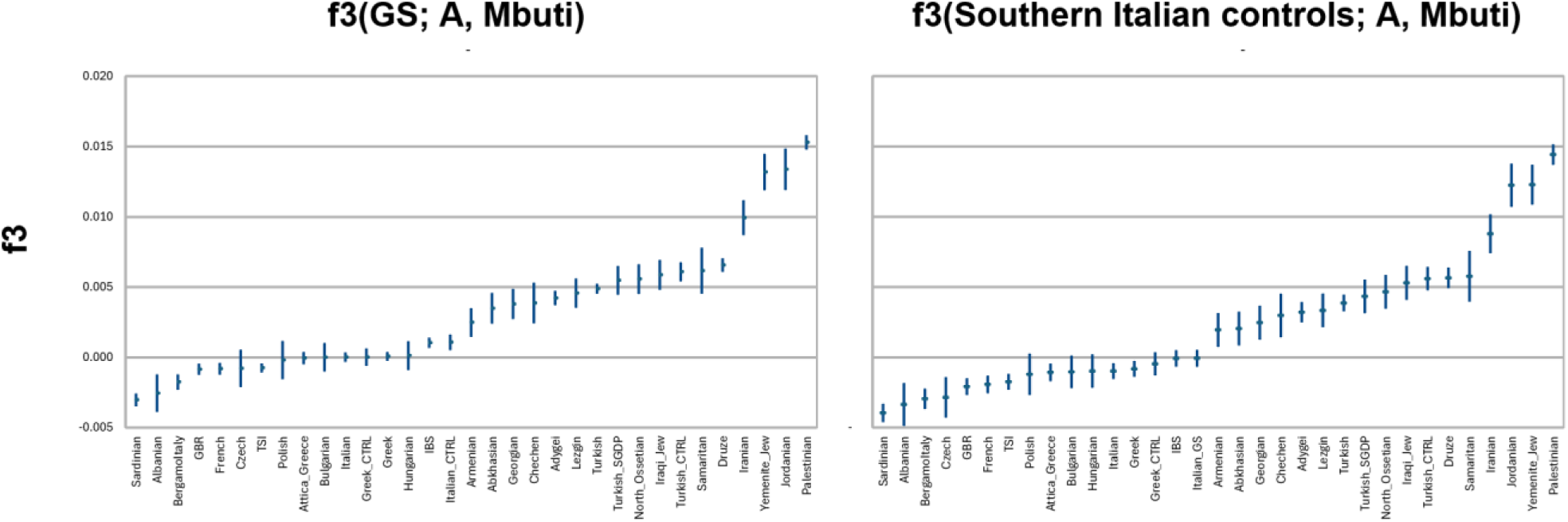
f3 statistics for GS (left) and Southern Italian controls (right) as targets for admixture between the 33 indicated Population_ID’s and the Mbuti as distant outgroup. For both GS and the Southern Italian controls the order of populations producing f3 values from smallest to largest and the absolute f3 values are highly similar, indicating no gross differences between GS and Southern Italians in their ancestry relationships with other populations. Whiskers indicate +/- 1.96 s.e.

**Fig. 2b.**
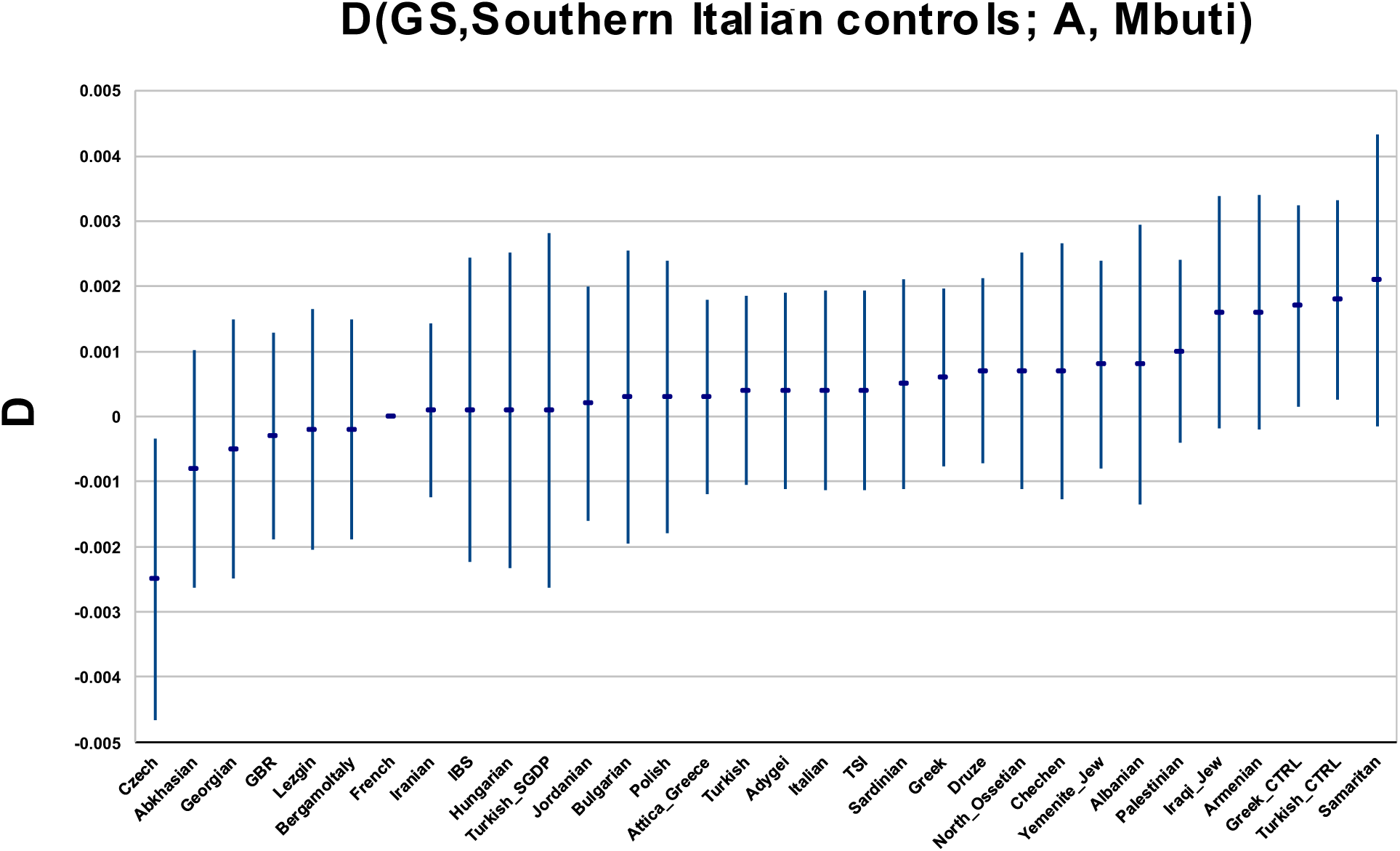
D statistics in the form reported above for 32 Population_ID’s, to test for the excess shared ancestry between GS and the indicated populations as compared to Southern Italian controls. Note the two positive significant values for Greek and Turkish controls. Whiskers indicate +/- 1.96 s.e.

We then wanted to verify the above suggestions on another set of variants (1K dataset), previously described as strong Ancestry Informative Markers, specifically for the European populations (Drineas et al. 2010), and nearly entirely independent of the 500K dataset.

Despite the dramatic reduction in the number of markers, the plot of PC1 and PC2 (1.67% and 0.6% of variance explained, respectively) obtained in this way replicated that of the 500K dataset, with PC1 displaying a longitudinal gradient and PC2 separating Sardinians and Caucasus populations at opposite sides (Fig. S5). However, all population clusters were less clearly separated, a quite expected result given that the original extraction of this set of markers did not consider Middle Eastern and Caucasus populations. The mapping positions of the GS samples were shifted towards Middle Eastern samples, at the (eastern) edge of the Italian cluster. When applying the expected sampling location procedure (Figs. 3, S6), the fidelity of population assignments was lower than in the 500K dataset, but the confidence intervals for each population_ID were still compatible with their original location, except for a shift towards the centroid for the Caucasus populations. Interestingly, GS underwent a gross repositioning, that brought it into the Aegean area, with confidence intervals that no longer reach the true geographical sampling position.

**Fig. 3.**
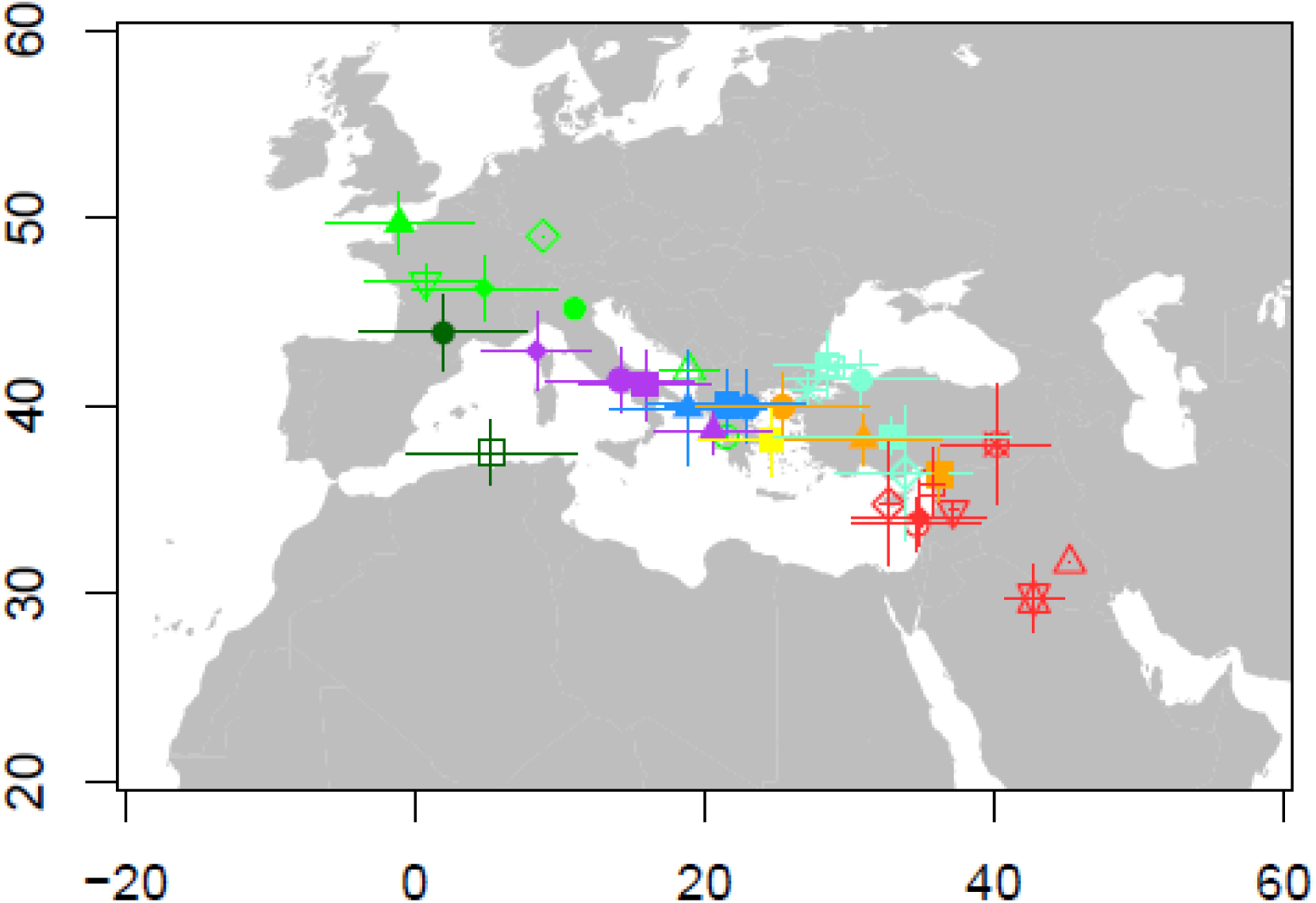
Plot of the 34 population_ID’s of the 1K dataset on their “Expected Sampling Location” (see Supplemental Text for definition). Symbols and colours are as in the previous PCA plot. Whiskers represent +/- 1 s.d

The above results suggested that a relevant part of GS ancestry could be spatially rooted in areas peopled by the Greeks. This does not necessarily imply that contributions to GS occurred in the period of the First Great Colonization (started in the 8th century BCE) which led to numerous settlements in the whole of Southern Italy and the first introduction of the Greek language there. A non-mutually exclusive possibility is that GS shares a little proportion of its ancestry with more easterly populations, whose contributions caused a partial rightward shift on the map and these, too, may have occurred over a long time span.

We then wanted to precisely put our GS samples in the context of subpopulations of the Balkan peninsula, Sicily, Crete, Aegean area and Cappadocia. With a panel of 25,645 SNPs genotyped in our samples and the samples studied by (Paschou et al. 2014), we obtained the PCA plot shown in Figs. 4, S7. The GS samples departed clearly from all subpopulations of Southern Balkan peninsula outside Greece as well as from the Italians [TSI and 13 HGDP Italians described in Supplemental Table 2 of (Paschou et al. 2014)]. The GS samples overlapped mostly with samples from Sicily and South-Eastern Peloponnese (Laconia), towards Crete and the Dodecanese (islands of the South-Eastern Aegean Sea).

**Fig. 4.**
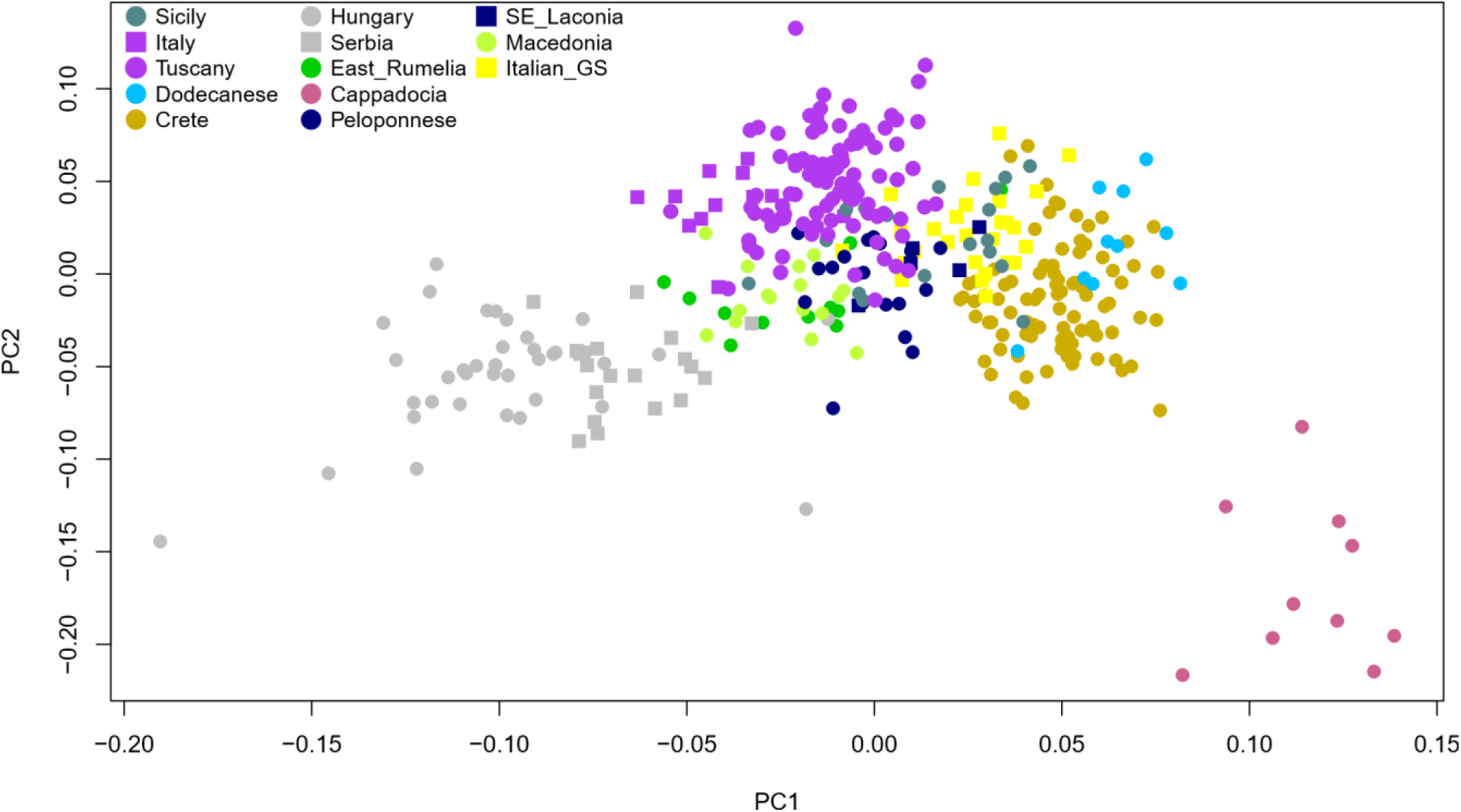
Plot in the space of PC1 and PC2 of 344 subjects from the Balkan peninsula, Italy, Crete and Anatolia typed by (Paschou et al. 2014), plus the 27 GS subjects

We searched for possible sources of recent affinity by analyzing patterns of sharing of 147,012 high confidence (Fig. S8) fragments identical by descent (IBD). The general pattern of sharing among samples (Figs. S9, S10) of the 34 population_ID’s closely replicated previous findings (Ralph and Coop 2013). In particular, sharing within populations was generally higher than between populations, with sharing among GS samples stronger than with any other ID. Also, a strongly significant decay with geographical distance was confirmed. We partitioned the complete list of fragments to replicate the analysis by (Ralph and Coop 2013); these authors estimated that only a small proportion of blocks longer than 2 cM are inherited from longer ago than 4,000 years, whereas for the most part, blocks longer than 4 cM come from 500–1,500 years ago. For the 3-5 cM range we found that GS shared almost equally with Italy and Greece, in the context of a generalized stronger sharing between European ID’s. While the same pattern was reproduced (to a lower magnitude) also for blocks longer than 5 cM, here GS displayed some degree of sharing with populations of Middle Eastern ancestry, a feature not replicated by Southern Italian controls or TSI. When plotted directly on maps (Fig. S11) the sharing of 3-5 cM blocks of all Italians was comparable and stronger towards Europe, but for blocks longer than 5 cM GS displayed a larger repertoire of possible connections. This was true also when GS was compared with Greek and Turkish ID’s.

In order to get an additional temporal dissection of the genomic contributions to GS, we considered variants reported by (Albers and McVean 2020), who obtained estimates for the date of appearance of derived alleles from the 1,000 Genomes data. We focused on markers with an estimated age of 300 generations or less which, assuming approximately 30 years/generation (Fenner 2005; Moorjani et al. 2016), are potentially able to report on contributions from the Neolithic onwards. Allele frequencies of these variants in the 1,000 Genomes EUR are almost invariably below 1% and in most cases below 0.5% (see Fig. 4 in the cited reference), and this was recapitulated in the frequency spectra of the 11,589 derived alleles among the 48 samples sequenced in this work (Fig. S14). Thus, conservatively, we limited this analysis to these 48 samples, to ensure that such rare variants be ascertained uniformly across samples The equal number of samples from each potential donor population and no evidence for an enrichment in rare alleles in the Turkish population, as previously reported (Kars et al. 2021), allowed direct comparisons for the degree of sharing (Fig. 5, S13).

**Fig. 5.**
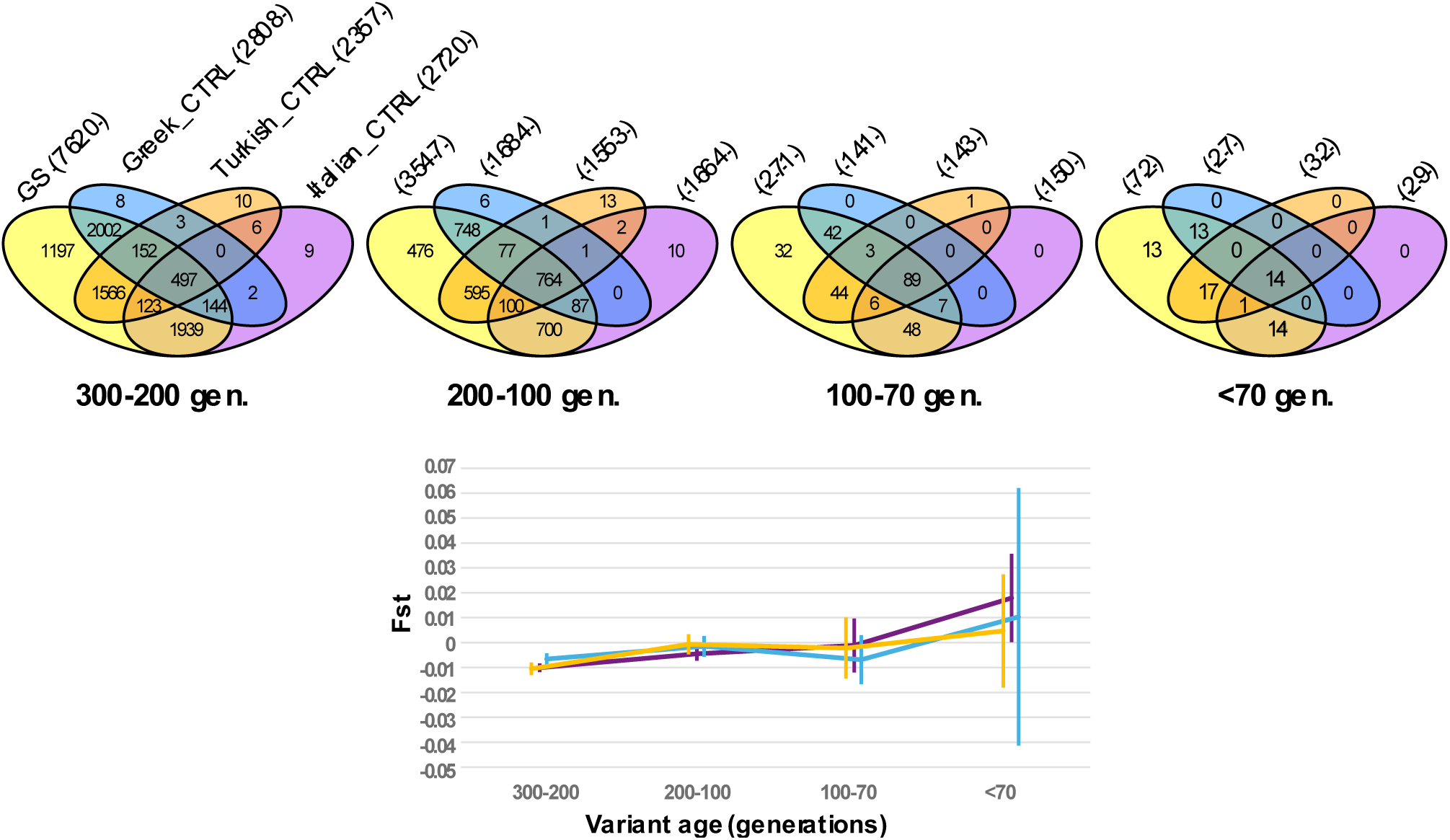
Top: Sharing of derived alleles at positions with estimated age in the indicated intervals. The Italian samples were downsampled to 6, so that the Venn diagrams are based on 27, 6, 6 and 6 subjects for GS (yellow), Greek (blue), Turkish (orange) and Italian (purple) controls, respectively, and occurrences within cells are thus directly comparable. The total number of derived alleles with non-null frequency in each group are reported in parentheses. Bottom: Weir and Cockeram’s *F_ST_* and 95% c.i. for comparisons between GS and Italian (not downsampled), Greek and Turkish controls. Colors as above.

For markers with ages in the range of 300-200 generations, i.e. in the Neolithic up to the onset of Bronze Age, GS samples shared a nearly equal number of derived alleles with Greeks and Southern Italians, and higher than with the Turkish controls. This pattern denotes a Neolithic structuring with stronger gene flow within Southern Italy and between it and Greece (Mathieson et al. 2018). A similar situation is observed for markers in the range 200-100 generations, but it is reversed for markers aged 100-70 or less than 70 generations. In these latter cases the number of markers shared between GS and Anatolia equals or even surpasses those with Greece and Southern Italy. Note that this analysis captures the surge in connectivity that occurred during the same periods (Antonio et al. 2019; Lazaridis et al. 2022a; Lazaridis et al. 2022b; Antonio et al. 2024). In fact, in the 300-200 generations range, the number of markers shared by all sample groups (central cell in the diagrams) is definitely lower than those shared between GS and each of the other three groups. Conversely, in the 200-100 and 100-70 generation ranges it is higher and, at 70 generations becomes comparable.

When measured with *F_ST_*, differentiation between GS and Turkish_CTRL controls remained null throughout the timepoints, while it increased with Italian controls to approach significance for markers younger than 70 generations.

### GS in the ancient context

In order to obtain clearer insights on the tempo of formation of the GS pool, we analysed our samples in the context of 1,275 modern and 4,021 ancient samples (Mallick et al. 2024; Mallick and Reich 2024) covering the Paleolithic to the Middle Age. In the PCA (Figs. 6, S15), the GS group clustered between the Southern European and Middle Eastern populations, closely aligning with modern Southern Italian individuals. Ancient Greek individuals exhibited a temporal gradient along the vertical axis. By contrast, Byzantine genomes were arranged along a predominantly horizontal cline corresponding to a west-to-east geographic distribution (left to right), with individuals from Western Anatolia clustering closest to the GS group.

**Fig. 6.**
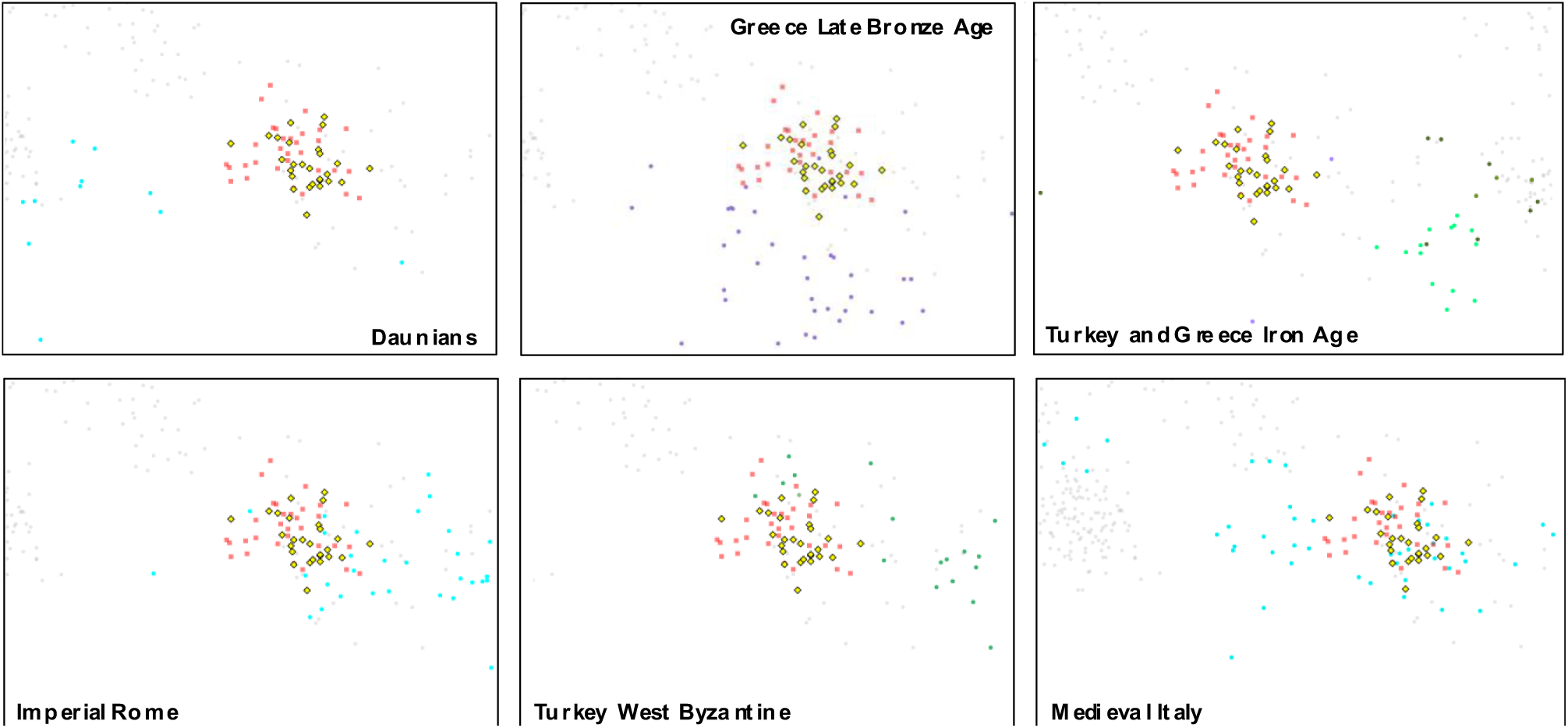
Close-up of the central part of PCA plot of Fig. S15, showing the positioning of the Daunians with respect to the Southern Italian (pink squares)/GS (yellow lozenges) cluster. In the remaining panels the other sample groups that overlap the same cluster. In the top right panel Iron Age Turkish subgroups (shades of green) and Greeks (violet) are shown together.

A focus on the GS group revealed no overlap with the Iron Age Daunians, in agreement with their distinctiveness from modern-day Apulians and Southern Italians in general (Aneli et al. 2022). Similarly, no overlap with other Iron Age Italian groups could be observed. The Greek Late Bronze Age group showed the highest affinity to GS but no overlap, indicating additional components to explain the current composition of the latter. The first ancient Italian samples showing some overlap with GS are those from the Imperial period, i.e. a period that postdates the Latinization of Apulia, for which an ancestry shift towards the Eastern Mediterranean has been reported (Antonio et al. 2019). Interestingly, it has been suggested that the eastern ancestry shift observed in the Italian Imperial gene pool actually started with the First Great Colonization of Southern Italy (Ravasini et al. 2025). However, Greek and Anatolian Iron Age samples do not overlap GS, either. The fall of the Western Roman Empire led to a new ancestry shift in the Italian peninsula towards continental Europe: these samples form an elongated cline from east (in continuity with the Imperial period) to west (in relation to Germanic movements) (Speidel et al. 2025), representing the ultimate attractors for the modern Italian (and GS) gene pool.

In conclusion, our analysis seems to point to a possible Greek influx in the first formation of the GS (and Southern Italian) genetic variation, followed by other major contributions from further north along the Italian peninsula and from the East. On the other hand, among the samples of the Byzantine period, only groups from present-day Western Anatolia partly overlap with GS, possibly due to internal structuring already revealed among Neolithic samples from the Anatolian peninsula (Mathieson et al. 2018), and that may have persisted through the Byzantine period till today [Fig. S3 by (Kars et al. 2021)].

### Genomics within GS

The question of whether the GS community behaved as a closed endogamous population or was open to external contributions can be tackled on genomic grounds. First, we replicated the IBD analysis by partitioning each pair of carriers of IBD’s according to the municipality of origin (Fig. S16). Interestingly, at this geographical scale the general pattern of stronger within than between sharing was reverted. In fact, the average number of fragments shared between any two GS samples is one order of magnitude higher than between GS and other ID’s samples (Fig. S10), but average sharing between GS subjects of the same municipality is generally lower than between subjects of different municipalities. This was further confirmed by the analysis of runs of homozygosity (ROH), i.e. blocks with identical strings of marker alleles on the two autosomes of the same subject. Our analysis revealed lower incidences of ROH’s in Europeans than Turkish (Kars et al. 2021) and Middle Easterners (Fig. S17), especially for longer blocks. In this general framework we found no evidence for increased occurrence of ROH’s in GS as compared to other Italians and in particular for longer ROH’s, which reflect population isolation and grandparental endogamy (McQuillan et al. 2008). For ROH’s longer than 1500 kb our F_roh_ was 0.003, a value consistent with the absence of common paternal and maternal ancestors in the last five generations (McQuillan et al. 2008), i.e. a period in which the use of Greek language was a more distinctive signature of GS than today. The above analyses represent the first genomic documentation of the wide connectivity in GS (Aprile 2015; Romano 2019). Also, the detected patterns of sharing do not reveal obvious relationships with the historical pattern of adherence to language or religious practices, which were reported to vary among municipalities (Magurano 2020).

**Fig. 7.**
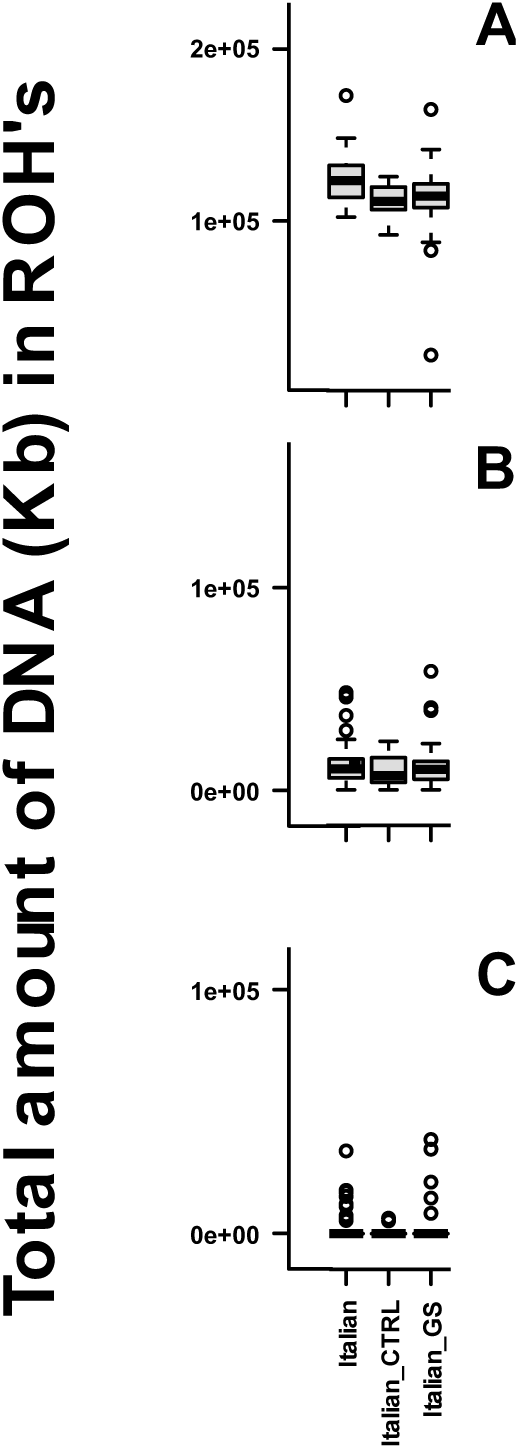
Close-up of the boxplot of amount of DNA (in Kb) represented in ROH’s longer than given thresholds in the autosomes of the 27 GS samples compared to Italian_CTRL and Italians (complete figure in Fig. S17). ROH’s longer than 500 Kb (A), 1,500 Kb (B) and 5,000 Kb (C).

## Discussion

The background of this work is the long-standing linguistic debate on the origins of the Greek language (Griko) spoken in the enclave of Grecìa Salentina. One is that this Greek dialect is a direct descendant of the Medieval Greek varieties introduced during Byzantine rule especially between the 9th and 11th centuries CE. The other view is that it derives from the Greek-speaking communities that settled in Southern Italy in the First Great Colonization (for a more thoroughly discussion see Supplemental Text). The third possibility is the more recent synthesis postulating a linguistic continuity since ancient times (cf. Introduction). As far as abundant population movements impacted Apulia and the rest of Southern Italy in all the relevant time windows, the idea of a demic-mediated transfer of language is more likely than a purely cultural transmission. We then wanted to test which of these hypotheses was best supported on genomic grounds. To this aim, we used whole genome sequencing of 27 natives of GS and 21 controls (Tables S1, S2), to take full advantage of an unbiased ascertainment of variants (Bergström et al. 2020). We paid particular attention to embedding the sequencing data of the above samples in a dataset in which the two main areas where GS ancestry could be putatively placed (i.e.the Greek and Anatolian peninsulas) were evenly represented, a feature that is seldom found in the genomic literature on the area. This required the merging of sequencing data from a variety of sources (Mallick et al. 2016; Gilly et al. 2018; Bergström et al. 2020; Kars et al. 2021; Byrska-Bishop et al. 2022) and a subsequent stringent filtering that rendered precisely comparable the samples from the same population(s) but different sources.

All analyses performed on the full dataset point to the same major conclusion, i.e. that the GS genomic endowment bears the same main features of the rest of Southern Italy (Fiorito et al. 2016; Raveane et al. 2019). This is characterized by a massive contribution of a Neolithic component that overlayered on the pre-existing background and is nowadays primarily evident in major Mediterranean islands (Fernandes et al. 2020; Marcus et al. 2020), followed by an additional entry of a component that reached the North-Central Mediterranean in the Bronze Age and was variably dubbed in the literature (Iranian Neolithic, Anatolian Bronze Age). Thus, we found no definite evidence for a single source population for GS outside Italy.

This observation alone, coupled with the analysis of internal connectivity, reaffirms that GS has been all but a genetically closed community, despite retaining the Greek language as a distinctive feature. In this regard GS is part of the consolidated “Mediterranean genetic continuum” (Sarno et al. 2014; Haber et al. 2017; Lazaridis et al. 2017; Sarno et al. 2017; Feldman et al. 2019).

As to a possible major legacy of the Great Colonization in GS, only some of our analyses indicate a slightly enhanced similarity of GS with modern and ancient Greece with respect to the rest of Italy, i.e. the D statistics and the PCA with the 1K dataset (Figs. 2,3). We note here that, given the ease of maritime contacts from the Neolithic onwards (Rowley-Conwy 2011; Paschou et al. 2014) a quantitative evaluation of the genetic contribution of the Great Greek colonization in Southern Italy is and will remain difficult to attain. A generalized similarity between Southern Italy and the Peloponnese was detected (Raveane et al. 2022), but Bronze Age Southern Greece already harboured the same main components mentioned above (Clemente et al. 2021), which may have moved well before and independently of the settlers of the colonies in Southern Italy. Also, our analysis (Fig. 6) shows that Anatolian and Greek Iron Age ancestries shaped only marginally Southern Italy and GS. Archaeological and genetic (Lazaridis et al. 2022b) evidence does not support the simplistic idea of a generalized inter-mixing between Greek colonists and local populations. This seems particularly unlikely for the Messapians, in view of their conflicting relationships with the powerful Greek colony of Taranto.

Our analysis of GS in the context of Balkan and Cappadocian samples by (Paschou et al. 2014) rules out a quantitatively major autosomal contribution from the Eastern Adriatic coast, as would be expected on the basis of some historical sources mentioning Illyria as a possible homeland of the Messapians (Supplemental Text) and on genomic analyses in the Picenes, (another Adriatic people from further north along the Italian peninsula) where a Balkan genetic influx was suggested (Ravasini et al. 2024). Instead, in our analysis, GS departs from other peninsular Italians and maps among Sicilians and Eastern Aegeans and Cretans (Drineas et al. 2019). People fleeing from Crete towards Sicily and Southern Italy are reported as legendary forerunners of the Messapians (Supplemental Text). However, as this plot (Fig. 4), too, is characterized by strong correlation with geography, it is possible that the observed shift could be the effect of a minor amount of influxes from further east.

Suggestions of a slightly increased eastern contributions to GS derive from three lines of results, i.e. the enhanced yellow component in ADMIXTURE analysis, the pattern of IBD sharing and the sharing of recent dated alleles (Figs. 1, 5, S11). All of these results point to recent events, possibly of the last two millennia. In particular, the first one recapitulates the geographic pattern of African admixture around the Northern Mediterranean detected by (Moorjani et al. 2011), who estimated a date of 62 generations ago for the entry in Southern Italy, i.e. in the Roman Imperial period. Given the high representation of this component in the Middle East, a secondary contact rather than direct African-European contacts should be considered.

The three results above align with the finding, in GS, of Y chromosomal lineages rooted in the Near East (Menato et al. 2025) and with the overlap with some Western Anatolia’s Byzantine samples (Fig. 6). It is possible that these represent a genuine remainder of a Byzantine contribution in modern GS, in the same period proposed for a major contribution in the Balkans (Olalde et al. 2023), but they may also be the result of the general “Easternization” that occurred from the end of the Republican period throughout the Western Roman Empire period (Antonio et al. 2019; Ravasini et al. 2025).

Before and after the schism of the Roman Empire, and despite the large variation of the Eastern Roman Empire dominions over centuries, Greek was a commonly spoken language across a huge territory, spanning from the Central Mediterranean to the Near East and Egypt.

This means that, beyond the settlers of the Greek colonies in Apulia and the rest of Southern Italy, many of the eastern incomers of the subsequent 20 centuries (Supplemental Text) were actually Greek speakers. By measuring genetic differentiation across different temporal transects in an area broadly coinciding with that covered by our dataset, (Moots et al. 2023) and (Antonio et al. 2024) have noticed a stable degree of population structure since the Bronze Age, despite a marked surge in individual dispersal, contradicting population genetic theory, which would predict a collapse of population structuring. The “transient mobility” model of these authors is based on “the hypothesis that in the historical period there was an increasing decoupling of movement and reproduction”. As far as GS is concerned, a possible addendum is that the marked linguistic and cultural novelties carried over by long distance migrants acted as barriers to inter-marriage though not to trade and other social interactions.

We consider this model particularly fit to interpret the linguistic vs genetic histories of GS, especially in light of some pulses (e.g. monks and religious refugees of late antiquity/Middle Age) of eastern immigration. In other words, as contrasted with its high connectivity with the short-distance surrounding population background, GS would have been genetically impacted only to a minor extent by long distance immigrants (supposedly from Greek-speaking areas).

The reason why Greek has persisted only in GS, as compared to the rest of the former Magna Graecia, may lie in the synergy of two processes: 1) its somewhat neglected position, in an area shifted away from both immediate coastal landing places, and affected by enduring territorial disputes between the Byzantine Empire and a long list of local rulers, and 2) the exposure to reiterated influxes of long-distance incomers from all the Greek-speaking Eastern Mediterranean (see Supplemental Text).

In other words, the results of the present work do not support either of the two mutually exclusive “Classical” vs “Byzantine” hypotheses. Instead, they constitute a genetic validation of the more recent linguistic hypothesis (Fanciullo 2001) of the persistence *in situ* of a language that was instrumental to the resident population and thus maintained and cultivated, but sometimes rejuvenated by additional later Greek inputs. Byzantine rule would thus represent a major phase in the consolidation of Greek in the area, rather than its point of origin.

## Methods

### Sampling

We used samples collected in two temporally distinct campaigns. The first one was performed in 1994 (samples 20-27 in Table S1). The use of these samples in genomic studies was approved by the Ethical Committee Fondazione IRCCS Policlinico San Matteo (protocol number 0028298/22). The second one began in 2019 and is still ongoing (samples 1-19 in Table S1). Research procedures and the form for informed consent were approved by the local Ethics Committee (Comitato etico ASL Lecce, verbali n. 34 4/7/2019; n. 35 del 25/7/2019; n. 41 14/1/2020). Eleven out of the 19 participants from this series reported knowledge and use of the Griko language.

This design was applied to control for gross shifts among localities and across generations. In both cases, written informed consent was obtained after the aims and scopes of the project were illustrated to the participants and their communities.

All the research was performed in accordance with the relevant guidelines and regulations reported in the abovementioned documents and those agreed upon by the scientific community. In this respect, because the present work did not involve any issue relevant for the donor’s health, only the relevant prescriptions of the WMA Declaration of Helsinki and COE Oviedo Convention were obeyed.

Biological samples (buccal swabs) were anonymized upon collection. DNA was prepared with standard methods. Exclusively based on DNA quality, eight (7M, 1F) and nineteen (13M, 6F) from the first and the second campaigns, respectively, were entered into the whole-genome sequencing pipeline.

The project received the endorsement of the Union of the Municipalities (https://www.unionegreciasalentina.le.it/).

Details on the sequencing pipeline, variant calling and other computational methods are provided in the Supplemental Text.

## Supporting information

PCA PLOTS 1-20

## Acknowledgements

We are grateful to all the donors for their enthusiastic participation in this investigation. The authors would like to thank the ‘Project MinE ALS Sequencing Consortium’.

## Authors’ contributions

Conceptualization of the work: AN, EDA, OS, VS

Wet DNA work: CJ, EM, FZ

Analysed the data: GC, MG, FM

Management of computational facilities and support: GP, MH-C

Provided reagents (samples): BT, FC, NA, OS, VS

Linguistic and historical supervision: CG, GL

Wrote the paper: AN, EDA

All authors read and approved the last version of the manuscript.

## Data availability

Genotypes of 27 GS and 21 control samples for the 500K and 1K dataset markers are being submitted to EGA (https://submission.ega-archive.org/).

## Ethical statement

To obey all the prescriptions of the EU General Data Protection Regulation, we planned appropriate safeguards for processing data under the research exemption. The samples have been anonymized. Only dedicated, restricted-access bioinformatic infrastructures were used. Electronic data management was supervised by an institutional technology director. The research complied with the DNSH principle of the E.U.

## Declaration of generative AI and AI-assisted technologies in the writing process

No generative AI or AI-assisted technologies were used in the preparation of this manuscript and the supplementary materials.

