## Supplemental text and Figs. S1-S17 for "LAYERED GENOMIC COMPOSITION AND LINGUISTIC CONTINUITY IN GRECÌA SALENTINA"

#### Disclaimer

Throughout the main text and this text we tried to adhere as much as possible to the use of the terms Italy/Italian and Greece/Greek in their geographic acceptation, i.e. to indicate the respective peninsulas, without reference to the past or present political entities/borders. As to the Turkish peninsula (present day Türkiye) we used the term Anatolia in its broad definition, i.e. to include regions to the East of the Euphrates River. The use of this nomenclature was dictated by the fact that the time periods addressed in this work predate the arrival of Turkic peoples in the area. For sample and sample groups nomenclature we used the term Turkish, as they refer to present-day subjects.

#### Geographic, population and linguistic context

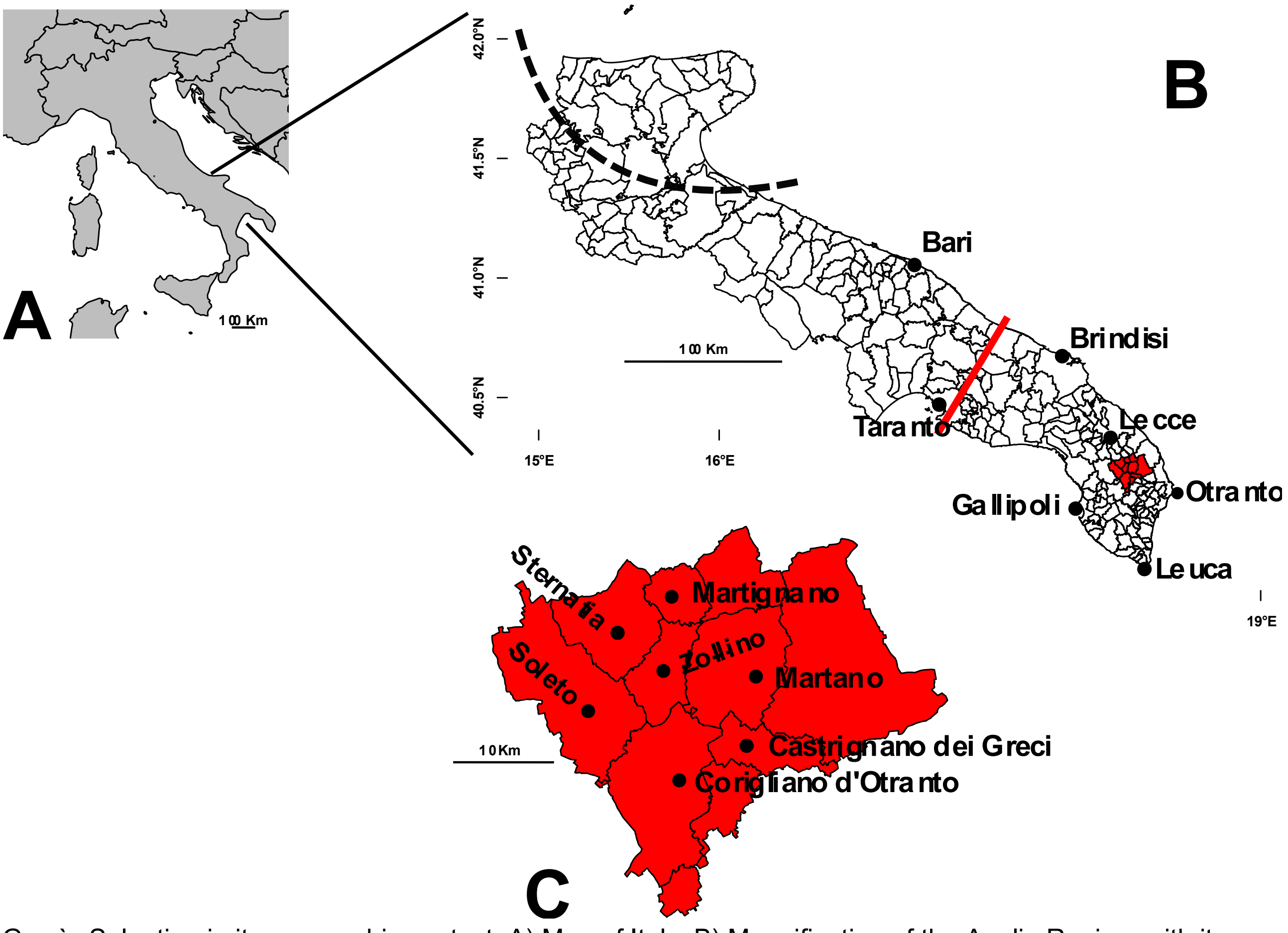


Grecìa Salentina in its geographic context. A) Map of Italy; B) Magnification of the Apulia Region, with its municipalities and GS highlighted in red. Cities mentioned in this text are indicated. The red line indicates the boundary of Salento. The dashed line indicates the approximate boundaries of ancient Daunia (Aneli et al. 2022); C) Magnification of the 10 Greek-speaking municipalities of GS. The 7 main centres of provenance of the 27 GS samples who underwent WGS are shown.

Linguistics (GL, CG)

The origins of the modern Greek-speaking enclaves in Southern Italy (SI) are a controversial, and essentially unsolved, issue in linguistics. In Italy, Greek-speaking communities are found in Salento (Grecìa Salentina, henceforth GS) and Southern Calabria (Bovesìa). These two Greek-speaking enclaves are the relics of a historically much more widespread community, whose presence in Southern Italy has been massive and uniform at least since the Great Greek Western Colonization (8th-5th century BC): the whole area (roughly including large portions of the regions of Sicily, Calabria, Basilicata, and Apulia) was Greek-speaking before the spread of Latin (Rohlfs 1972; Fanciullo 2001). This Greek linguistic substratum, along with the pre-Roman Italic one, has influenced in various respects the local Romance dialects which derive from Latin [(Ledgeway 2013) and literature cited].

The Greek dialect spoken in GS (now less widespread than in the past) is usually labelled Griko. One hypothesis is that this dialect is a direct descendant of the Medieval Greek varieties introduced in SI during Byzantine rule (Morosi 1870; Parlangeli 1953), especially between the 9th and 11th centuries CE, when waves of Greek-speaking settlers, soldiers, monks, and administrators are believed to have migrated to the region. According to this view, this dialect is essentially a “Modern” Greek dialect, that originates from the same linguistic source as most present-day Greek varieties (i.e., the so-called Hellenistic Koiné). Another hypothesis is that it directly derives from the Greek-speaking communities that settled in SI during the First Great Colonization beginning in the 8th century BCE (Rohlfs 1924). Linguistically, this would imply that Griko originates from ancient Greek varieties that were spoken before the “linguistic unification” brought about in Hellenistic times through the Koiné. Although the debate in the literature has been intense, no unambiguous evidence has thus far emerged to decisively support either hypothesis. Recently, a third, more nuanced account was proposed by (Fanciullo 1996; Fanciullo 2001) and has since been developed in subsequent work (Ralli 2006; Squillaci 2017; Ledgeway et al. 2018). With reference to Salento, this view envisages a complex history of continuity, renewed diffusion, and prolonged Greek–Romance contact from Antiquity onward.

The Greek identity of Griko is etymologically obvious for most of the vocabulary, but can even more precisely be assessed through quantitative tools applied also to other linguistic levels including grammatical rules (Guardiano et al. 2016; Guardiano et al. 2021).

Indeed, the full picture of the linguistic background of SI is complex. On the upper bound of the chronology, the emergence of Greek, Albanian and other supposed Paleo-Balkan languages can well be considered a consequence of the entry of Yamnaya people into the Balkan Peninsula 5-4.5 Kya (Lazaridis et al. 2022). Only after a few millennia was historical Greek documented and then expanded well beyond the Balkan peninsula.

As for the western side of the Adriatic, we know that in Southern Italy (SI: roughly including large portions of Sicily, Calabria, Basilicata, and Apulia, comprehensively known as Magna Graecia) the Greek language was introduced as a superstrate along with the First Colonization (since the 8th century BCE), i.e. much later. From then on, it spread across the area, where it was regularly spoken, along with other local languages: in Salento the two most relevant ones were probably Oscan and Messapic (the latter presumably related to Illyrian and thus also coming from the Balkans). Then came the superimposition of Latin, starting from the 2nd-1st century BCE (Rohlfs 1924). However, the latter was not abrupt: Latin and Greek varieties lived together for centuries and influenced each other with various outcomes. For example, the development of several

Romance dialects in the area has been heavily influenced by contact with Greek (Ledgeway 2013) and, viceversa, the current Greek varieties of SI have been influenced by contact with Romance, particularly in recent times, possibly with regional and temporal differences in the process of merging as suggested at least by syntactic features (Guardiano and Stavrou 2019; Guardiano and Stavrou 2020; Guardiano and Stavrou 2021). This long-lasting mixing has produced a network of layers of linguistic traits that cannot be easily disentangled. The consequence is that linguistic analysis cannot produce any uncontroversial timing for the origins of the Greek dialects now spoken in SI.

Sociolinguistically, Greek in SI is in regression/obsolescence: "true" native speakers have almost disappeared (these varieties are no longer acquired as first languages), no Greek speaker is monolingual (most are native speakers of Italian, and often of a Romance dialect), and Romance varieties are preferred for everyday interaction, because they cover a greater range of communicative needs (Sobrero and Romanello 1977; Gruppo di Lecce 1980; Miglietta and Sobrero 2006; Miglietta and Sobrero 2007; Romano and Marra 2008). The language has generally low social prestige: Greek-speaking communities are small and traditionally located in rural, poor areas. In Salento, there are traces of higher "vitality" and "resistance": as noted (Manolessou 2005), «in contrast to Calabria, the environment is an ally and not an enemy of the Greek language: Salento is a fertile plain, currently experiencing a period of economic and touristic development, something which has repercussions on the prestige of the Greek dialect». By contrast, in Calabria, Greek is nowadays "practically extinguished" (Martino 2009).


Aneli S, Saupe T, Montinaro F, Solnik A, Molinaro L, Scaggion C, Carrara N, Raveane A, Kivisild T, Metspalu M, Scheib CL, Pagani L (2022) The genetic origin of Daunians and the Pan-Mediterranean Southern Italian Iron Age context. Mol Biol Evol 39

Fanciullo F (1996) Fra Oriente e Occidente. Per una storia linguistica dell'Italia meridionale. ETS, Pisa, Italy

Fanciullo F (2001) On the origins of Modern Greek in Southern Italy. First international conference of Modern Greek dialects and linguistic theory: 67-77

Gruppo di Lecce (1980) Il caso Grecìa. In: Albano Leoni F (ed) I dialetti e le lingue delle minoranze di fronte all'italiano. Bulzoni, Roma, pp 343-403.

Guardiano C, Longobardi G, Cordoni G, Crisma P (2021) Formal syntax as a phylogenetic method. In: Janda RD, Joseph BD, Vance BS (eds) Handbook of Historical Linguistics II. Wiley Blackwell, London

Guardiano C, Michelioudakis D, Ceolin A, Irimia MA, Longobardi G, Radkevich N, Silvestri G, Sitaridou I (2016) South by Southeast. A syntactic approach to Greek and Romance microvariation. L'Italia dialettale 77: 96-166

Guardiano C, Stavrou M (2019) Adjective-noun combination in Romance and Greek of Southern Italy. Polydefiniteness revisited. J Greek Linguist 19: 3-57

Guardiano C, Stavrou M (2020) Dialect syntax between persistence and change. The case of Greek demonstratives. L'Italia dialettale 81: 121-158

Guardiano C, Stavrou M (2021) Modeling syntactic change under contact: the case of Italiot Greek. Languages 6: 74

Lazaridis I, Alpaslan-Roodenberg S, Acar A, Açıkkol A, Agelarakis A, Aghikyan L, Akyüz U, Andreeva D, Andrijašević G, Antonović D, Armit I, Atmaca A, Avetisyan P, Aytek A, Bacvarov K, Badalyan R, Bakardzhiev S, Balen J, Bejko L, Bernardos R, Bertsatos A, Biber H, Bilir A, Bodružić M, Bonogofsky M, Bonsall C, Borić D, Borovinić N, Bravo Morante G, Buttinger K, Callan K, Candilio F, Carić M, Cheronet O, Chohadzhiev S, Chovalopoulou ME, Chryssoulaki S, Ciobanu I, Čondić N, Constantinescu M, Cristiani E, Culleton BJ, Curtis E, Davis J, Demcenco TI, Dergachev V, Derin Z, Deskaj S, Devejyan S, Djordjević V, Duffett Carlson KS, Eccles LR, Elenski N, Engin A, Erdoğan N, Erir-Pazarcı S, Fernandes DM, Ferry M, Freilich S, Frînculeasa A, Galaty ML, Gamarra B,

Gasparyan B, Gaydarska B, Genç E, Gültekin T, Gündüz S, Hajdu T, Heyd V, Hobosyan S, Hovhannisyan N, Iliev I, Iliev L, Iliev S, İvgin İ, Janković I, Jovanova L, Karkanas P, Kavaz-Kındığılı B, Kaya EH, Keating D, Kennett DJ, Deniz Kesici S, Khudaverdyan A, Kiss K, Kılıç S, Klostermann P, Kostak Boca Negra Valdes S, Kovačević S, Krenz-Niedbała M, Krznarić Škrivanko M, Kurti R, Kuzman P, Lawson AM, Lazar C, Leshtakov K, Levy TE, Liritzis I, Lorentz KO, Łukasik S, Mah M, Mallick S, Mandl K, Martirosyan-Olshansky K, Matthews R, Matthews W, McSweeney K, Melikyan V, Micco A, Michel M, Milašinović L, Mittnik A, Monge JM, Nekhrizov G, Nicholls R, Nikitin AG, Nikolov V, Novak M, Olalde I, Oppenheimer J, Osterholtz A, Özdemir C, Özdoğan KT, Öztürk N, Papadimitriou N, Papakonstantinou N, Papathanasiou A, Paraman L, Paskary EG, Patterson N, Petrakiev I, Petrosyan L, Petrova V, Philippa-Touchais A, Piliposyan A, Pocuca Kuzman N, Potrebica H, Preda-Bălănică B, Premužić Z, Price TD, Qiu L, Radović S, Raeuf Aziz K, Rajić Šikanjić P, Rasheed Raheem K, Razumov S, Richardson A, Roodenberg J, Ruka R, Russeva V, Şahin M, Şarbak A, Savaş E, Schattke C, Schepartz L, Selçuk T, Sevim-Erol A, Shamoon-Pour M, Shephard HM, Sideris A, Simalcsik A, Simonyan H, Sinika V, Sirak K, Sirbu G, Šlaus M, Soficaru A, Söğüt B, Sołtysiak A, Sönmez-Sözer Ç, Stathi M, Steskal M, Stewardson K, Stocker S, Suata-Alpaslan F, Suvorov A, Szécsényi-Nagy A, Szeniczey T, Telnov N, Temov S, Todorova N, Tota U, Touchais G, Triantaphyllou S, Türker A, Ugarković M, Valchev T, Veljanovska F, Videvski Z, Virag C, Wagner A, Walsh S, Włodarczak P, Workman JN, Yardumian A, Yarovoy E, Yavuz AY, Yılmaz H, Zalzala F, Zettl A, Zhang Z, Çavuşoğlu R, Rohland N, Pinhasi R, Reich D, Davtyan R (2022) The genetic history of the Southern Arc: A bridge between West Asia and Europe. Science 377: eabm4247

Ledgeway A Greek Disguised as Romance? The Case of Southern Italy. 5th International Conference on Greek Dialects and Linguistic Theory In: Janse M, Joseph B, Ralli A, Bagriacik M, editors, 2013. Patras. Laboratory of Modern Greek Dialects, University of Patras 184-228

Ledgeway AN, Schifano N, Silvestri G (2018) Il contatto tra il greco e le varietà romanze nella Calabria meridionale. Lingue Antiche e Moderne 7: 95-133

Manolessou I (2005) The Greek dialects of Southern Italy: an overview. KAMPOS: Cambridge papers in Modern Greek 13: 103-125

Martino P (2009) L'affaire Bovesìa. Un singolare irredentismo. In: Consani C, Desideri P, Guazzelli F, Perta C (eds) Alloglossie e comunità alloglotte nell'Italia contemporanea. Bulzoni Roma, pp 251-275

Miglietta A, Sobrero A (2006) Politica linguistica e presenza del grico in Salento oggi. In: Guardiano C, Calaresu E, Carli A, Robustelli C (eds) Lingue, istituzioni, territori. Bulzoni, Roma, pp 209-226

Miglietta A, Sobrero A (2007) Spie morfologiche della resistenza del grico in Salento. Riv Ital Dialettologia 31: 19-28

Morosi G (1870) Studi sui Dialetti Greci Della Terra d'Otranto. Ed. Salentina, Lecce

Parlangeli O (1953) Sui dialetti romanzi e romaici del Salento. Memorie dell'Istituto Lombardo di Scienze e Lettere 25/3: 93-198

Ralli A (2006) Syntactic and morpho-syntactic phenomena in Modern Greek dialects: the state of the art. J Greek Linguist 7: 121-159

Rohlfs G (1924) Griechen und Romanen in Unteritalien. L.S. Olschki, Geneve

Rohlfs G (1972) Nuovi scavi linguistici nell'antica Magna Graecia. Istituto siciliano di studi bizantini e neoellenici, Palermo

Romano A, Marra PP (2008) Il griko nel terzo millennio: 'speculazioni' su una lingua in agonia. Parabita: Il Laboratorio

Sobrero AA, Romanello MT (1977) Una ricerca sociolinguistica in Salento. In: Simone G, Ruggiero G (eds) Aspetti sociolinguistici dell'Italia contemporanea, Roma, pp 373-384

Squillaci MO (2017) When Greek meets Romance: A morphosyntactic analysis of language contact in Aspromonte

Overview of population inputs in GS (AN)

This section is meant as a short, far from exhaustive, compendium of pieces of literature dealing with known episodes or historical events and processes with potential implications on the composition of the population of southern Apulia and the geographic subregion coinciding with modern GS.

The sole aim of this section is to illustrate the complexity of demographic processes in the area as reported by different authors, without entering historiographical debates.

The different paragraphs, in italics between quotes, were drawn as such from the indicated references, with no further comments. For the sake of clarity, we added modern names for cities, and few minor notes in brackets. Original texts in languages other than English were automatically translated with the Office 365 translator.

*“The Neolithic peopling is early, widespread and intense in Puglia. Open-air sites are occupied on the coasts and hills of the interior, while caves continue to be used, both for the burial of the dead and for the performance of religious cults. At least from the sixteenth century BC until almost the end of the millennium, the coasts of peninsular Italy and the large islands were frequented by sailors and merchants of Mycenaean culture, coming from mainland Greece and the Aegean islands…*
*The Mycenaean presence is attested in Puglia in more than twenty locations, among which the site of Scoglio del Tonno, in Taranto, stands out…*
*Towards the end of the twelfth century B.C., the Apennine civilization of peninsular Italy and the Mycenaean civilization of Greece entered an irreversible crisis, the causes of which are still the subject of discussion among scholars (invasions of other peoples, earthquakes, internal revolutions…*
*The situation stabilized again from the tenth century, that is, with the beginning of the Iron Age."*
(De Juliis 2014), automated translation from Italian.
See also (Vagnetti 2012)

*“Iapygians (Iapyghes) and Iapygia (Iapighia) are the ethnic names used in Greek sources beginning with the sixth century B.C.E. Milesian historian Hecataeus to refer generally to the indigenous populations who dwelt in the hinterland of the Greek colony of Taras [modern Taranto], that is, the region corresponding to modern Apulia. According to Greek tradition, the Iapygians were divided into three distinct populations: the Messapians (Messapioi) in the southern part of the region, extending from Leuca (Iapygian Cape) as far as Cailia and Egnatia; the Peucetians in western and central Apulia, from the river Bradano as far as the Murge and the Ofanto (Aufidus) river; and the Daunians further north, as far as the Gargano Peninsula. This is the picture clearly offered in a passage by the Hellenistic writer Nicander…*
*To begin with, we find a number of tales about the origins of these peoples and their centers. For example, at least from the fifth century B.C.E., as regards the Iapygians, the Sallentini, and several of their cities (Brundisium [modern Brindisi], Hydrous-Otranto, Lupiae-Lecce, Hyria-Oria or Vereto, and so on), the sources refer mainly to a noble and*

*ancient Cretan origin and to famous heroes of Greek mythology, such as Minos, Daedalus, Theseus, and Idomeneus….*
*In later texts of the Hellenistic and Roman period we also find traditions combining Cretan and Illyrian [modern Albania] origins"*
(Lombardo 2014), p.36; this reference contains transcriptions from works of ancient historians.

*"The contacts of the Iapygians, at least those of Messapia, with the Greeks must have been, therefore, intense and continuous throughout the second half of the eighth century [BCE], without leading, however, to real foundations of colonies. In fact, it is only at the end of that century, around 706 BC, that Taranto was created by colonists from Sparta. This was to be the only colony founded by the Greeks in Apulia, perhaps because of the objective difficulty of conquering a territory densely inhabited by well-organized and warlike people.*
*It also had an immediate effect on the civilization of the neighboring Messapians, who drew important cultural contributions from it, distinguishing themselves more and more from the other two Iapygian tribal groups.*
*From this passage of Antiochus of Syracuse it is evident, therefore, that the foundation of Taranto did not arise from a deliberate decision of Sparta, but was due to a group of Spartan rebels, the Parthenians, discriminated against in their homeland for their irregular birth and forced, therefore, to leave their land in search of a new homeland".*
(De Juliis 2014), automated translation from Italian.

*"Closer to Tarentum [Taranto] several small villages were occupied by Greeks. Of these we may mention Satyrium, just east to Tarentum.*
*We might expect to find some native influence in the matter of religion or customs upon the newly come Greeks, with whom there must surely have been intermarriage".*
(Boardman 1999), pp. 184, 190, 191.

*"The Taranto plain in the north-west of the Salento peninsula is the only place in Apulia where evidence has been found of a lasting Greek colonization venture. It was there that the polis of Taras developed, which, like Sybaris, is generally considered to have been one of the most powerful of Megale Hellas [Magna Grecia].*
*Taras' expansion proceeded primarily in the direction of its Salentine neighbours, and it ultimately carved out a territory in the Taranto plain…..*
*If the results of these systematic excavations and surveys are indeed representative of the many incidentally discovered Iron Age sites in Salento, we must conclude that during the 8th century BC the region's native communities engaged in settlement expansion, rural infill and reclamation of previously non-exploited or only marginally exploited landscapes. These processes seem to have involved virtually all major Salento landscape units, including the Taranto plain, and would have conditioned the circumstances in which early Greek colonisation took place".*
(Attema et al. 2010)

*“The political and military importance reached by Taranto in the 4^ century BC when it also assumed the leadership of the Italiote cities, makes this city the true protagonist of the deep and definitive Hellenization of the Apulian territory. This phenomenon includes the whole of the 4^ century and the first decades of the next, until the defeat suffered by the Taranto people in the war with the Romans”.*
(Carratelli 1996), p. 553; automated translation from Italian.

*“Then, only a few years later [334 BCE] the Tarentines had to call for help the Epirote king Alexander the Molossos, uncle of Alexander the Great, against the Messapians and the Lucanians.*
*He arrived in 334 at the request of Taranto and pursued a policy of conquest by successfully fighting the indigenous populations.”*
(Lefèvre 2007; Lombardo 2014)

*“With the end of the Second Samnite War, therefore, Taranto found itself exposed to the attack of the Lucanians, to whom the Romans were now added. In 303, therefore, it was forced to call on the Spartan prince Cleonymus for help to repel the two powerful adversaries. He managed to gather a large army, made up of 20,000 infantry and 2000 cavalry from Taranto, a large number of mercenaries and, again, the contingents sent by three Italiote pòleis and the Messapians. Regarding the latter, it should be noted that it is the first time that the traditional enemies of Taranto appear lined up as its allies against a common enemy, in this case the Romans”*
(De Juliis 2004); automated translation from Italian.

*“About two decades later, not only the Messapians but also the other indigenous peoples of southern Italy, with a few exceptions such as the Daunians of Arpi, supported the Tarentines' appeal to Pyrrhus against the Romans. Not by chance Rome had already taken root in northern Apulia, founding Luceria (313 B.C.E.) and then Venusia (298 B.C.E.). We know what the outcomes and the consequences were of the ensuing war with Pyrrhus (280-272 B.C.E.), bringing all the indigenous peoples of Southern Italy that had fought against it under the submission of Rome. The last of them to submit were the peoples of southern Apulia, and that required another war, the bellum Sallentinum. Two years of warfare and of Roman triumphs - in 267 B.C.E. "de Sallentineis" and the next year "de Sallentineis Messapieisque" (Fasti Triumphales Capitolini, II, XX) - resulted in the Roman conquest of Brundisium, whose magnificent natural harbor was to become with the foundation of a Latin colony in 243 B.C.E. and the subsequent completion of the Via Appia from Taras to Brundisium [modern Brindisi], the most important naval base for Roman expansion toward the Balkan Peninsula and Eastern Mediterranean”.*
(Lombardo 2014)

*“In December of 62 [BCE], after an absence of nearly six years, Pompey arrived in Brundisium. The general was generous to his forces, conducted a final assemblage on the shores of Italy and the ostentatiously dismissed every man to his home”.*
(Gruen 1974)

“*Jewish settlements in Apulia were very ancient. According to medieval tradition, they were founded by prisoners deported to Italy by Emperor Titus, after the destruction of the second temple. In two manuscripts of the so-called Josippon, for example, there is the addition that it would have been about 90,000 men, of whom about 5,000 settled in Taranto, Otranto and other cities of Puglia….*
*Conversions of Chrisitians and Jews went both ways in Bari.*
*Perhaps the Jewish community was Greek-speaking, an hypothesis which might be confirmed by the fact that the president of the community was protus”.*
(von Falkenhausen 2022), pp. 155, 300

*“After the first immigration of Hellenized Syrians and Egyptians in the seventh century, there came a second such influx in the following century, this time from Constantinople and Greece, refugees from the iconoclastic oppression of the Byzantine Emperors, Leo III the Isaurian and his son, Constantine V Copronymus (725-775). Some scholars have set the numbers of monks, ecclesiastics, and laymen who now found their way into southern Italy as high as 50,000. At any rate a larger number now appears to have come than in the seventh century. Coming from Constantinople and Greece, they strengthened and diversified the Greek element in Calabria. They also brought with them the Basilian and Chrysostomine liturgies of Constantinople. In the middle of the eighth century, apparently, the Isaurian emperors, provoked by Roman opposition to iconoclasm and by the papal alliance with the new Carolingian monarchy, detached eastern Illyricum, Calabria, and Sicily from the jurisdiction of Rome and transferred them to that of the patriarchate of Constantinople”*.
(Setton 1956)

*“In 537 [CE] 800 Thracian knights landed in Otranto under the command of John and a thousand conscripted cavalry under the command of Alexander and Marcetius”*
(von Falkenhausen 2022), p. 143; automated translation from Italian

*“After the Byzantine presence in the region in late antiquity weakened significantly following wars with the Goths and Longobard invasions, Basil I reconquered it about 870 and initiated population transfers from the Peloponnese and the Pontus.*
*The notion of continued Orthodox religious vitality is strengthened by the high numbers of post-Byzantine devotional and funerary texts. These painted and incised supplications to the Lord and, later, to the Virgin and saints, coupled with records of the dead in graffiti and on tombstones, are vivid testimonies to the regional endurance of Greek and the continued vitality of the Greek-speaking community”.*
(Safran 2015)

*[Period 873-888 CE] “Greeks from Pontic Herakleia settle at Gallipoli.*
*Freedmen given to Basil by Danelis (i.e. from the Peloponnese) settled in Italy.*
*More freedmen from legacy of Danelis settled in Italy”.*
(McCormick 1998)

*“The military conquest of southern Italy was followed by a certain repopulation of the territory by freedmen from a large estate in the Peloponnese”.*
(von Falkenhausen 2022)

*“The military interventions associated with the names of Gregory protospatharios, Prokopios protovestiarios and Nikephoros Phokas the Elder coincided with an apparently brief burst of administered migration of Byzantine populations, including freedmen, from the east into southern Italy”.*
(McCormick 1998)

*“From Theoph. cont. it is not clear whether S. [Stephanos Maxentios] was Strategos of the theme of Langobardia (according to Falkenhausen) and in this capacity commanded additional troops sent from the east of the empire in 882 or whether he was commander of an ad hoc army with which he was sent to Lower Italy. In the latter case, he may have been the successor of Procopius (# 26758) or Leon Apostyppes (# 24341), who had been strategos of the two themes of Thrace and Macedonia before his deposition. It is noteworthy that these two themes apparently fought in full force in Lower Italy, while Charsianon and Cappadocia assigned only a few elite regiments”.*
(Prosopographie der mittelbyzantinischen Zeit Online#27223)

*“Nikephoros remained in command of Charsianon until his appointment as the commander-in-chief (monostrategos, "single-general") against the Arabs in southern Italy in replacement of Stephen Maxentios, who had been defeated by the Arabs. This took place in 885, according to traditional dating. It is likely, however, that Nikephoros was originally sent to Italy already before that, at the head of a picked detachment of troops from Charsianon, which Theophanes Continuatus records as part of Maxentios' expeditionary force. His command involved the forces of several western themes (Thrace, Macedonia, Cephallenia, Longobardia and Calabria), but Theophanes Continuatus also reports that Nikephoros received further reinforcements from the themes of Asia Minor, including a Paulician detachment. Nikephoros' command in Italy lasted until his recall to Constantinople following the accession of Leo VI the Wise, in late 886”.*
(https://en.wikipedia.org/wiki/Nikephoros_Phokas_the_Elder)

*“The Paulicians were a religious sect and as such probably included elements of different ethnic origin, but the majority were no doubt Armenians. When finally during the reign of Basil I (867-886) their strongholds were taken and razed to the ground, their army defeated and their leader killed (872), they were forced to abandon their homes and were settled elsewhere in the Empire. We know that some of them were settled in Southern Italy, in the regions under the jurisdiction of the Empire.*
*The repeopling and economic rehabilitation of the country were no doubts the reason for the numerous transfers made in the eight century”.*
(Charanis 1946)

*"Southern Italy was retained against Moslem efforts. Basil I organized the region called Longobardia in the south- east more or less as a Byzantine province or "theme" and, as part of his reassertion of Byzantine authority, sent some three thousand Greek (and presumably Slavic) colonists into this area, where they appear to have been settled in the region west of Bari".*
(Setton 1956)

*[10th century CE]…"there was a continuous influx to Bari of soldiers and officials from Constantinople and other provinces. Often the governors brought their children and relatives with them, and although generally the catepans remained in Italy for a short time, their employees could also remain there for decades.*
*Among the immigrants there were a large number of Armenians. In fact, important Armenian contingents had taken part in the Byzantine reconquest of southern Italy and then had been settled in areas depopulated by the long periods of civil war and Arab raids".*
(von Falkenhausen 2022), p. 79

*"Early in the summer of 1480, Gedik Ahmed Pasha, received orders from the sultan to cross the straits of Otranto from Vlore [in Albania] to Apulia, only some forty-six miles distant. He… was at the head of a fleet consisting of 140 vessels-40 galleys, 60 one-masters, and 40 freight ships-and bearing an army of 18,000 soldiers (exaggerated to 100,000 in some accounts) and 700 horses. The devastating invasion is described in every conceivable detail in native Apulian sources".*
*(Babinger 1978) [see also https://en.wikipedia.org/wiki/Ottoman_conquest_of_Otranto for details on the troops involved on both sides]*


Attema PAJ, Burgers G-JLM, van Leusen PM (2010) Rethinking early Greek - indigenous encounters in southern Italy. Regional pathways to complexity. Amsterdam University Press, pp 119-134
Babinger F (1978) Mehmed the Conqueror and his time. Princeton University Press
Boardman J (1999) The Greeks overseas : their early colonies and trade. 4th ed. Thames and Hudson, London, U.K.
Carratelli GP (1996) I Greci in Occidente. Bompiani, Firenze, Italy
Charanis P (1946) On the question of the Hellenization of Sicily and Southern Italy during the Middle Ages. Am Hist Rev 52: 74-86
De Juliis EM (2004) Greci e Italici in Magna Grecia: Un rapporto difficile. Laterza, Bari, Italy
De Juliis EM (2014) Popoli e culture della Puglia preromana. La preistoria, le genti indigene, i coloni greci. In: Salvemini B, Massafra A (eds) Storia della Puglia. 1. Dalle origini al Seicento. Gius. Laterza & Figli Spa
Gruen ES (1974) The last generation of the Roman Republic. Univ of California Press
Lefèvre F (2007) Histoire du mond grec antique. Librairie Générale Francaise, Paris
Lombardo M (2014) Iapygians: The indigenous populations of ancient Apulia in the fifth and fourth centuries BCE. In: Carpenter TH, Lynch KM, Robinson EGD (eds) The Italic People of Ancient Apulia. New Evidence from Pottery for Workshops, Markets, and Customs. Cambridge University Press, Cambridge, U.K., pp 36-68
McCormick M (1998) The Imperial Edge: Italo-Byzantine identity, movement and integration, A.D. 650-950. In: Ahrweiler H, Laiou AE (eds) Studies on the internal diaspora of the Byzantine Empire. Dumbarton Oaks, Washinghton, D.C.
Safran L (2015) Greek in the Salento: Byzantine and Post-Byzantine public texts. In: Rhoby A (ed) Inscriptions in Byzantium and beyond. Austrian Academy of Sciences Press, pp 227-240
Setton KM (1956) The Byzantine Background to the Italian Renaissance Proc Am Phil Soc 100: 1-76

Vagnetti L (2012) Western Mediterranean. In: Cline EH (ed) The Oxford Handbook of the Bronze Age Aegean. Oxford University Press, Oxford, U.K., pp 890–906
von Falkenhausen V (2022) Studi sull'Italia bizantina. Viella, Rome, Italy

#### Methods

-Sampling procedures

Grecìa Salentina. (VS)

We used samples collected in two temporally distinct campaigns.

The first one was performed in 1994. Exclusively based on DNA quality, eight (7M, 1F) samples from this campaign entered the study (20-27 in Table S1). The use of these samples in genomic studies was approved by the Ethical Committee Fondazione IRCCS Policlinico San Matteo (protocol number 0028298/22).

The second one began in 2019 and is still ongoing. Biological sample collection within the GS area was conducted across several municipal sessions, facilitated by local authorities who provided administrative spaces for participant engagement. Recruitment and sampling efforts spanned various centers, with the following subject distributions: Sternatia (24), Calimera (18), Martano (9), Castrignano de' Greci (8), Martignano (5), Soleto (3), Corigliano d'Otranto (3), Zollino (2), and Carpignano Salentino (1). Research procedures and the form for informed consent were approved by the local Ethics Committee (Comitato etico ASL Lecce, verbali n. 34 4/7/2019; n. 35 del 25/7/2019; n. 41 14/1/2020). Exclusively based on DNA quality, nineteen (13M, 6F) samples from this campaign entered the study (1-19 in Table S1). Eleven out of the 19 participants from this series reported knowledge and use of the Griko language.

This design was applied to control for gross shifts among localities and across generations. In both cases, the aims and scopes of the project were illustrated to the participants and their communities. Prior to enrollment, all individuals received a detailed briefing regarding the study's scientific objectives and subsequently provided their written informed consent.

All the research was performed in accordance with the relevant guidelines and regulations reported in the abovementioned documents and those agreed upon by the scientific community. In this respect, because the present work did not involve any issue relevant for the donor's health, only the relevant prescriptions of the WMA Declaration of Helsinki and COE Oviedo Convention were obeyed.

Biological samples (buccal swabs) were anonymized upon collection. DNA was prepared with standard methods.

The project received the endorsement of the Union of the Municipalities (https://www.unionegreciasalentina.le.it/).

Internal controls. (AN, CJ, FC, NA)

Twentyone samples (9 Italian_CTRL, 6 Greek_CTRL and 6 Turkish_CTRL, respectively) were included in the sequencing pipeline as controls. For Italy and Türkiye, they were selected from regions mainly outside the influence of ancient Greek colonies.

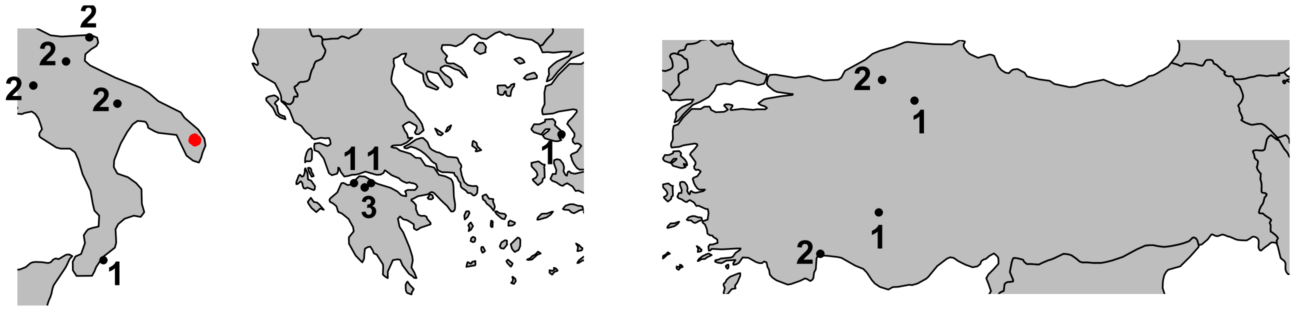


Maps of Southern Italy (left), Greece (center) and Türkiye (right) showing the provenance of the control samples sequenced in this work. Numbers indicate samples from the same location.GS is shown in red as reference.

High depth genome sequencing (AN, EDA)

-Library construction, alignment and variant calling

Each sample was prepared according to the Illumina TruSeq DNA sample preparation guide to obtain a final library with a 300-400 bp average insert size. Multiple indexing adapters were ligated to the ends of the DNA fragments to prepare them for hybridization onto a flow cell.

The BCL/cBCL (base call) binary files were converted into FASTQ files using the Illumina package bcl2fastq2-v2.20.0. The demultiplexing option (--barcode-mismatches) was set to perfect match (value: 0).

Paired-end sequences generated by the HiSeq instrument were mapped to the human genome using iSAAC aligner (iSAAC-04.18.11.09 (c) 2010-2017 Illumina, Inc.) with the UCSC assembly hg38 (Dec. 2013) reference sequence.

Strelka (2.9.10 (c) 2009-2018 Illumina, Inc.) was used to identify single-nucleotide variants (SNVs) and short insertions and deletions (indels). The final coverage was 25.6x, on average, with only two samples below 20x. Summary results of the sequencing effort are reported in Table S1.

The multisample vcf file was lifted over to GRCh37/hg19 with picard (Broad Institute 2019).

-Reference WGS dataset assembly (GC, AN)

Five whole genome datasets were included in the study as reference:
1,000 genomes (Byrska-Bishop et al. 2022). The data were downloaded from https://ftp.1000genomes.ebi.ac.uk/vol1/ftp/data_collections/1000G_2504_high_coverage/working/20201028_3202_phased/ as VCF files in GRCh38/hg38. We first selected four European populations (IBS, TSI, GBR, FIN). CEUs were superseded by the French represented in HGDP and SGDP (see below). The chromosome-specific files were concatenated and lifted to GRCh37/hg19 with GATK LiftoverVcf, resulting in 69,777,498 sites in 404 samples.

HGDP (Bergström et al. 2020). This dataset was obtained in VCF format as a subset of the dataset described in (D'Atanasio et al. 2023), containing genotypes for chromosomes 1-22 and lifted-over to GRCh37/hg19 with respect to the original. We selected 249 subjects from regions including Europe, the Caucasus, Sardinia, the Middle East and North Africa. We used French subjects from this dataset to supersede the 1,000 Genomes CEU's.

SGDP (Mallick et al. 2016). We downloaded the 279 fully public genomes in GRCh37/hg19 from https://reichdata.hms.harvard.edu/pub/datasets/sgdp/ as VCF files. We selected 56 subjects from regions including Europe, the Caucasus, Sardinia, Middle East and North Africa.

EGAD0001001440. This dataset was obtained on the basis of Wellcome Sanger Institute Data Access Agreement and was downloaded as CRAM files in GRCh37/hg19. It includes 100 Greek samples originally described in (Gilly et al. 2018), sequenced at an average depth 30x on ILLUMINA,HiSeq X Ten, paired end. Variant calling was performed with GATK Haplotype Caller, (Van der Auwera and O'Connor 2020) by relaxing the "heterozygosity" parameter (actually a prior on the rate of variant occurrence per base pair) to 0.002, to match the recommendations by (Mallick et al. 2016) for population studies. Individual gVCF files were first collated for each chromosome and then concatenated for all autosomes in a single VCF file. Information on geographic provenance was limited to Attica (Central-Eastern Greece) for 19 samples and generic Greece for the remaining 81 samples.

The MinE dataset. This dataset (Project MinE ALS Sequencing Consortium 2018; van Rheenen et al. 2021)was obtained on the basis of a data sharing agreement and was provided as plink1.9 files (Purcell et al. 2007) in GRCh37/hg19. It includes samples originally described in (Kars et al. 2021), with technical details reported therein.
The initial dataset consisted of 847 subjects, partitioned as follows:
69 and 613 Italians and Turkish subjects, respectively, reported as affected by conditions listed in the original publication;
1 and 141 Italians and Turkish subjects, respectively, reported as non-affected;
3 and 20 Italians and Turkish subjects, respectively, with missing condition;
From these, we selected a subset of 211 subjects, consisting of the 141 Turkish non-affected and the 70 Italians with known but blinded affection status, to keep the size of the Italian and Turkish subsets in line with other national groups.
Information on the precise geographic provenance was missing and all samples were assigned the same geographic coordinates (35.5E; 38.7N and 12.5E; 41.9N, for Turkish and Italians, respectively).

-Preliminary data cleaning

All five datasets were restricted to sites within the 20141020.strict_mask.whole_genome.bed accessed at ftp.1000genomes.ebi.ac.uk/vol1/ftp/release/20130502/supporting/accessible_genome_masks/

Using BCFtools (Li 2011) statistics, samples were filtered out from each dataset, separately, if one or more of the following conditions applied:
-corrupted data file(s);
-outlier for average depth;
-outlier for the overall n. of SNPs;
-outlier for the transition/transversion ratio;
-outlier for heterozygous/homozygous sites ratio;
-outlier for the n. of singletons

The five datasets were merged with BCFtools, without assumptions on genotypes at untyped positions in any of the dataset (i.e. without using the missing2ref option). This resulted in 134M sites in 1002 samples (hereafter dubbed the 134M dataset).

Sites were then filtered out if the fraction of missing genotypes exceeded 2% and if they were indel variants (different allele length between ALT and REF). In addition, only strictly biallelic SNPs were retained. This resulted in 4,374,664 SNPs in 1002 individuals (hereafter dubbed the 4.4M dataset).

A preliminary analysis of diversity in this dataset revealed outlying population groups that obscured the subtle variations within the Mediterranean space (e.g. HGDP Russians, Bedouins and Mozabites, 1,000genomes FIN and SGDP Estonians and Saharawis), in line with previous analyses (Lao et al. 2008; Behar et al. 2010; Skoglund et al. 2014).

After the removal of the above samples, the 4.4M dataset was further filtered, by applying LD-pruning with an r2 threshold of 0.5 (plink2 –indep-pairwise 200 50 0.5), controlling for Hardy-Weinberg equilibrium (85 positions with p<1e-6 removed), and retaining sites with MAF >= 0.03. This resulted in 503,578 sites in 799 samples (hereafter dubbed the 500K dataset, Table S2). As this dataset was devoid of representatives of African-related ancestry, it was integrated with the following groups for some analysis, by keeping the same list of variable sites:
-The 25 HGDP Mozabites for ADMIXTURE analysis;
-Eight HGDP Mbuti as a distant outgroup in the f3/D statistics calculations.

The 500K dataset consisted of 34 ID-identified groups of samples (Table S2). In some instances, samples with the same geographic provenance were kept separate, e.g. when a more specific geographic assignment was available for a subset of them. Note that in many cases multiple ID’s for the same population or multiple samples within the same population_ID derived from different sources and were sequenced on different platforms. This enabled a powerful control against batch effects potentially generated at any step leading to the final individual genotype description.

In the presentation of some analyses samples from the 34 Population_ID’s were lumped into 12 broader areas, i.e. Caucasus, C-Europe, Greece, Italy, Italy-GS, the Middle-East, N-Europe, Other-Balkan, Sardinia, SW-Asia, Türkiye and W-Europe (Table S2).

Throughout the text we will use the term “sample” to indicate an individual, and “population_ID” to indicate a group of individuals reported from the same national population and the same or different sequencing source.

Behar DM, Yunusbayev B, Metspalu M, Metspalu E, Rosset S, Parik J, Rootsi S, Chaubey G, Kutuev I, Yudkovsky G, Khusnutdinova EK, Balanovsky O, Semino O, Pereira L, Comas D, Gurwitz D, Bonne-Tamir B, Parfitt T, Hammer MF, Skorecki K, Villems R (2010) The genome-wide structure of the Jewish people. Nature 466: 238-242
Bergström A, McCarthy SA, Hui R, Almarri MA, Ayub Q, Danecek P, Chen Y, Felkel S, Hallast P, Kamm J, Blanché H, Deleuze J-F, Cann H, Mallick S, Reich D, Sandhu MS, Skoglund P, Scally A, Xue Y, Durbin R, Tyler-Smith C (2020) Insights into human genetic variation and population history from 929 diverse genomes. Science 367: eaay5012
Broad Institute (2019) Picard Toolkit. Broad Inst. GitHub Repos.
Byrska-Bishop M, Evani US, Zhao X, Basile AO, Abel HJ, Regier AA, Corvelo A, Clarke WE, Musunuri R, Nagulapalli K (2022) High-coverage whole-genome sequencing of the expanded 1000 Genomes Project cohort including 602 trios. Cell 185: 3426-3440. e3419
D'Atanasio E, Risi F, Ravasini F, Montinaro F, Hajiesmaeil M, Bonucci B, Pistacchia L, Amoako-Sakyi D, Bonito M, Onidi S, Colombo G, Semino O, Destro Bisol G, Anagnostou P, Metspalu M, Tambets K, Trombetta B, Cruciani F (2023) The genomic echoes of the last Green Sahara on the Fulani and Sahelian people. Curr Biol 33: 5495-5504.e5494
Gilly A, Suveges D, Kuchenbaecker K, Pollard M, Southam L, Hatzikotoulas K, Farmaki AE, Bjornland T, Waples R, Appel EVR, Casalone E, Melloni G, Kilian B, Rayner NW, Ntalla I, Kundu K, Walter K, Danesh J, Butterworth A, Barroso I, Tsafantakis E, Dedoussis G, Moltke I, Zeggini E (2018) Cohort-wide deep whole genome sequencing and the allelic architecture of complex traits. Nat Commun 9: 4674
Kars ME, Başak AN, Onat OE, Bilguvar K, Choi J, Itan Y, Çağlar C, Palvadeau R, Casanova JL, Cooper DN, Stenson PD, Yavuz A, Buluş H, Günel M, Friedman JM, Özçelik T (2021) The genetic structure of the Turkish population reveals high levels of variation and admixture. Proc Natl Acad Sci U S A 118: e2026076118
Lao O, Lu TT, Nothnagel M, Junge O, Freitag-Wolf S, Caliebe A, Balascakova M, Bertranpetit J, Bindoff LA, Comas D, Holmlund G, Kouvatsi A, Macek M, Mollet I, Parson W, Palo J, Ploski R, Sajantila A, Tagliabracci A, Gether U, Werge T, Rivadeneira F, Hofman A, Uitterlinden AG, Gieger C, Wichmann H-E, Rüther A, Schreiber S, Becker C, Nürnberg P, Nelson MR, Krawczak M, Kayser M (2008) Correlation between genetic and geographic structure in Europe. Curr Biol 18: 1241-1248
Li H (2011) A statistical framework for SNP calling, mutation discovery, association mapping and population genetical parameter estimation from sequencing data. Bioinformatics 27: 2987-2993
Mallick S, Li H, Lipson M, Mathieson I, Gymrek M, Racimo F, Zhao M, Chennagiri N, Nordenfelt S, Tandon A, Skoglund P, Lazaridis I, Sankararaman S, Fu Q, Rohland N, Renaud G, Erlich Y, Willems T, Gallo C, Spence JP, Song YS, Poletti G, Balloux F, van Driem G, de Knijff P, Romero IG, Jha AR, Behar DM, Bravi CM, Capelli C, Hervig T, Moreno-Estrada A, Posukh OL, Balanovska E, Balanovsky O, Karachanak-Yankova S, Sahakyan H, Toncheva D, Yepiskoposyan L, Tyler-Smith C, Xue Y, Abdullah MS, Ruiz-Linares A, Beall CM, Di Rienzo A, Jeong C, Starikovskaya EB, Metspalu E, Parik Jr, Villems R, Henn BM, Hodoglugil U, Mahley R, Sajantila A, Stamatoyannopoulos G, Wee JTS, Khusainova R, Khusnutdinova E, Litvinov S, Ayodo G, Comas D, Hammer MF, Kivisild T, Klitz W, Winkler CA, Labuda D, Bamshad M, Jorde LB, Tishkoff SA, Watkins WS, Metspalu M, Dryomov S, Sukernik R, Singh L, Thangaraj K, Pääbo S, Kelso J, Patterson N, Reich D (2016) The Simons Genome Diversity Project: 300 genomes from 142 diverse populations. Nature 538: 201-206
Project MinE ALS Sequencing Consortium (2018) Project MinE: study design and pilot analyses of a large-scale whole-genome sequencing study in amyotrophic lateral sclerosis. Eur J Hum Genet 26: 1537-1546
Purcell S, Neale B, Todd-Brown K, Thomas L, Ferreira MAR, Bender D, Maller J, Sklar P, de Bakker PIW, Daly MJ, Sham PC (2007) PLINK: A tooset for whole-genome association and population-based linkage analyses. Am J Hum Genet 81: 559-575
Skoglund P, Malmström H, Omrak Aa, Raghavan M, Valdiosera C, Günther T, Hall P, Tambets K, Parik Jr, Sjögren K-Gr, Apel J, Willerslev E, Stora J, Götherström A, Jakobsson M (2014) Genomic diversity and admixture differs for stone-age Scandinavian foragers and farmers. Science 344: 747-750
Van der Auwera GA, O'Connor BD (2020) Genomics in the cloud: Using Docker, GATK and WDL in Terra (1st ed.). O'Reilly Media
van Rheenen W, van der Spek RAA, Bakker MK, van Vugt J, Hop PJ, Zwamborn RAJ, de Klein N, Westra HJ, Bakker OB, Deelen P, Shireby G, Hannon E, Moisse M, Baird D, Restuadi R, Dolzhenko E, Dekker AM, Gawor K, Westeneng HJ, Tazelaar GHP, van Eijk KR, Kooyman M, Byrne RP, Doherty M, Heverin M, Al Khleifat A, Iacoangeli A, Shatunov A, Ticozzi N, Cooper-Knock J, Smith BN,

Gromicho M, Chandran S, Pal S, Morrison KE, Shaw PJ, Hardy J, Orrell RW, Sendtner M, Meyer T, Başak N, van der Kooi AJ, Ratti A, Fogh I, Gellera C, Lauria G, Corti S, Cereda C, Sproviero D, D'Alfonso S, Sorarù G, Siciliano G, Filosto M, Padovani A, Chiò A, Calvo A, Moglia C, Brunetti M, Canosa A, Grassano M, Beghi E, Pupillo E, Logroscino G, Nefussy B, Osmanovic A, Nordin A, Lerner Y, Zabari M, Gotkine M, Baloh RH, Bell S, Vourc'h P, Corcia P, Couratier P, Millecamps S, Meininger V, Salachas F, Mora Pardina JS, Assialioui A, Rojas-García R, Dion PA, Ross JP, Ludolph AC, Weishaupt JH, Brenner D, Freischmidt A, Bensimon G, Brice A, Durr A, Payan CAM, Saker-Delye S, Wood NW, Topp S, Rademakers R, Tittmann L, Lieb W, Franke A, Ripke S, Braun A, Kraft J, Whiteman DC, Olsen CM, Uitterlinden AG, Hofman A, Rietschel M, Cichon S, Nöthen MM, Amouyel P, Traynor BJ, Singleton AB, Mitne Neto M, Cauchi RJ, Ophoff RA, Wiedau-Pazos M, Lomen-Hoerth C, van Deerlin VM, Grosskreutz J, Roediger A, Gaur N, Jörk A, Barthel T, Theele E, Ilse B, Stubendorff B, Witte OW, Steinbach R, Hübner CA, Graff C, Brylev L, Fominykh V, Demeshonok V, Ataulina A, Rogelj B, Koritnik B, Zidar J, Ravnik-Glavač M, Glavač D, Stević Z, Drory V, Povedano M, Blair IP, Kiernan MC, Benyamin B, Henderson RD, Furlong S, Mathers S, McCombe PA, Needham M, Ngo ST, Nicholson GA, Pamphlett R, Rowe DB, Steyn FJ, Williams KL, Mather KA, Sachdev PS, Henders AK, Wallace L, de Carvalho M, Pinto S, Petri S, Weber M, Rouleau GA, Silani V, Curtis CJ, Breen G, Glass JD, Brown RH, Jr., Landers JE, Shaw CE, Andersen PM, Groen EJN, van Es MA, Pasterkamp RJ, Fan D, Garton FC, McRae AF, Davey Smith G, Gaunt TR, Eberle MA, Mill J, McLaughlin RL, Hardiman O, Kenna KP, Wray NR, Tsai E, Runz H, Franke L, Al-Chalabi A, Van Damme P, van den Berg LH, Veldink JH (2021) Common and rare variant association analyses in amyotrophic lateral sclerosis identify 15 risk loci with distinct genetic architectures and neuron-specific biology. Nat Genet 53: 1636-1648

#### Data analyses

1. Other datasets.

-The population dataset generated by (Paschou et al. 2014) was used to precisely put our GS samples in the context of subpopulations of the Balkan peninsula, Crete, Aegean area and Cappadocia. The original dataset consisted of 75194 sites typed with the Illumina OMNI 2.5 or OMNI1-QUAD chip platforms and filtered as described in the original paper. Allele encoding in the original plink files and liftover from NCBI36/hg18 caused the reduction of usable SNPs to 25645 after merging with our GS samples (hereafter 25K dataset)..

- We also used two lists of variants for specific purposes. The set of 997 variants proposed as useful ancestry informative markers (AIMs) for the European population (Drineas et al. 2010) was considered (hereafter 1K dataset). This variant set is largely independent from the 500K dataset, as far as only 78 sites are shared between the two and thus represents a valid control for the results obtained on the latter. The list was downloaded from https://www.cs.purdue.edu/homes/pdrineas/documents/POPRESAIMS/
All variants for the 799 samples were extracted from the 4.4 M dataset.

- In order to get a temporal dissection of the diversity patterns in our analyses, we considered variants reported by (Albers and McVean 2020). We downloaded the chromosome-specific csv files from http://human.genome.dating/download/index and concatenated them to obtain a list of 8,032,251 variants with an estimated age of 300 generations or less, potentially useful to inform on the last 9,000 years. We considered the age figures obtained with the joint method, which takes into account both the molecular and recombination clocks (see original paper). As the vast majority of these young variants have a low frequency, they were lost during the filtering process leading to the 500K dataset. We thus extracted them from the 134M dataset. As different variant calling procedures used in the 5 reference datasets affected severely the representation of rare alleles in the 134M dataset, we limited the subsequent analyses to our 27 GS and 21 control samples, which underwent a homogeneous variant calling pipeline. Overall, we obtained 11589 variable sites with known ancestral allele, of which 11,510, 4563, 4660 and 4085 had non-null frequency in the GS, Italian, Greek and Turkish controls, respectively  We considered variants in the age intervals 300-200 (n=7661), 200-100 (n=3584), 100-70 (n=272) and <70 (n=72) generations. From these sets we counted the occurrence of ancestral alleles in GS, Italian_CTRL (downsampled to 6), Greek_CTRL and Turkish_CTRL and represented the sharing as Venn diagrams. In order to measure the sharing conditioned on the allele frequency, we calculated Weir and Cockeram's Fst on the same sets, without downsampling the Italian_CTRL, with plink2.0.

2. Principal Component Analysis - PCA (GC, AN). This was performed in all cases with EIGENSOFT SMARTPCA (Patterson et al. 2006). The relevant vcf files were imported in plink and converted to the EIGENSTRAT format with CONVERTF. The exclusion of outliers was disabled with the -m flag, and 20 eigenvectors were requested in output. In all runs all samples were used for PC calculation. The resulting evec files were imported in R for visualization and further analysis.

3. EXPECTED SAMPLING LOCATION (GC, AN) We observed that in all PCA's multiple PC's conveyed relevant geographic information. To exploit the combined power of

these relationships we used multiple regression for longitude and latitude, separately. The true sampling coordinate for each subject was used as dependent variable and regressed against the first 20 PC's (Coordinate ~ PC1:PC2 + intercept). Each sample was then assigned the corresponding value expected from the regression, which then represents an expectation conditional on the overall composition of the dataset (Table S3).

4. Admixture (GC, AN). This was performed with the program ADMIXTURE (Alexander et al. 2009) on the set of 799 samples plus the 25 Mozabite and variants in the 500K dataset, with 5 replicates and random seed. The K parameter was varied between 2 and 12 and results checked with the cross-validation plot. The results for each value of K were condensed with pong (Behr et al. 2016), which provides the fraction of highly similar replicates.

5. IBD (GC, AN) Estimation of the sharing of chromosomal fragments identical by descent was performed with IBIS (Seidman et al. 2020) with the genetic map obtained at https://github.com/adimitromanolakis/geneticMap-GRCh37 and the 500K dataset. IBIS was used with the -min_l 3 and -maxDist 0.17 options to select IBD fragments of 3 cM or longer and given a maximum distance between adjacent SNPs of 0.17 cM, as recommended. The output was manipulated with R functions. Of the 147,012 fragments identified, 124,003 were 3 to 5 cM long and 23,009 were longer than 5 cM. Error density, i.e. the proportion of SNPs breaking the perfect identity of fragments, was always lower than 0.004. For each of the two subsets we calculated the average amount of cM and the average number of fragments shared in pairwise comparisons between subjects of the same or different Population_IDs, broader areas or GS municipalities (Tables S2, S4).

6. ROH (GC, AN) Runs of homozygosity were estimated with plink1.9 (Purcell et al. 2007) from the 4.4M dataset. The following options were used: --homozygous_snp 20 to set the minimum number of homozygous variants in a ROH; --homozyg-density 50 to consider only chromosomal regions with at least one SNP/50 kb; --homozyg-window-snp 30 as a moving window; --homozyg-gap 1000 as a maximum SNP spacing. The seach started from a minimum ROH length of 500 kb (131,162 segments identified). The output was imported in R to proceed with further analyses and slicing to ROH lengths of 1.5 (5212 segments) and 5 Mb (857 segments), to directly compare with (McQuillan et al. 2008).

7. GS IN THE CONTEXT OF aDNA (EDA, MG) To explore potential source populations for the GS group, PCA was performed (Patterson et al. 2006) on the 27 modern GS genomes and 4021 relevant ancient and modern Eurasian individuals from the Allen Ancient DNA Resource (AADR) v62.0_HO (Mallick et al. 2024; Mallick and Reich 2024). More specifically, PC's were computed using the 27 GS genomes together with 1,275 modern individuals (grey dots) from Northern Europe, Southwestern Europe, the Balkans, the Near East, and the Caucasus. The remaining ancient individuals from the same regions were subsequently projected onto the resulting PCA space.

Albers PK, McVean G (2020) Dating genomic variants and shared ancestry in population-scale sequencing data. PLOS Biol 18: e3000586

Alexander DH, Novembre J, Lange K (2009) Fast model-based estimation of ancestry in unrelated individuals. Genome Res 19: 1655-1664

Behr AA, Liu KZ, Liu-Fang G, Nakka P, Ramachandran S (2016) pong: fast analysis and visualization of latent clusters in population genetic data. Bioinformatics 32: 2817-2823
Drineas P, Lewis J, Paschou P (2010) Inferring geographic coordinates of origin for Europeans using small panels of ancestry informative markers. PLoS ONE 5: e11892
Mallick S, Micco A, Mah M, Ringbauer H, Lazaridis I, Olalde I, Patterson N, Reich D (2024) The Allen Ancient DNA Resource (AADR) a curated compendium of ancient human genomes. Sci Data 11: 182
Mallick S, Reich D (2024) The Allen Ancient DNA Resource (AADR): a curated compendium of ancient human genomes. Harvard Dataverse, V7.0 data release [Nov 16, 2022]. Dataset 2024.
McQuillan R, Leutenegger A-L, Abdel-Rahman R, Franklin CS, Pericic M, Barac-Lauc L, Smolej-Narancic N, Janicijevic B, Polasek O, Tenesa A, MacLeod AK, Farrington SM, Rudan P, Hayward C, Vitart V, Rudan I, Wild SH, Dunlop MG, Wright AF, Campbell H, Wilson JF (2008) Runs of homozygosity in European populations. Am J Hum Genet 83: 359-372
Paschou P, Drineas P, Yannaki E, Razou A, Kanaki K, Tsetsos F, Padmanabhuni SS, Michalodimitrakis M, Renda MC, Pavlovic S, Anagnostopoulos A, Stamatoyannopoulos JA, Kidd KK, Stamatoyannopoulos G (2014) Maritime route of colonization of Europe. Proc Natl Acad Sci USA 111: 9211-9216
Patterson N, Price AL, Reich D (2006) Population structure and eigenanalysis. PLoS Genet 2: e190
Purcell S, Neale B, Todd-Brown K, Thomas L, Ferreira MAR, Bender D, Maller J, Sklar P, de Bakker PIW, Daly MJ, Sham PC (2007) PLINK: A tooset for whole-genome association and population-based linkage analyses. Am J Hum Genet 81: 559-575
Seidman DN, Shenoy SA, Kim M, Babu R, Woods IG, Dyer TD, Lehman DM, Curran JE, Duggirala R, Blangero J, Williams AL (2020) Rapid, phase-free detection of long Identity-by-Descent segments enables effective relationship classification. Am J Hum Genet 106: 453-466

#### Supplemental Figures

S1. PCA analysis (500K dataset)

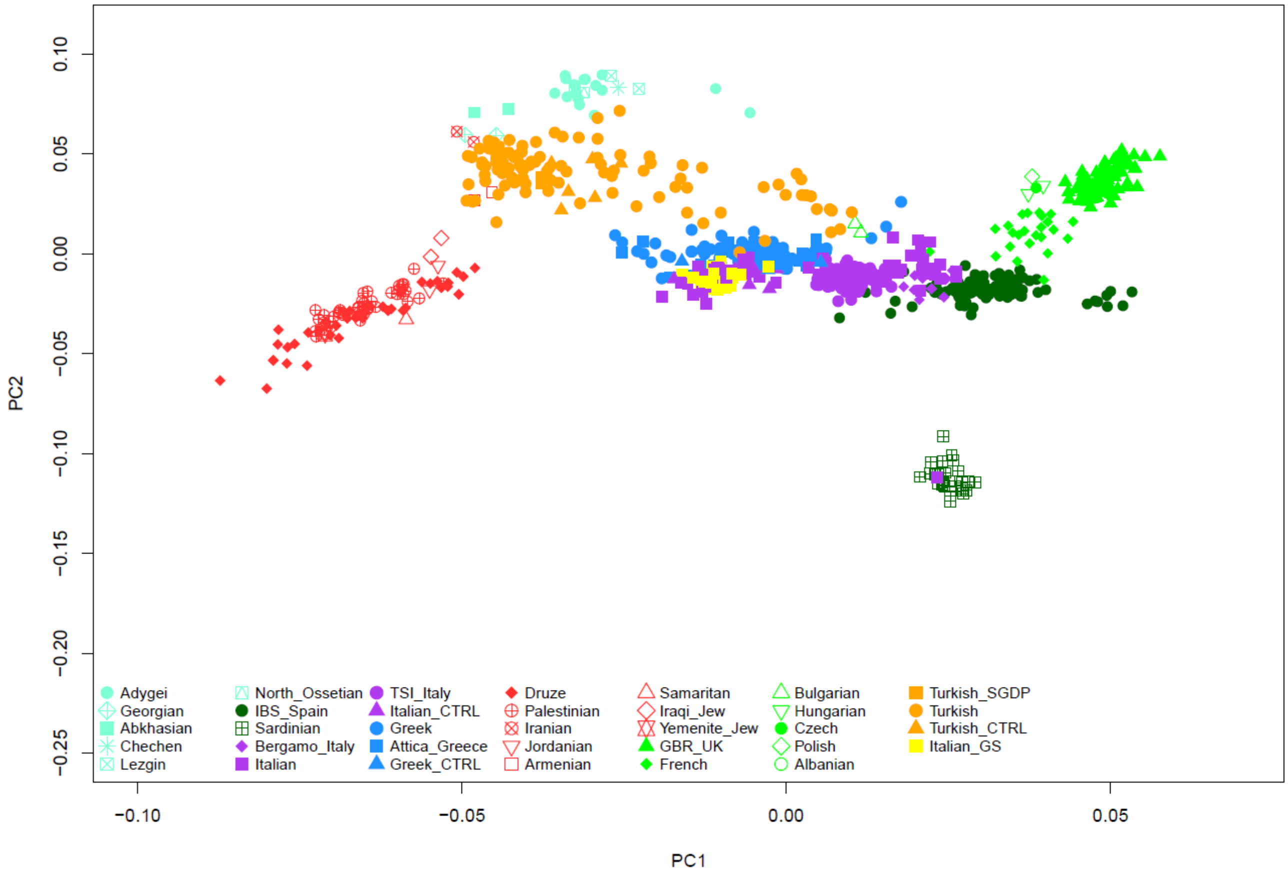


Plot of the 799 subjects of the 500K dataset in the space of PC1 vs. PC2. The plots of PC1 vs the following PCs are reported as supplemental figures in a separate file.

S2. PCA analysis (500K dataset)

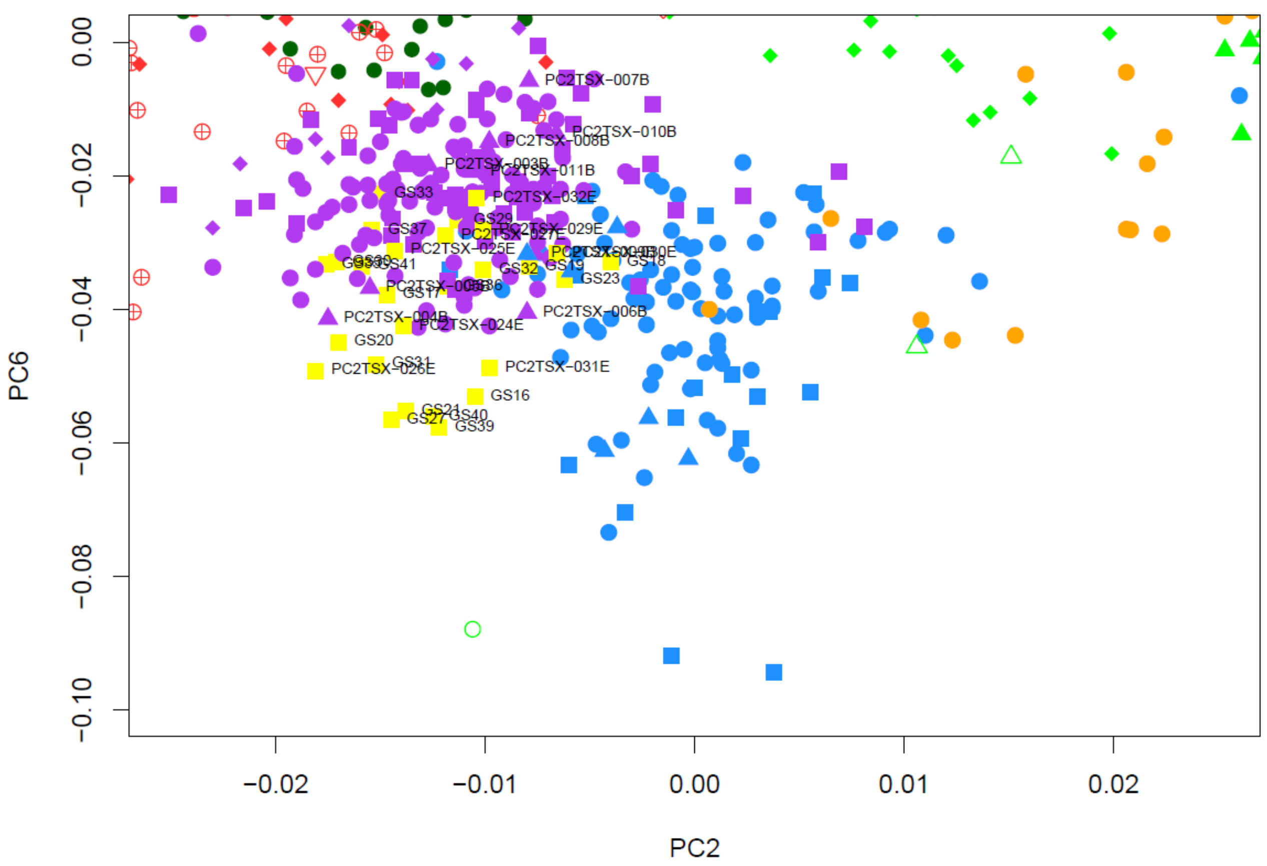


Plot of the 27 GS subjects (yellow squares), Italian (purple symbols) and Greek samples (light blue symbols) in the space of PC2 vs. PC6 as determined on the 500K dataset. Other subjects fall outside the displayed area. All symbols as in Fig. S2.

S3. Expected sampling locations based on PCA analysis (500K dataset)

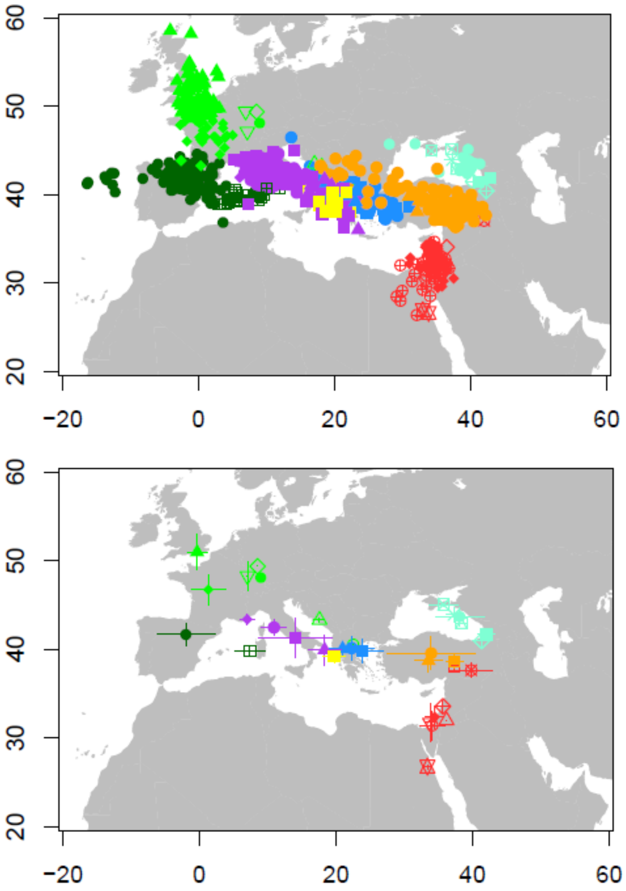


Plot of the 799 samples (top) of the 500K dataset on their “Expected Sampling Location” (see Text for definition). Symbols and colours are as in the original PCA plot (Fig. S2). “Expected Sampling Location” averages for the 34 population_ID’s (bottom). Whiskers represent +/- 1 s.d.

#### S4. ADMIXTURE analysis (500K dataset)

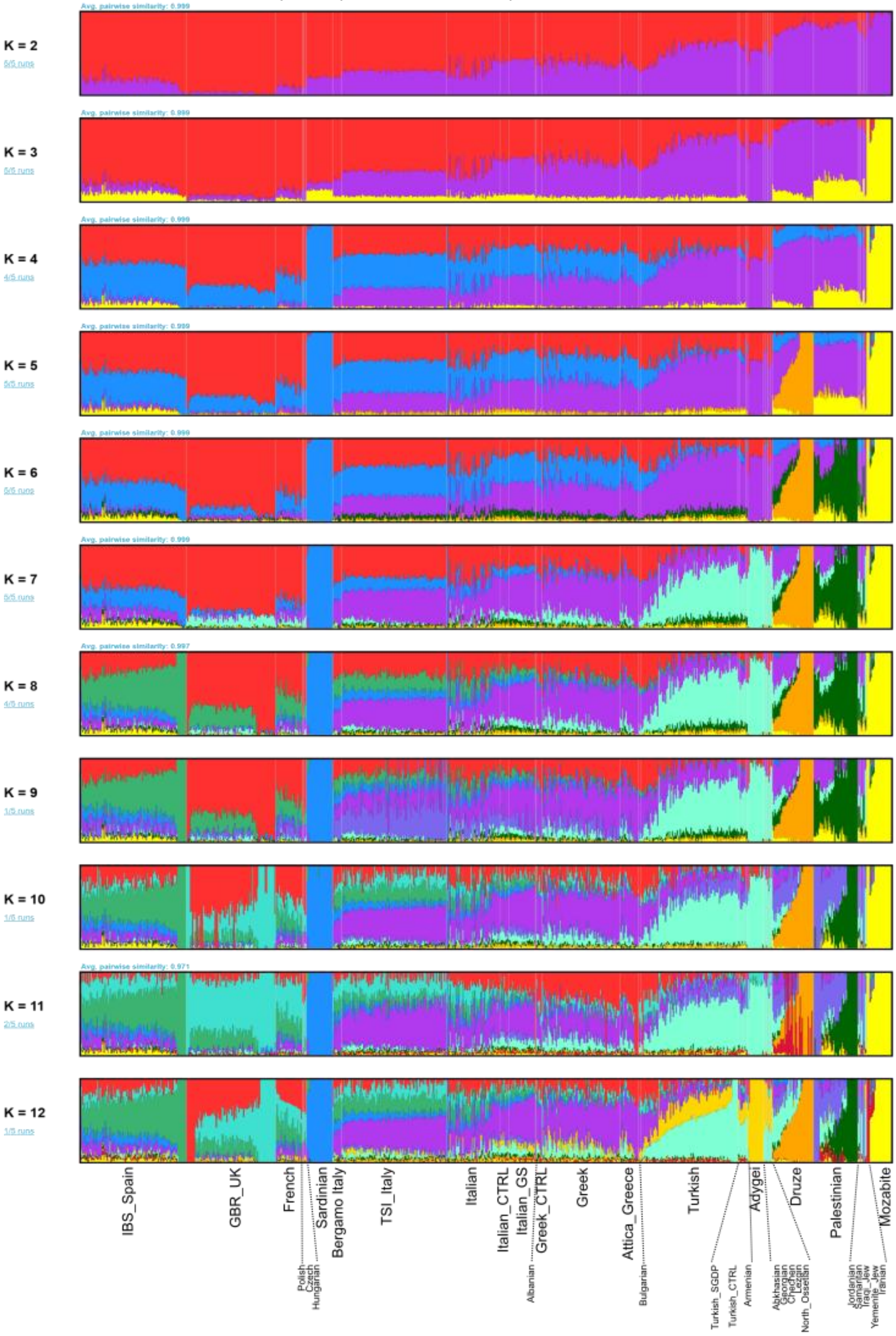


Admixture analysis of the 500K dataset for K 2 to 12. The samples included in each vertical belt are shown at bottom. Each plot is based on 5 runs, summarized with pong. The number of runs producing coherent partitions in K components is reported in light blue to the left.

#### S5. PCA analysis 1K dataset

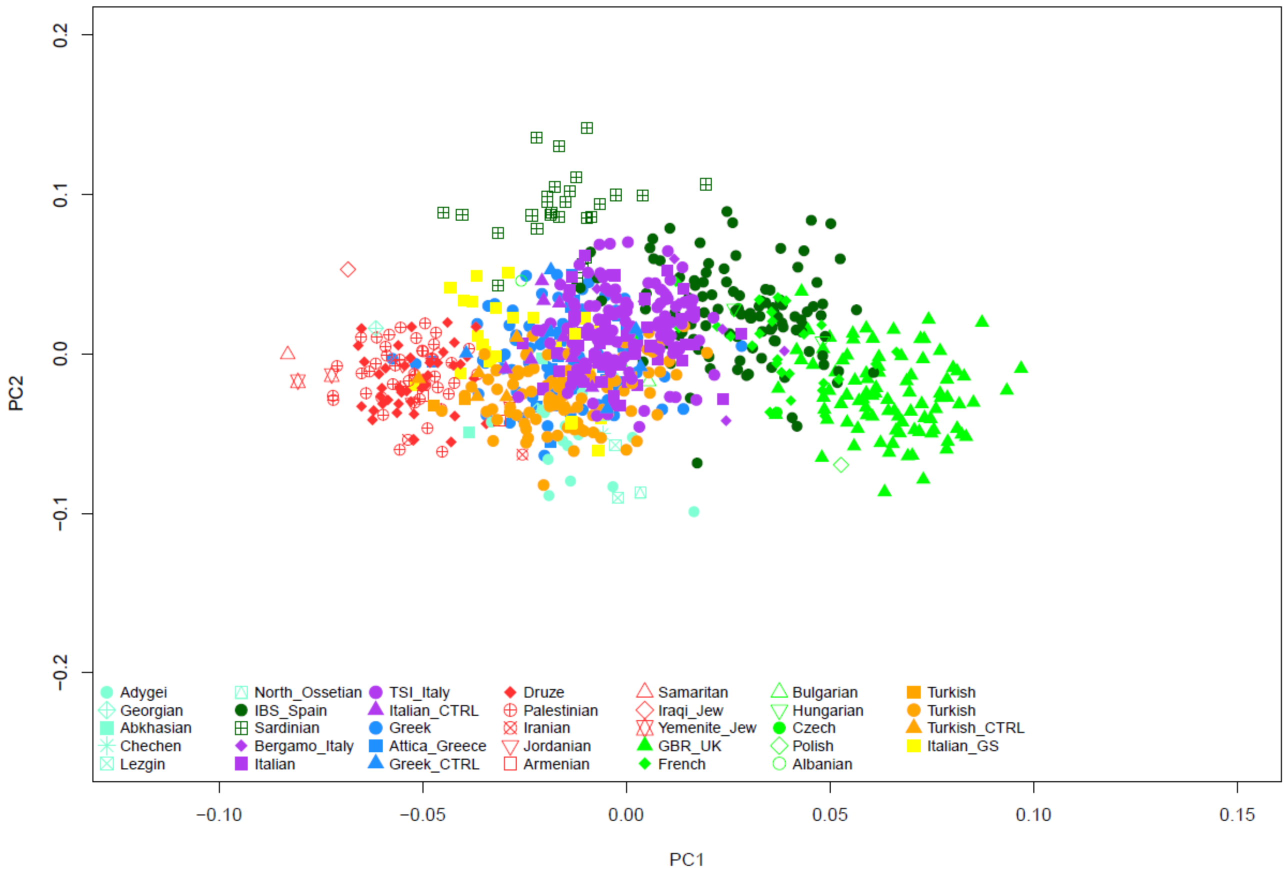


Plot of the 799 samples genotyped at 997 AIMs for the European populations (Drineas et al. 2010) in the space of PC1 vs. PC2.

#### S6. Expected sampling locations based on PCA analysis (1K dataset)

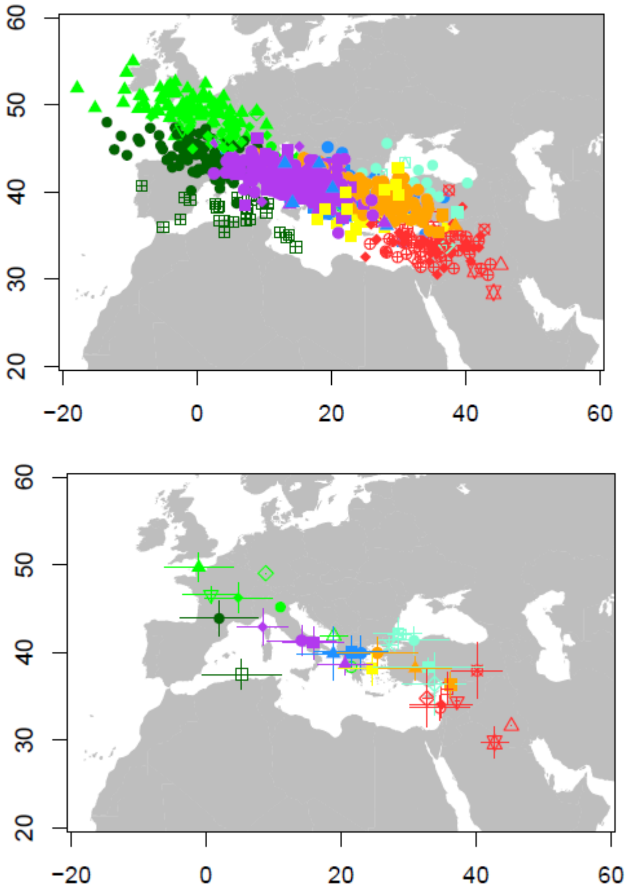


Plot of the 799 samples (top) of the 1K dataset on their “Expected Sampling Location” (see Text for definition). Symbols and colours are as in the original PCA plot (Fig. S5). “Expected Sampling Location” averages for the 34 population samples (bottom). Whiskers represent +/- 1 s.d


Drineas P, Lewis J, Paschou P (2010) Inferring geographic coordinates of origin for Europeans using small panels of ancestry informative markers. PLoS ONE 5: e11892

S7. PCA analysis 25K dataset

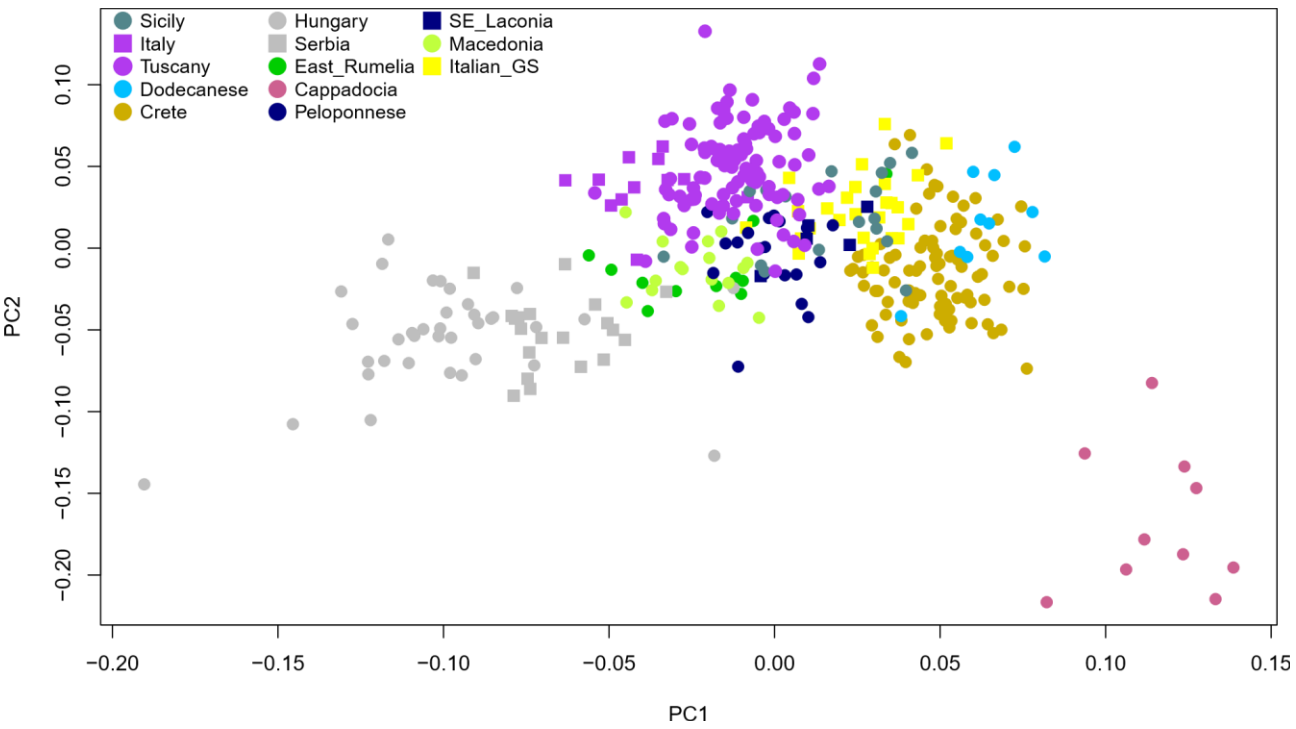


Plot in the space of PC1 and PC2 of 344 samples from the Balkan peninsula, Italy, Crete and Türkiye typed by (Paschou et al. 2014), plus the 27 GS subjects

Paschou P, Drineas P, Yannaki E, Razou A, Kanaki K, Tsetsos F, Padmanabhuni SS, Michalodimitrakis M, Renda MC, Pavlovic S, Anagnostopoulos A, Stamatoyannopoulos JA, Kidd KK, Stamatoyannopoulos G (2014) Maritime route of colonization of Europe. Proc Natl Acad Sci USA 111: 9211-9216

S8. IBD sharing

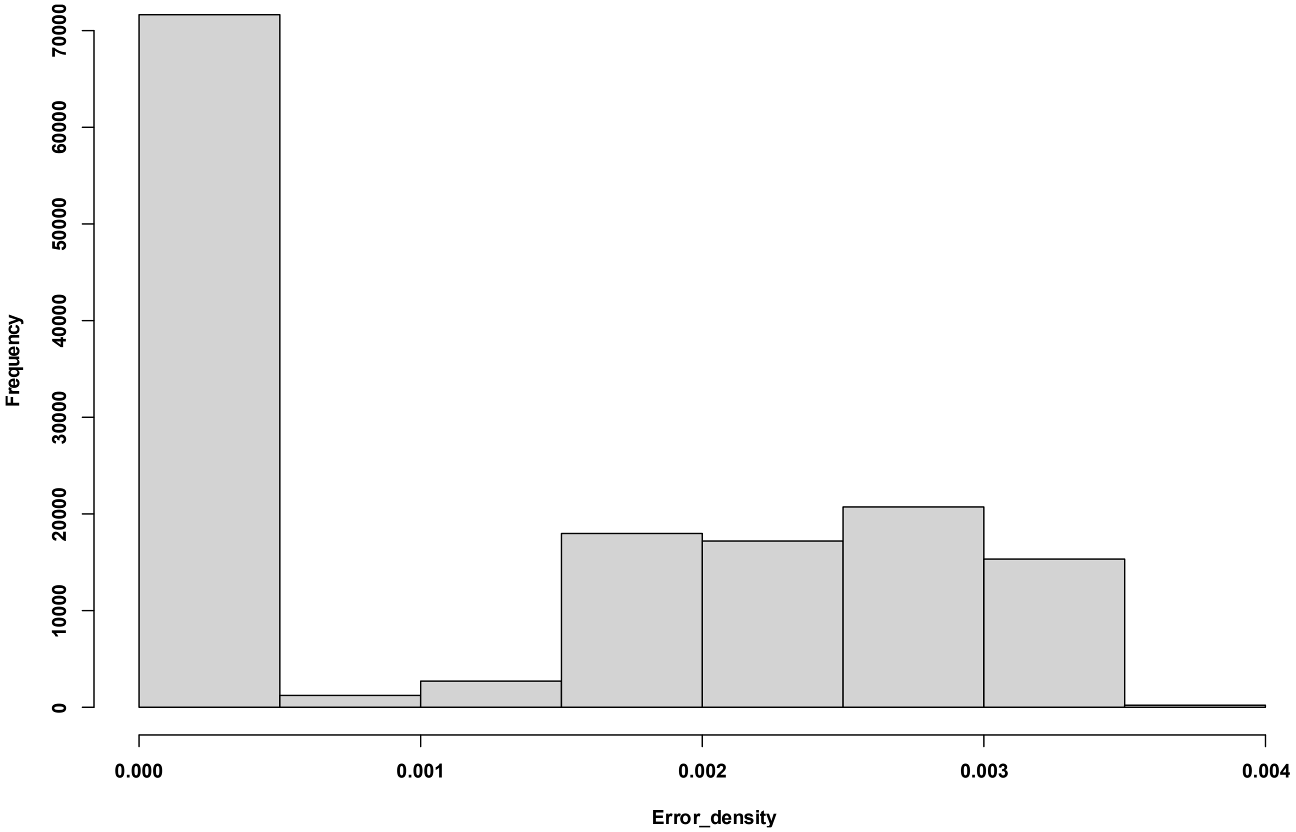


Histogram of the error density in 147,012 IBD fragments longer than 3 cM.

#### S9. Decay of IBD sharing with distance

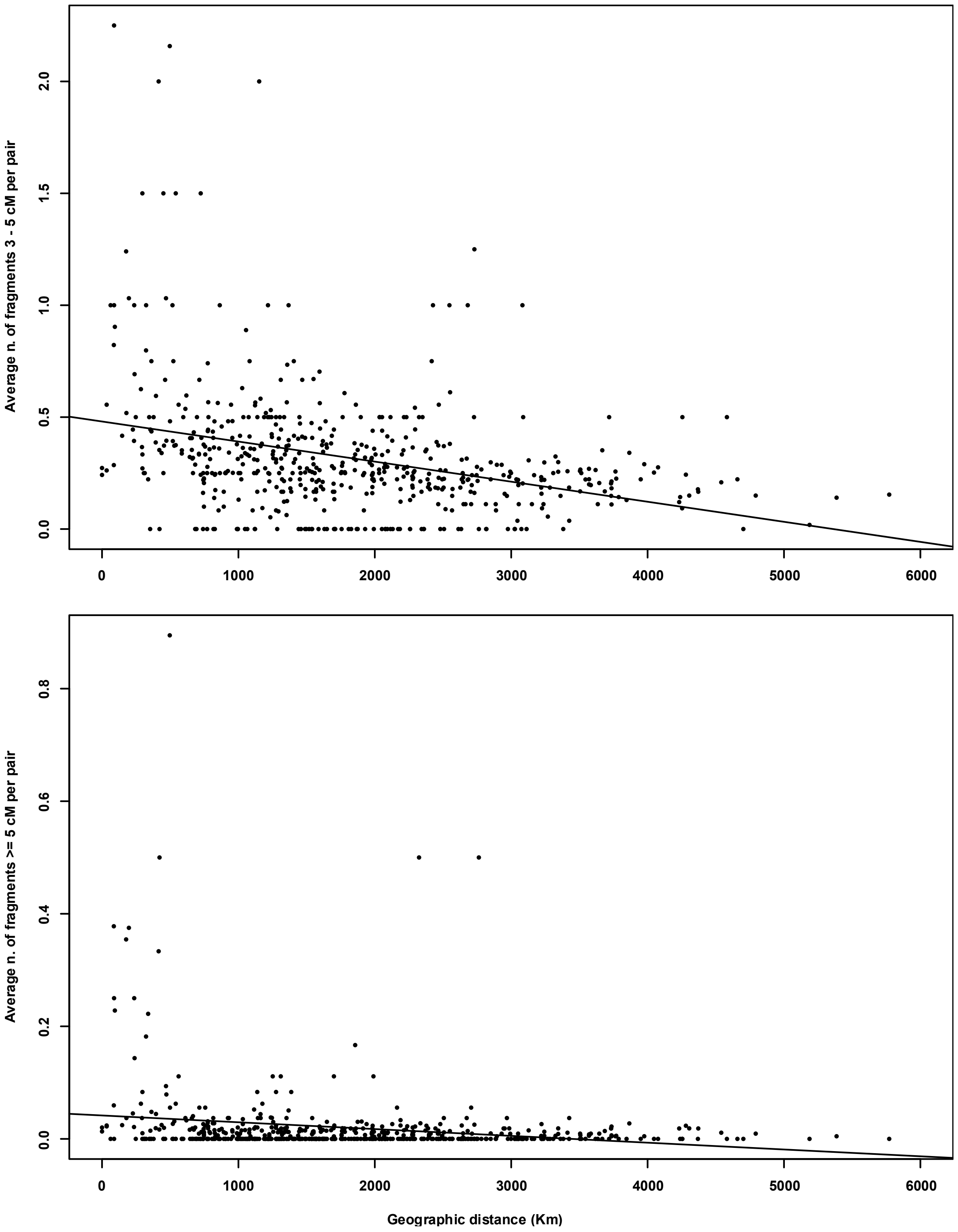


Decay of sharing of IBD fragments of length 3-5 cM (top) and ≥5 cM (bottom) with distance. Each dot represents the average sharing between subjects from different Population_IDs (comparisons between subjects of the same Population_ID are omitted). Note the different Y scale for short (above) and long (below) fragments. Compare with Figure 3 by (Ralph and Coop 2013).

S 10.

#### Average n. of shared IBD fragments

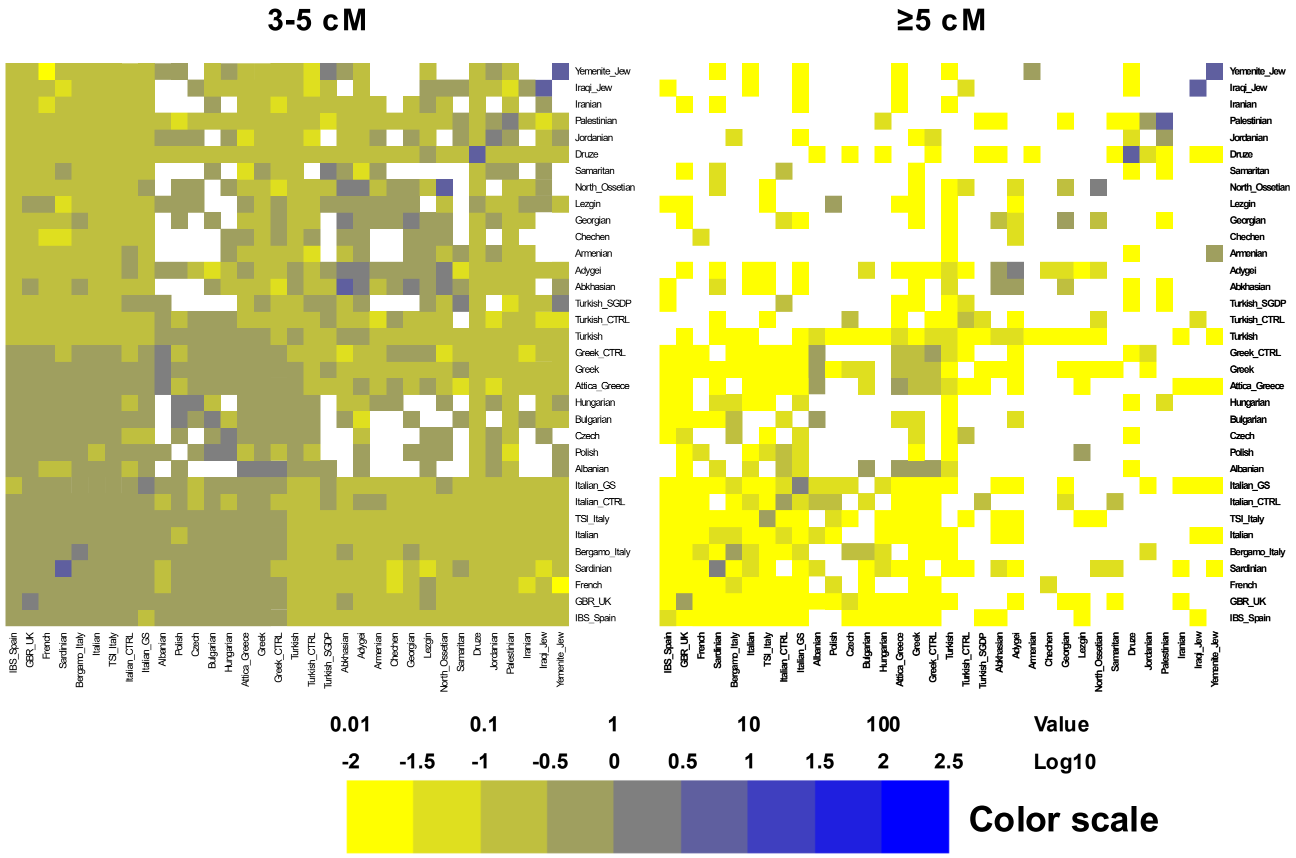


Average n. of IBD fragments shared in pairwise comparisons between subjects. The color scale is the same for the two panels. Sharing is generally higher within than between population groups.

#### S 11. IBD sharing

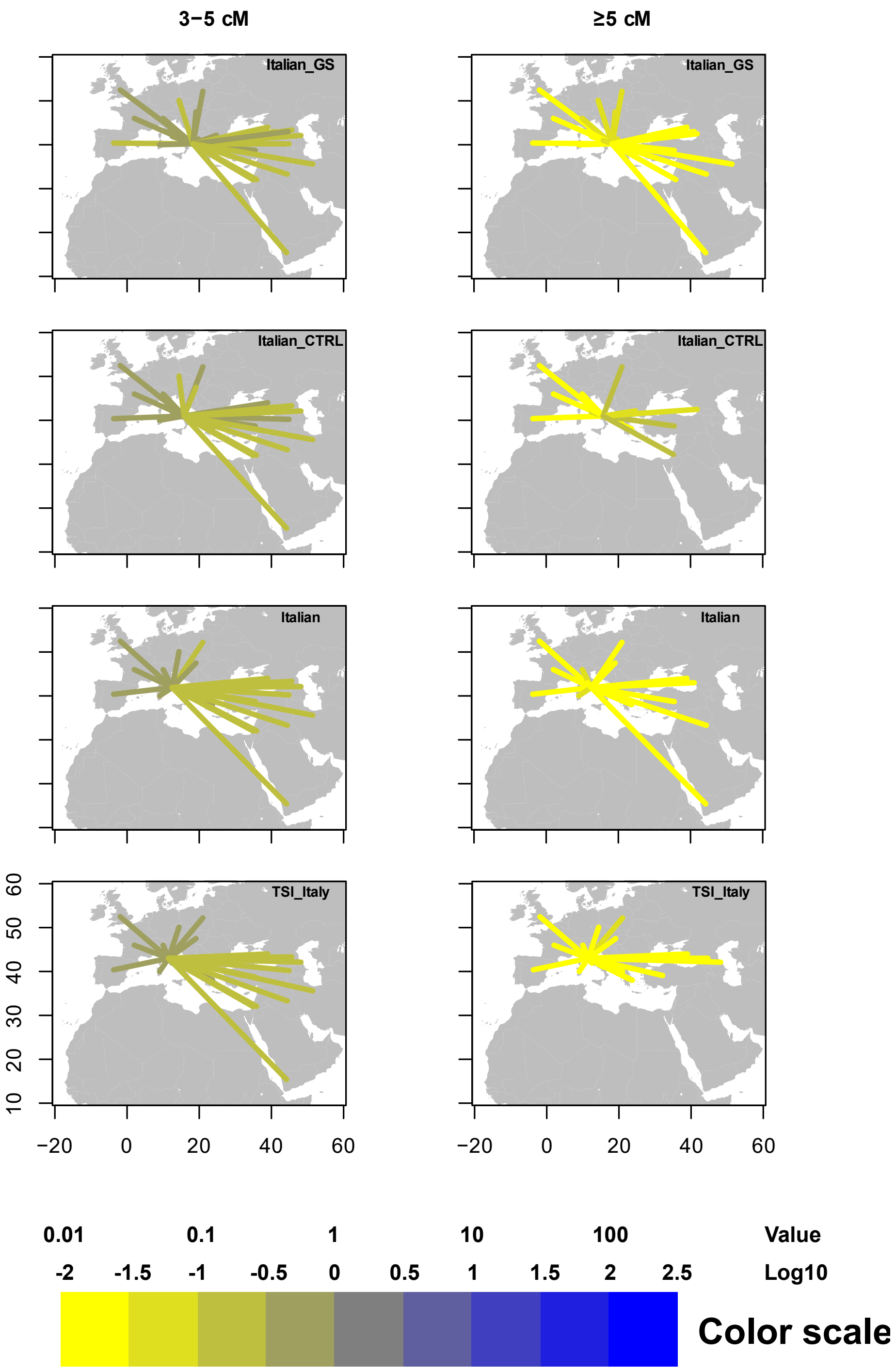

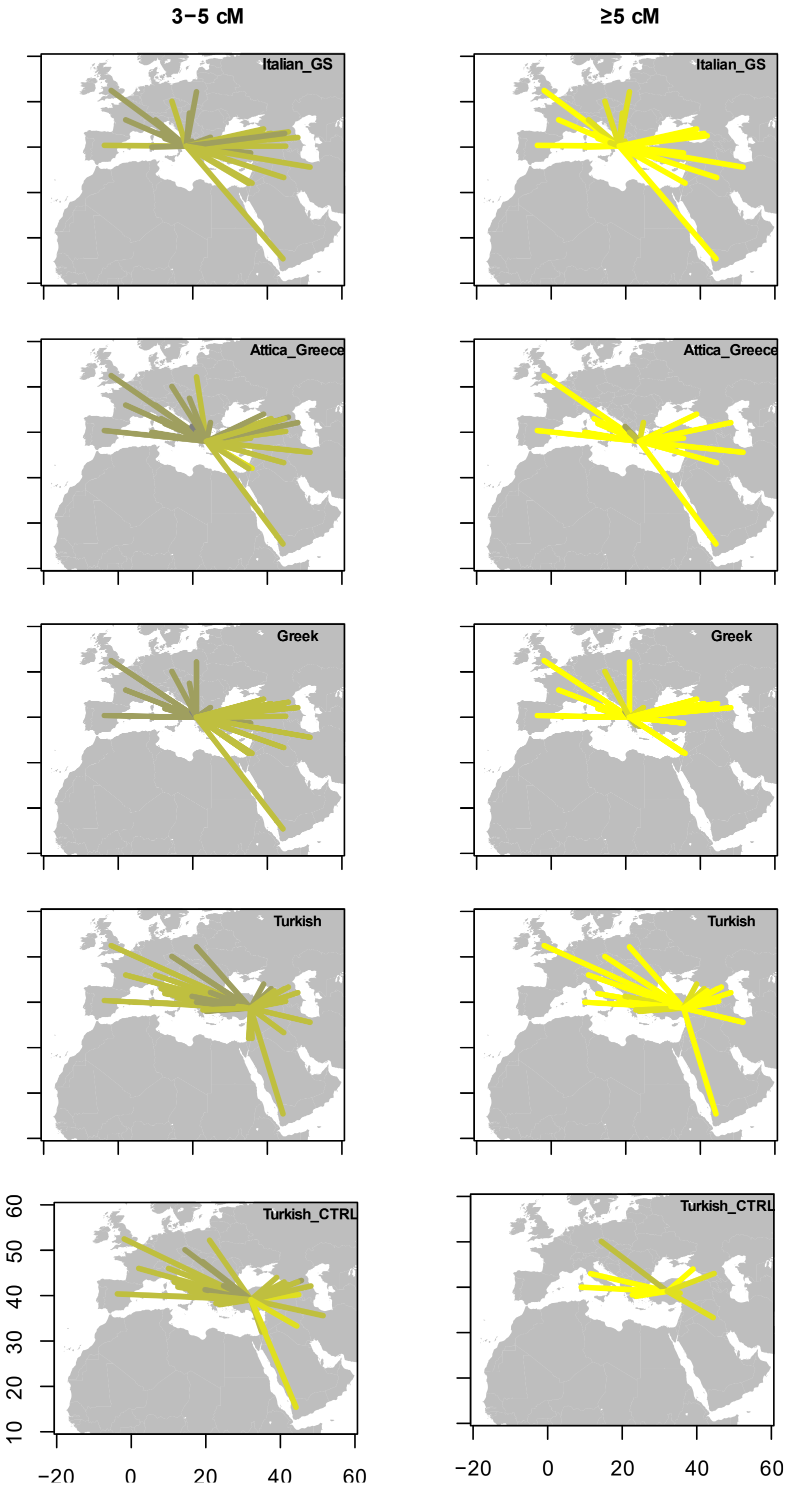


Maps comparing the sharing of IBD fragments recovered in GS subjects and Population_IDs from Italy (top) or Greece and Türkiye (bottom). Lines connect the target Population_ID and each of the others harbouring shared IBD's. Lines are coloured according to the average number of shared IBD's in pairwise comparisons (same scale across all panels).

Ralph P, Coop G (2013) The geography of recent genetic ancestry across Europe. PLoS Biol 11: e1001555

#### S 12. f3 and D statistics

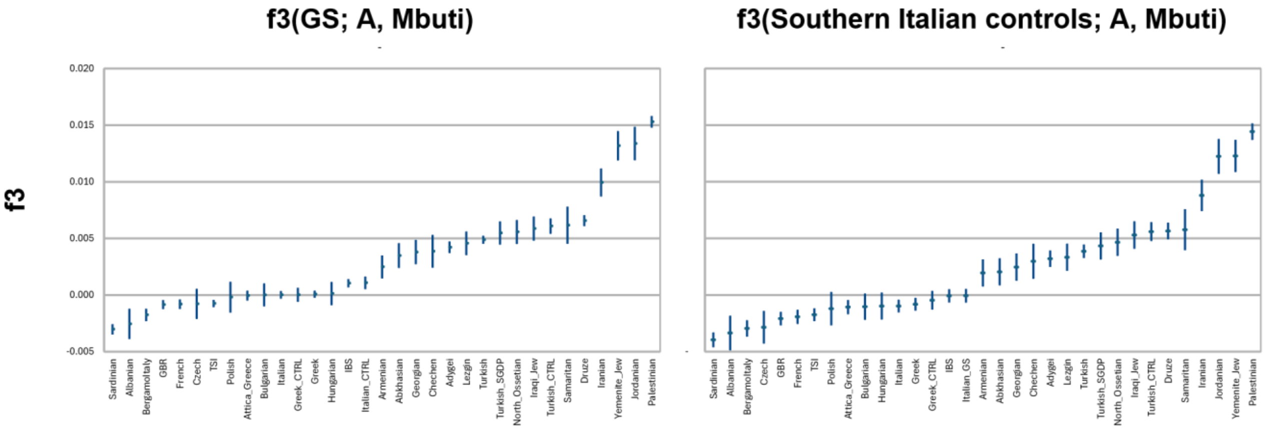


f3 statistics for GS (left) and Southern Italian controls (right) as targets for admixture between the indicated populations and the Mbuti as distant outgroup. For both GS and the Southern Italian controls the order of populations producing f3 values from smallest to largest and the absolute f3 values are highly similar, indicating no gross differences between GS and Southern Italians in their ancestry relationships with other populations. Whiskers indicate +/- 1.96 s.e.

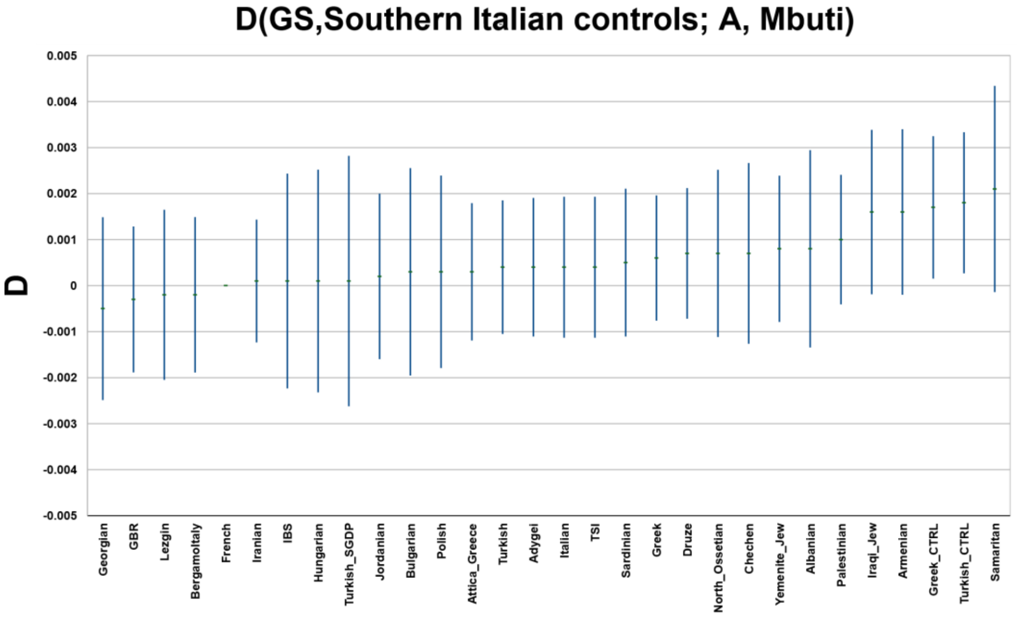


D statistics in the form reported above, to test for the excess shared ancestry between GS and the indicated populations as compared to Southern Italian controls. Note the two positive significant values for Greek and Turkish controls. Whiskers indicate +/- 1.96 s.e

.

#### S13. Sharing of recent dated alleles

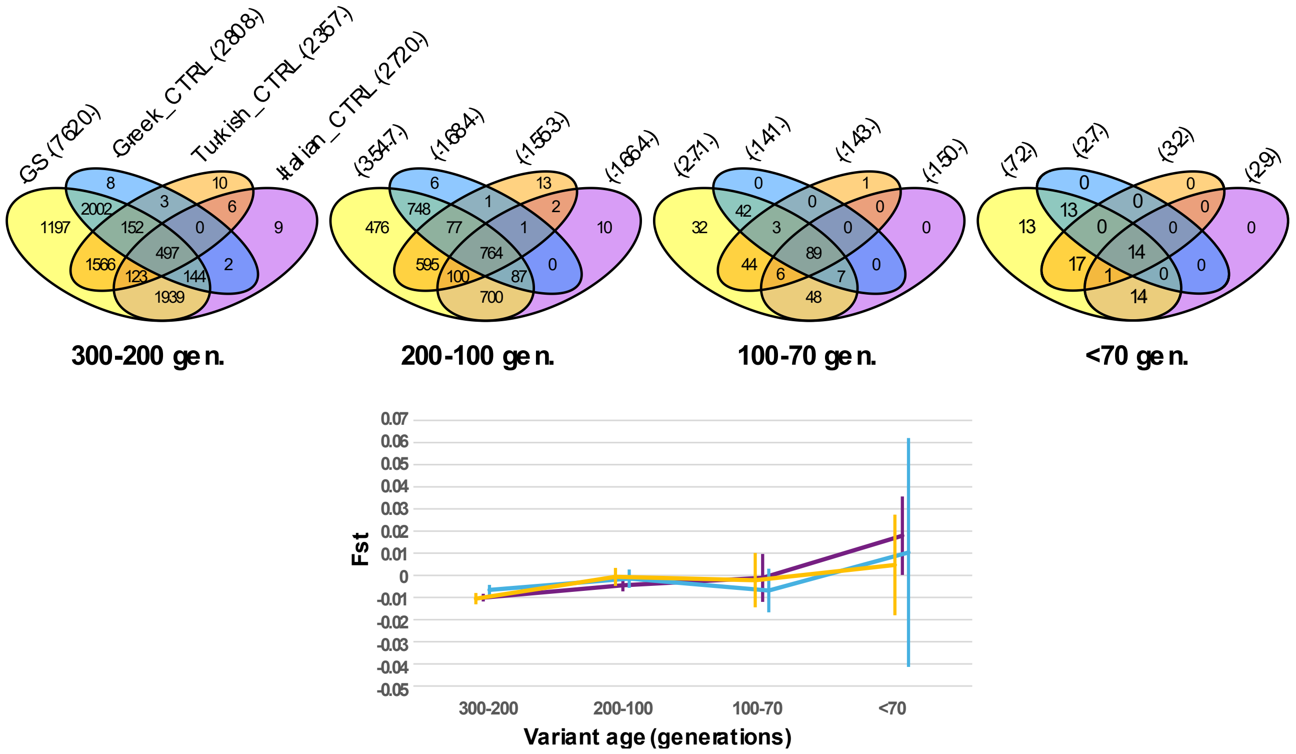


Sharing of derived alleles at positions with estimated age as indicated (top). The Italian samples were downsampled to 6, so that the Venn diagrams are based on 27, 6, 6 and 6 subjects for GS, Greek, Turkish and Italian controls, respectively, and occurrences within cells are thus directly comparable. Weir and Cockeram’s Fst and 95% c.i. for comparisons between GS and Italian controls (not downsampled, deep blue), Greek controls (light blue) and Turkish controls (orange)

#### S14. Allele frequency spectra of dated variants

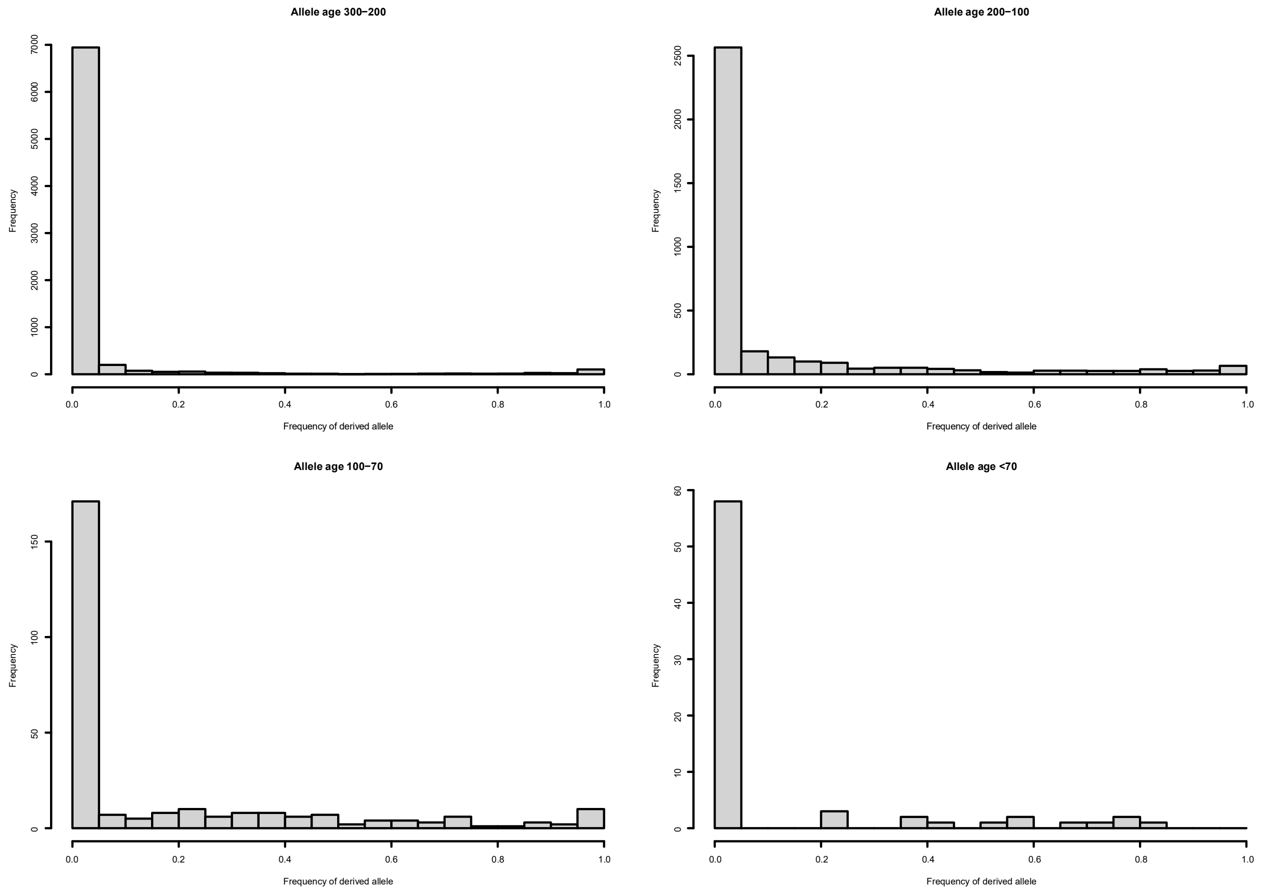


Allele frequency spectra of variants in the indicated age ranges (note the different scales on the y axes).

#### S15. GS in the context of aDNA samples

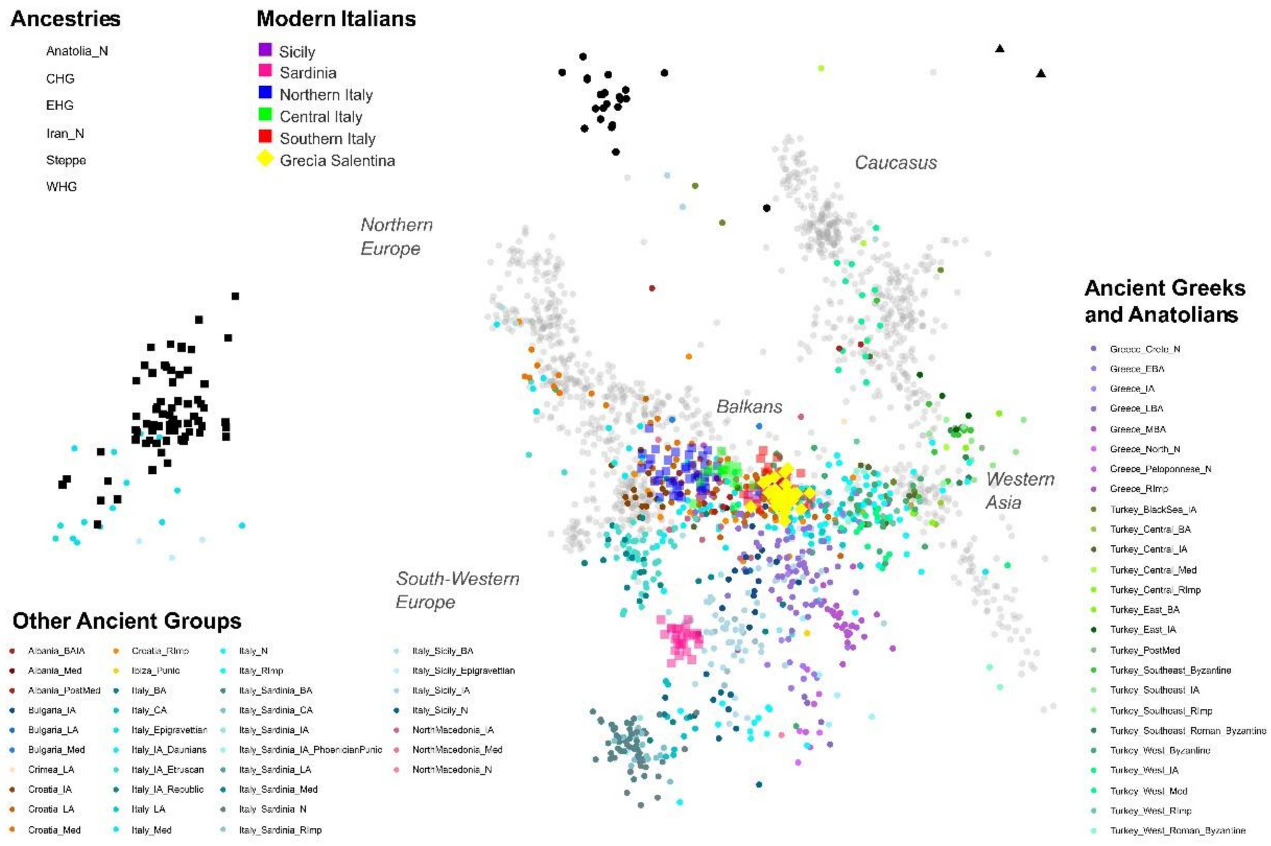


Principal component analysis (PCA) of 27 individuals from Grecìa Salentina (yellow diamonds), 1,275 modern Eurasians (grey dots and coloured squares), and 2,746 ancient genomes (coloured dots and black symbols). Principal components were computed using the modern dataset (including GS), and the ancient genomes were subsequently projected onto the resulting PCA space. The first part of each population label refers to the present-day political entity in which the sampling location is situated. Abbreviations: CHG = Caucasus Hunter-Gatherer; EHG = Eastern Hunter-Gatherer; WHG = Western Hunter-Gatherer; N = Neolithic; CA = Copper Age; EBA = Early Bronze Age; MBA = Middle Bronze Age; LBA = Late Bronze Age; IA = Iron Age; RImp = Roman Imperial; LA = Late Antiquity; Med = Medieval; PostMed = Post-Medieval.

### S16. Connectivity within GS

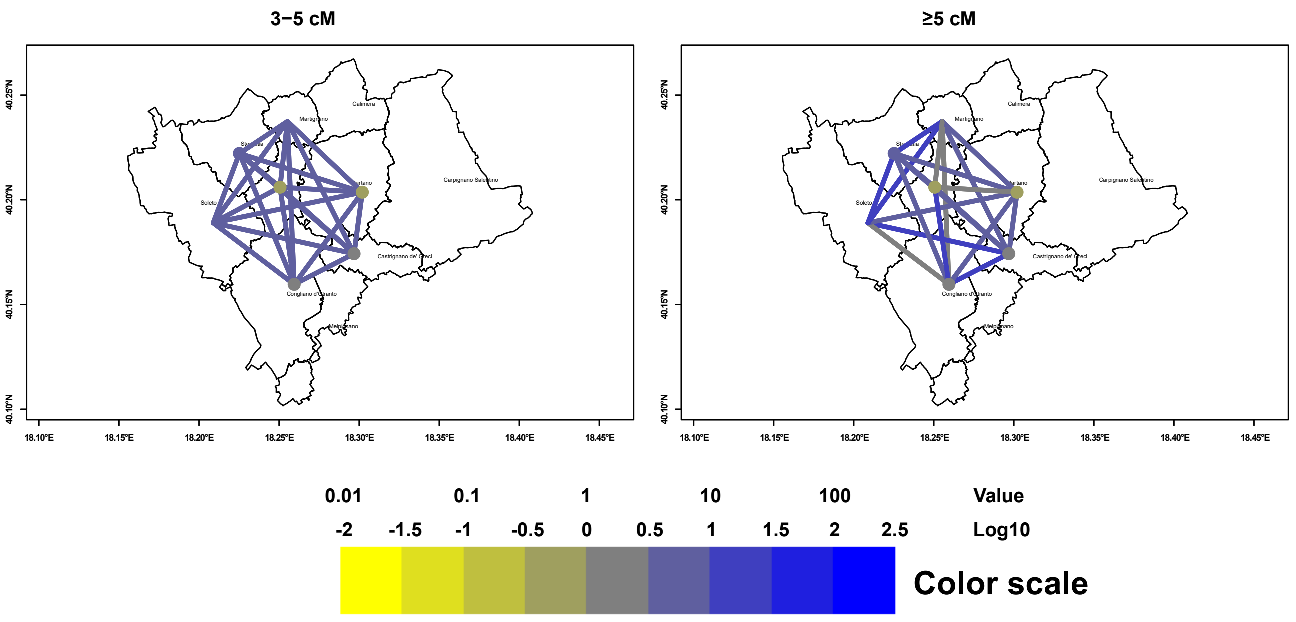


Average n. of IBD fragments shared in pairwise comparisons between subjects from different (lines) or the same (dots) municipality in GS.

S17. Runs of homozygosity

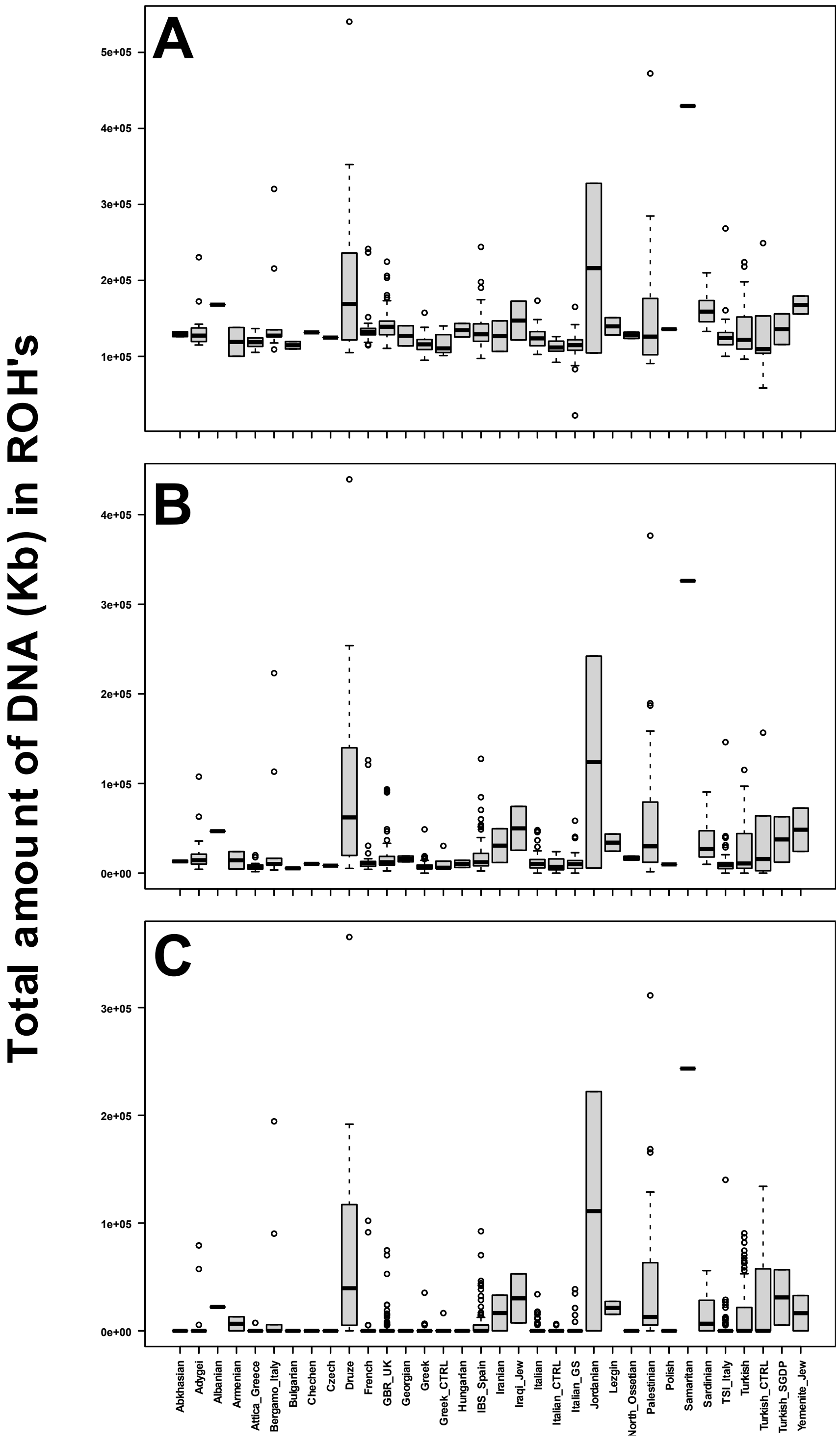


Boxplot of amount of DNA (in Kb) represented in ROH's longer than given thresholds in the autosomes of the 799 subjects of the 500 Kb dataset (Table S2) by population_ID. ROH's longer than 500 Kb (A), 1,500 Kb (B) and 5,000 Kb (C). Black lines indicate the medians, grey boxes the inter-quartile range (IQR), whiskers extend to +/- 1.5IQR and outliers are shown as individual points.
