## Supplementary material for "LAYERED GENOMIC COMPOSITION AND LINGUISTIC CONTINUITY IN GRECÌA SALENTINA": PCA PLOTS 1-20

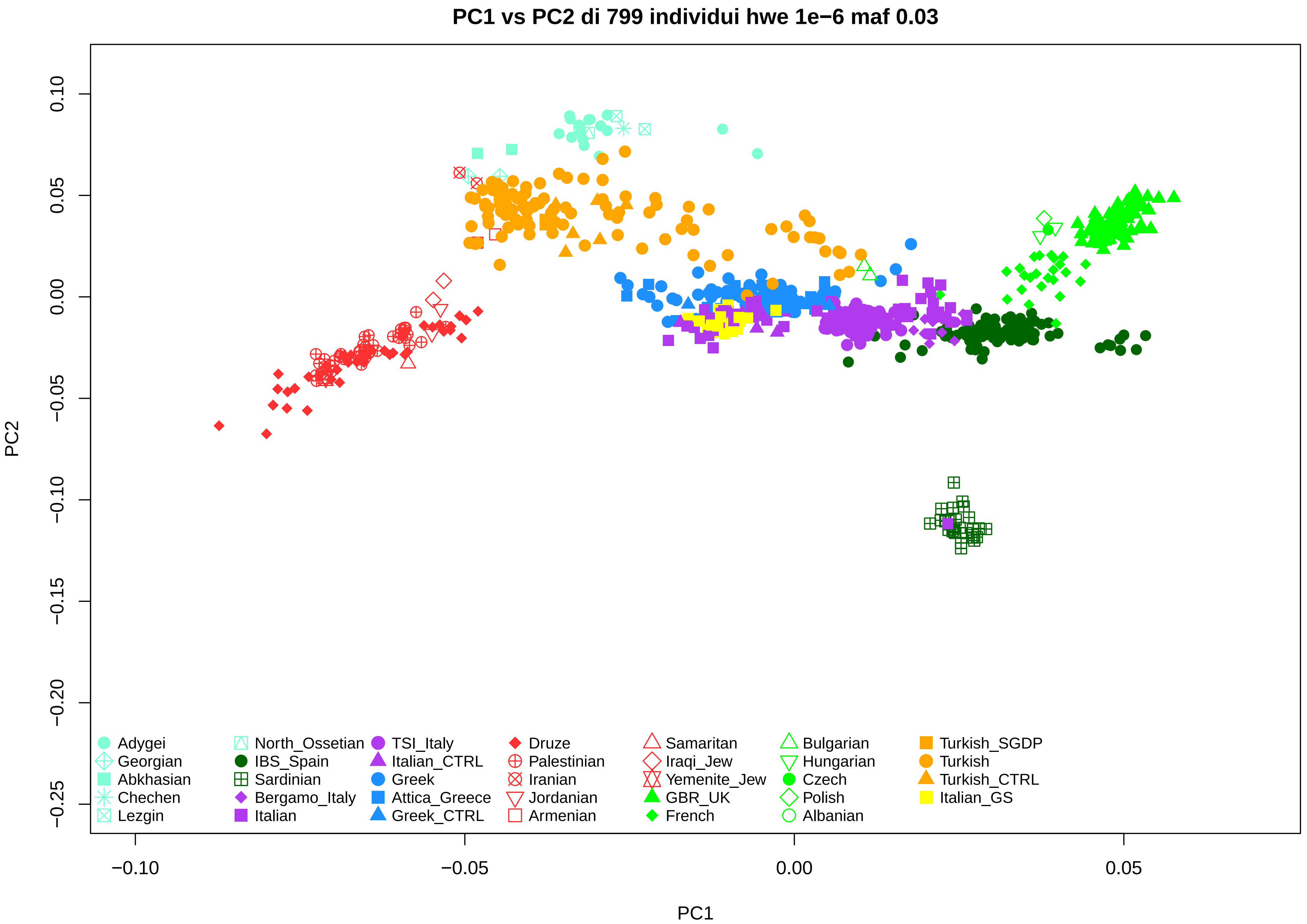
PC1 vs PC2 di 799 individui hwe 1e−6 maf 0.03
PC2
PC1
0.10
0.05
0.00
−0.05
−0.10
−0.15
−0.20
−0.25
−0.10
−0.05
0.00
0.05
Adygei
Georgian
Abkhasian
Chechen
Lezgin
North_Ossetian
IBS_Spain
Sardinian
Bergamo_Italy
Italian
TSI_Italy
Italian_CTRL
Greek
Attica_Greece
Greek_CTRL
Druze
Palestinian
Iranian
Jordanian
Armenian
Samaritan
Iraqi_Jew
Yemenite_Jew
GBR_UK
French
Bulgarian
Hungarian
Czech
Polish
Albanian
Turkish_SGDP
Turkish
Turkish_CTRL
Italian_GS

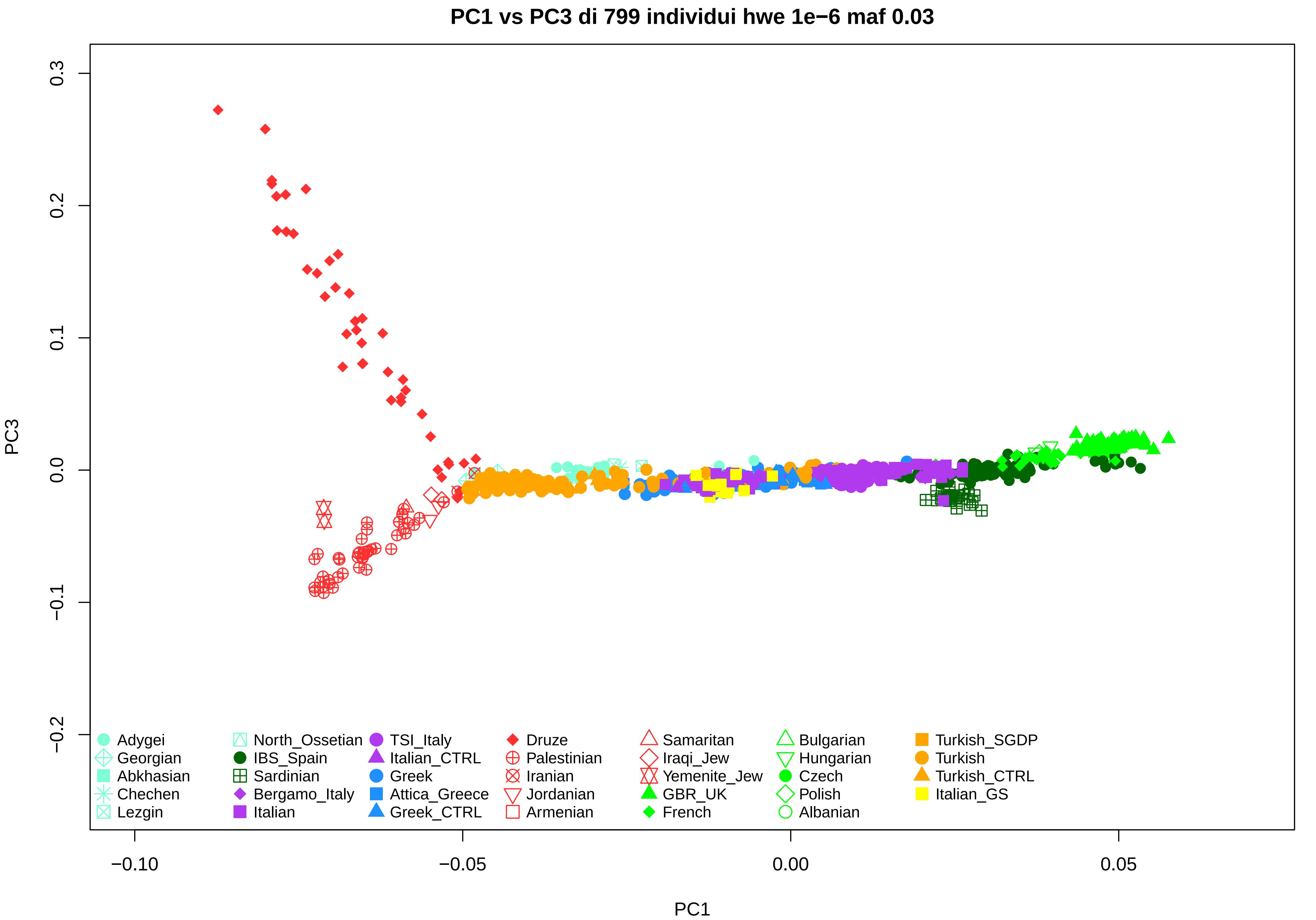
PC1 vs PC3 di 799 individui hwe 1e−6 maf 0.03
PC3
PC1
0.3
0.2
0.1
0.0
−0.1
−0.2
−0.10
−0.05
0.00
0.05
Adygei
Georgian
Abkhasian
Chechen
Lezgin
North_Ossetian
IBS_Spain
Sardinian
Bergamo_Italy
Italian
TSI_Italy
Italian_CTRL
Greek
Attica_Greece
Greek_CTRL
Druze
Palestinian
Iranian
Jordanian
Armenian
Samaritan
Iraqi_Jew
Yemenite_Jew
GBR_UK
French
Bulgarian
Hungarian
Czech
Polish
Albanian
Turkish_SGDP
Turkish
Turkish_CTRL
Italian_GS

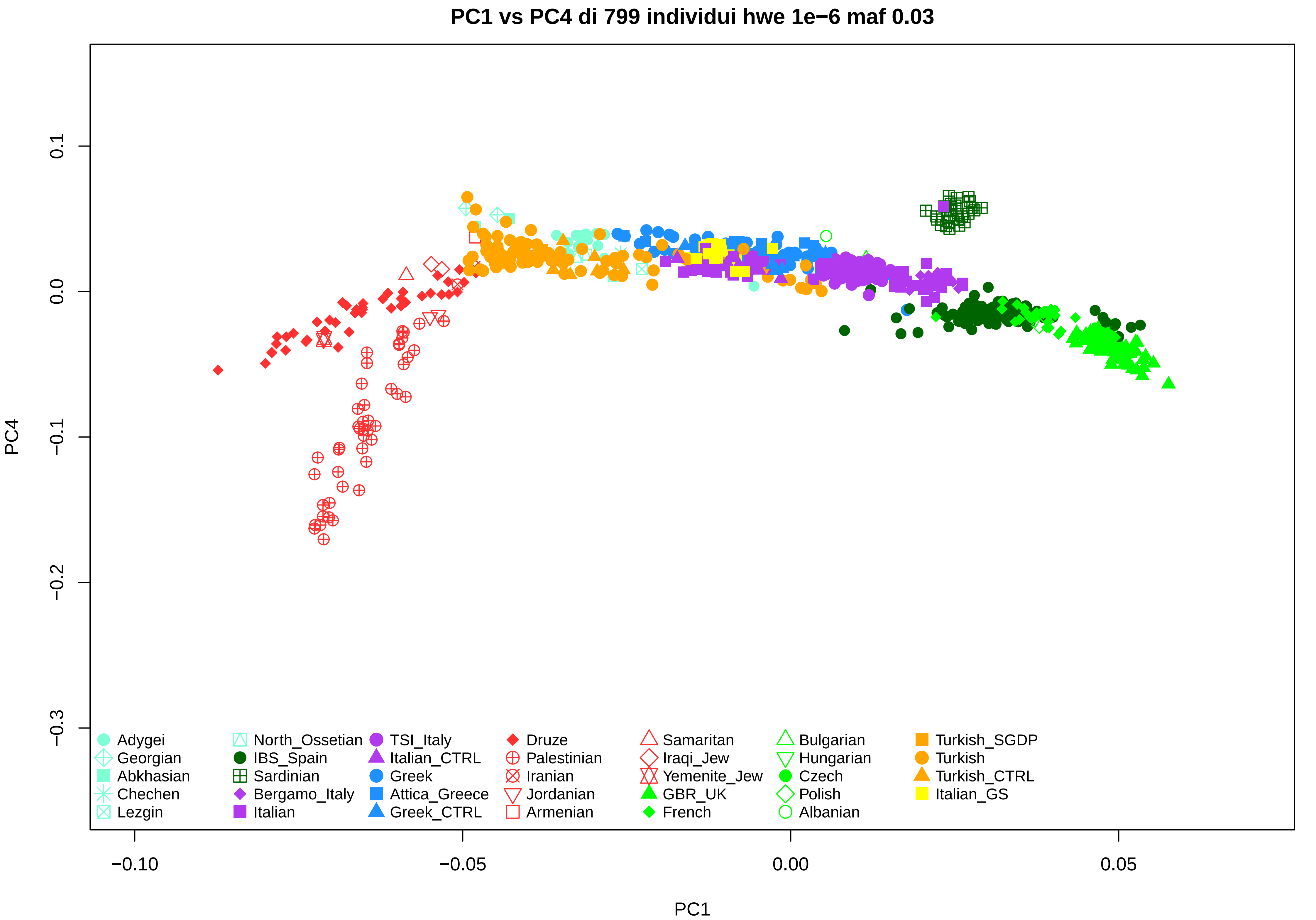
PC1 vs PC4 di 799 individui hwe 1e−6 maf 0.03
PC4
PC1
0.1
0.0
−0.1
−0.2
−0.3
−0.10
−0.05
0.00
0.05
Adygei
Georgian
Abkhasian
Chechen
Lezgin
North_Ossetian
IBS_Spain
Sardinian
Bergamo_Italy
Italian
TSI_Italy
Italian_CTRL
Greek
Attica_Greece
Greek_CTRL
Druze
Palestinian
Iranian
Jordanian
Armenian
Samaritan
Iraqi_Jew
Yemenite_Jew
GBR_UK
French
Bulgarian
Hungarian
Czech
Polish
Albanian
Turkish_SGDP
Turkish
Turkish_CTRL
Italian_GS

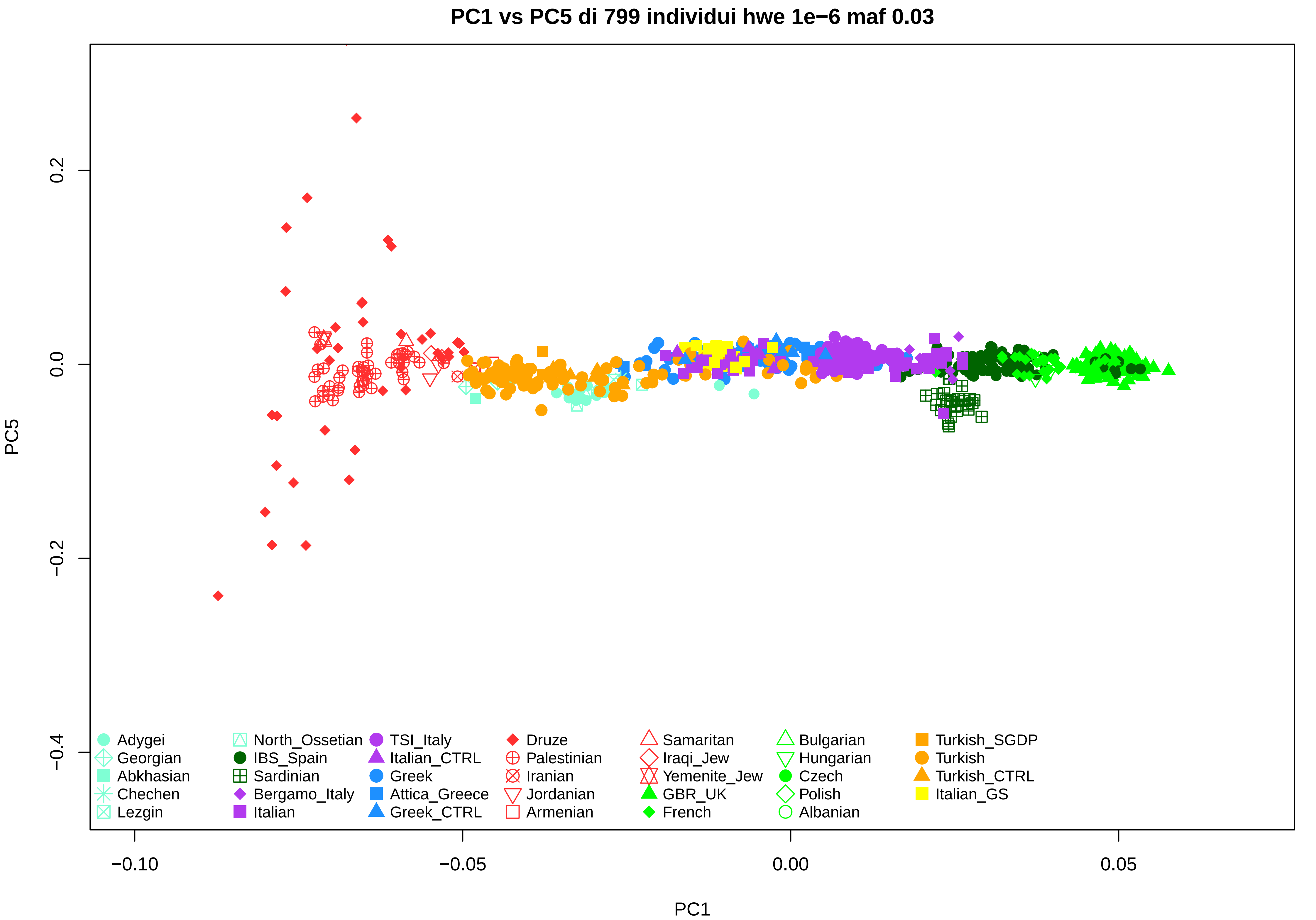
PC1 vs PC5 di 799 individui hwe 1e−6 maf 0.03
PC5
PC1
0.2
0.0
−0.2
−0.4
−0.10
−0.05
0.00
0.05
Adygei
Georgian
Abkhasian
Chechen
Lezgin
North_Ossetian
IBS_Spain
Sardinian
Bergamo_Italy
Italian
TSI_Italy
Italian_CTRL
Greek
Attica_Greece
Greek_CTRL
Druze
Palestinian
Iranian
Jordanian
Armenian
Samaritan
Iraqi_Jew
Yemenite_Jew
GBR_UK
French
Bulgarian
Hungarian
Czech
Polish
Albanian
Turkish_SGDP
Turkish
Turkish_CTRL
Italian_GS

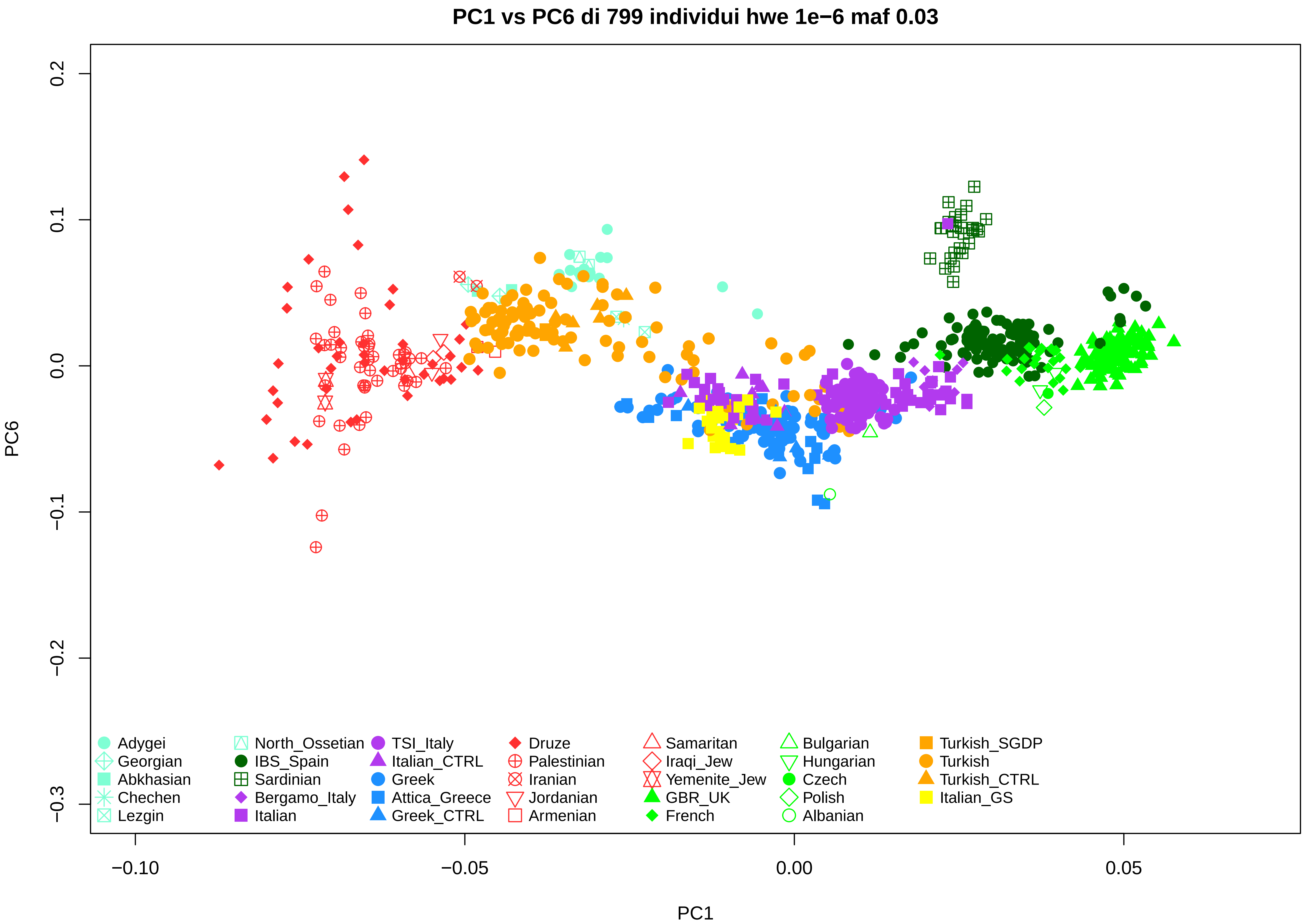
PC1 vs PC6 di 799 individui hwe 1e−6 maf 0.03
PC6
PC1
0.2
0.1
0.0
−0.1
−0.2
−0.3
−0.10
−0.05
0.00
0.05
Adygei
Georgian
Abkhasian
Chechen
Lezgin
North_Ossetian
IBS_Spain
Sardinian
Bergamo_Italy
Italian
TSI_Italy
Italian_CTRL
Greek
Attica_Greece
Greek_CTRL
Druze
Palestinian
Iranian
Jordanian
Armenian
Samaritan
Iraqi_Jew
Yemenite_Jew
GBR_UK
French
Bulgarian
Hungarian
Czech
Polish
Albanian
Turkish_SGDP
Turkish
Turkish_CTRL
Italian_GS

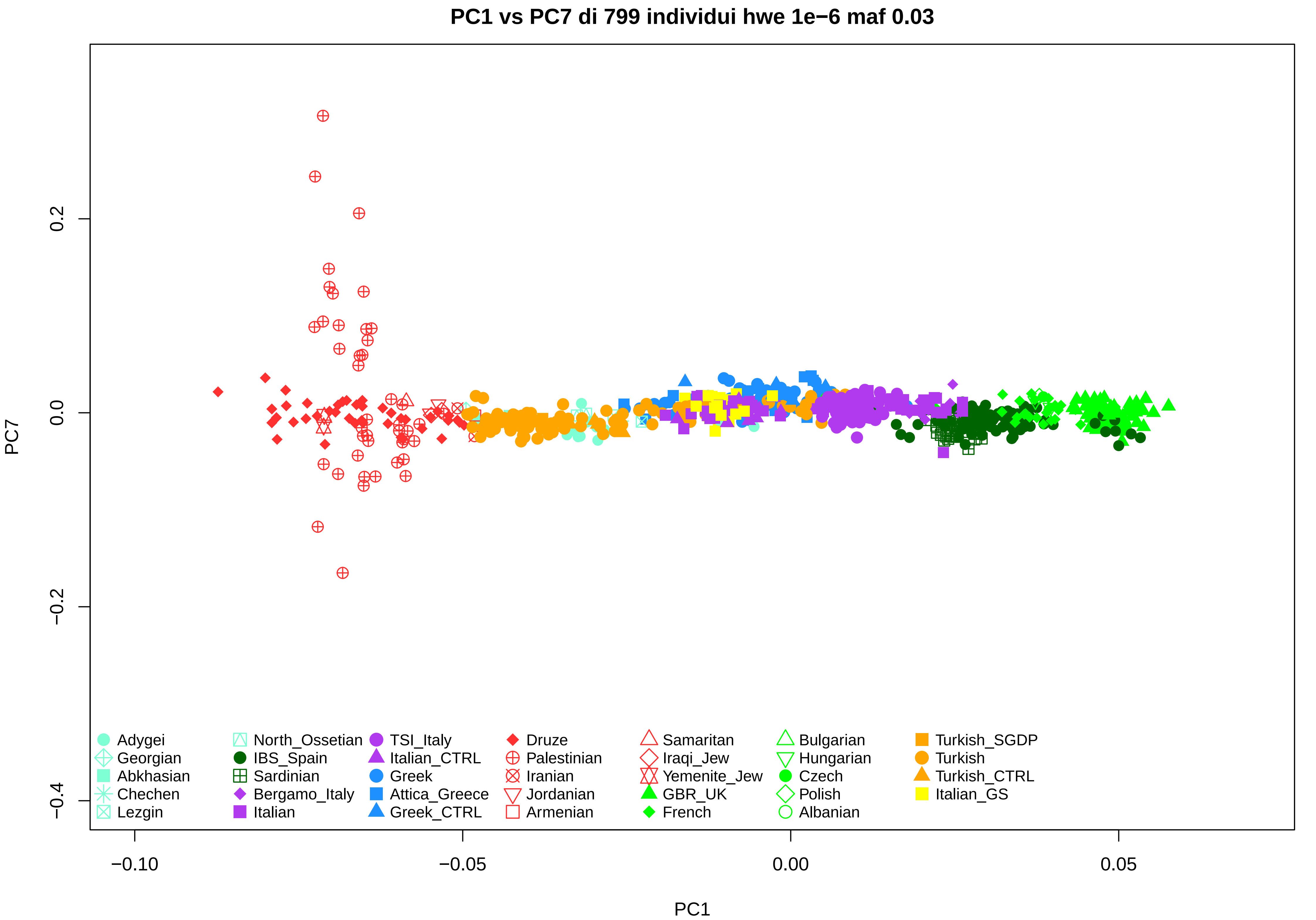

PC1 vs PC7 di 799 individui hwe 1e−6 maf 0.03
PC7
PC1
0.2
0.0
−0.2
−0.4
−0.10
−0.05
0.00
0.05
Adygei
Georgian
Abkhasian
Chechen
Lezgin
North_Ossetian
IBS_Spain
Sardinian
Bergamo_Italy
Italian
TSI_Italy
Italian_CTRL
Greek
Attica_Greece
Greek_CTRL
Druze
Palestinian
Iranian
Jordanian
Armenian
Samaritan
Iraqi_Jew
Yemenite_Jew
GBR_UK
French
Bulgarian
Hungarian
Czech
Polish
Albanian
Turkish_SGDP
Turkish
Turkish_CTRL
Italian_GS

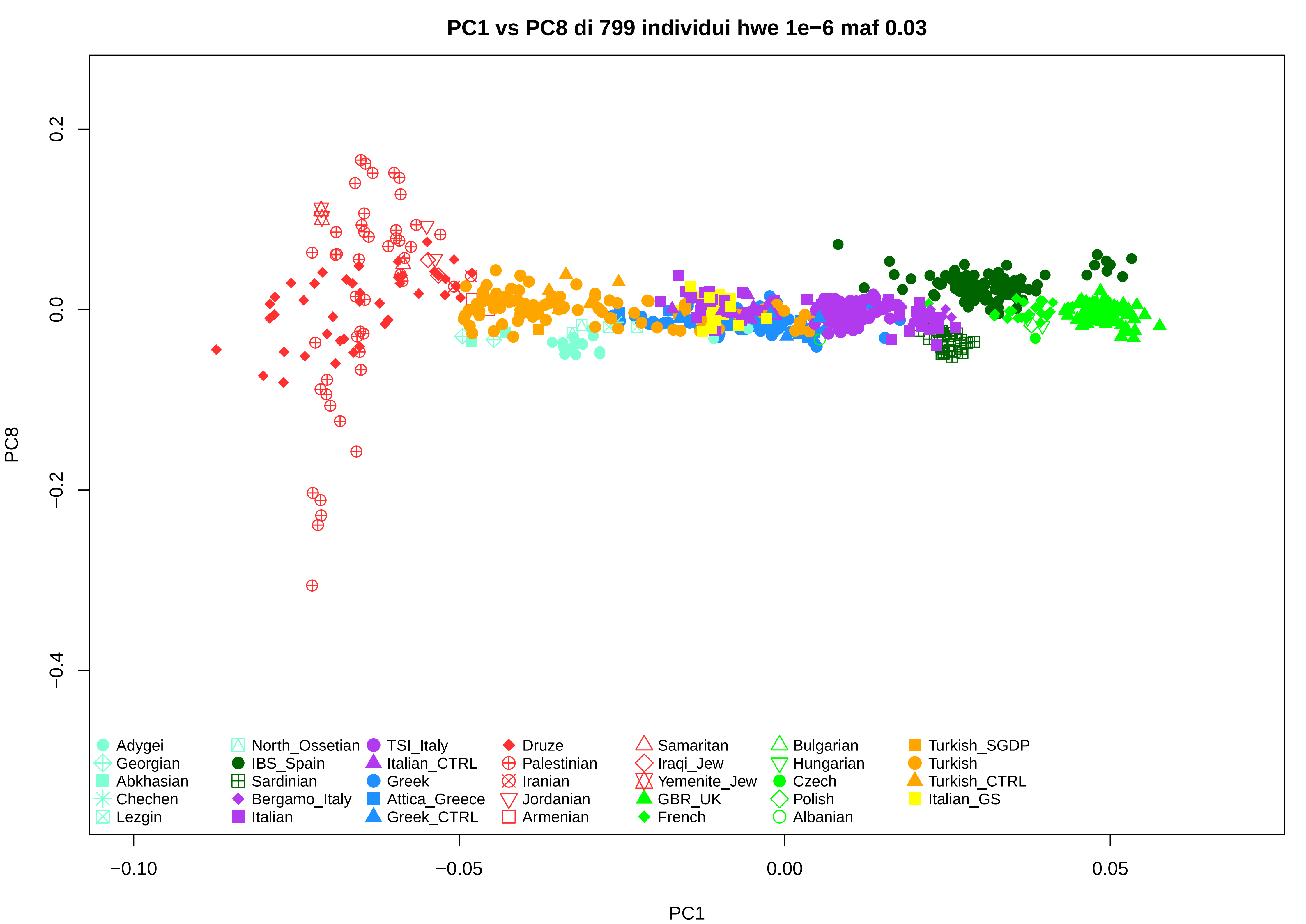

PC1 vs PC8 di 799 individui hwe 1e−6 maf 0.03
PC8
0.2
0.0
−0.2
−0.4
PC1
−0.10
−0.05
0.00
0.05
Adygei
Georgian
Abkhasian
Chechen
Lezgin
North_Ossetian
IBS_Spain
Sardinian
Bergamo_Italy
Italian
TSI_Italy
Italian_CTRL
Greek
Attica_Greece
Greek_CTRL
Druze
Palestinian
Iranian
Jordanian
Armenian
Samaritan
Iraqi_Jew
Yemenite_Jew
GBR_UK
French
Bulgarian
Hungarian
Czech
Polish
Albanian
Turkish_SGDP
Turkish
Turkish_CTRL
Italian_GS

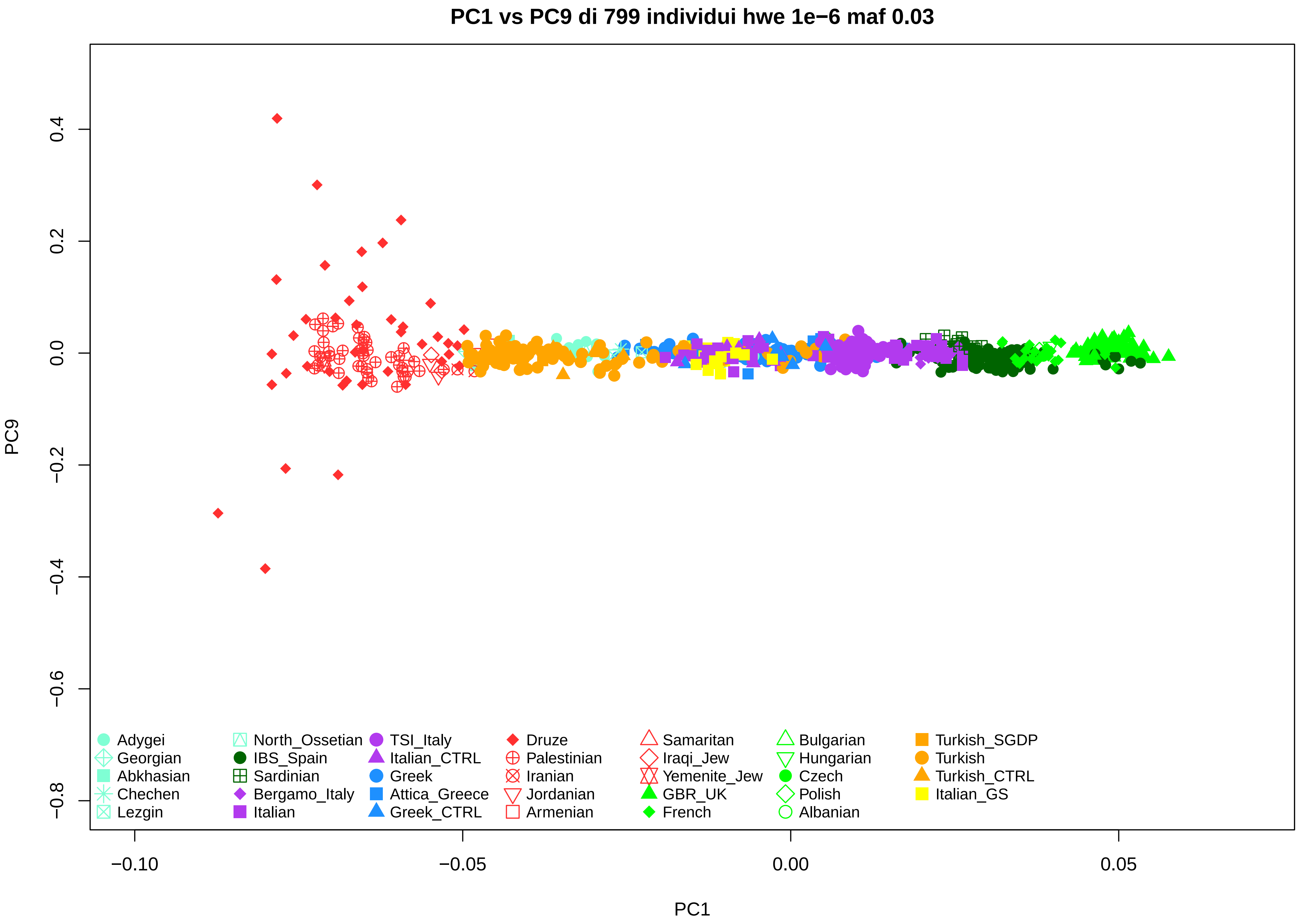

PC1 vs PC9 di 799 individui hwe 1e−6 maf 0.03
PC9
PC1
0.4
0.2
0.0
−0.2
−0.4
−0.6
−0.8
−0.10
−0.05
0.00
0.05
Adygei
Georgian
Abkhasian
Chechen
Lezgin
North_Ossetian
IBS_Spain
Sardinian
Bergamo_Italy
Italian
TSI_Italy
Italian_CTRL
Greek
Attica_Greece
Greek_CTRL
Druze
Palestinian
Iranian
Jordanian
Armenian
Samaritan
Iraqi_Jew
Yemenite_Jew
GBR_UK
French
Bulgarian
Hungarian
Czech
Polish
Albanian
Turkish_SGDP
Turkish
Turkish_CTRL
Italian_GS

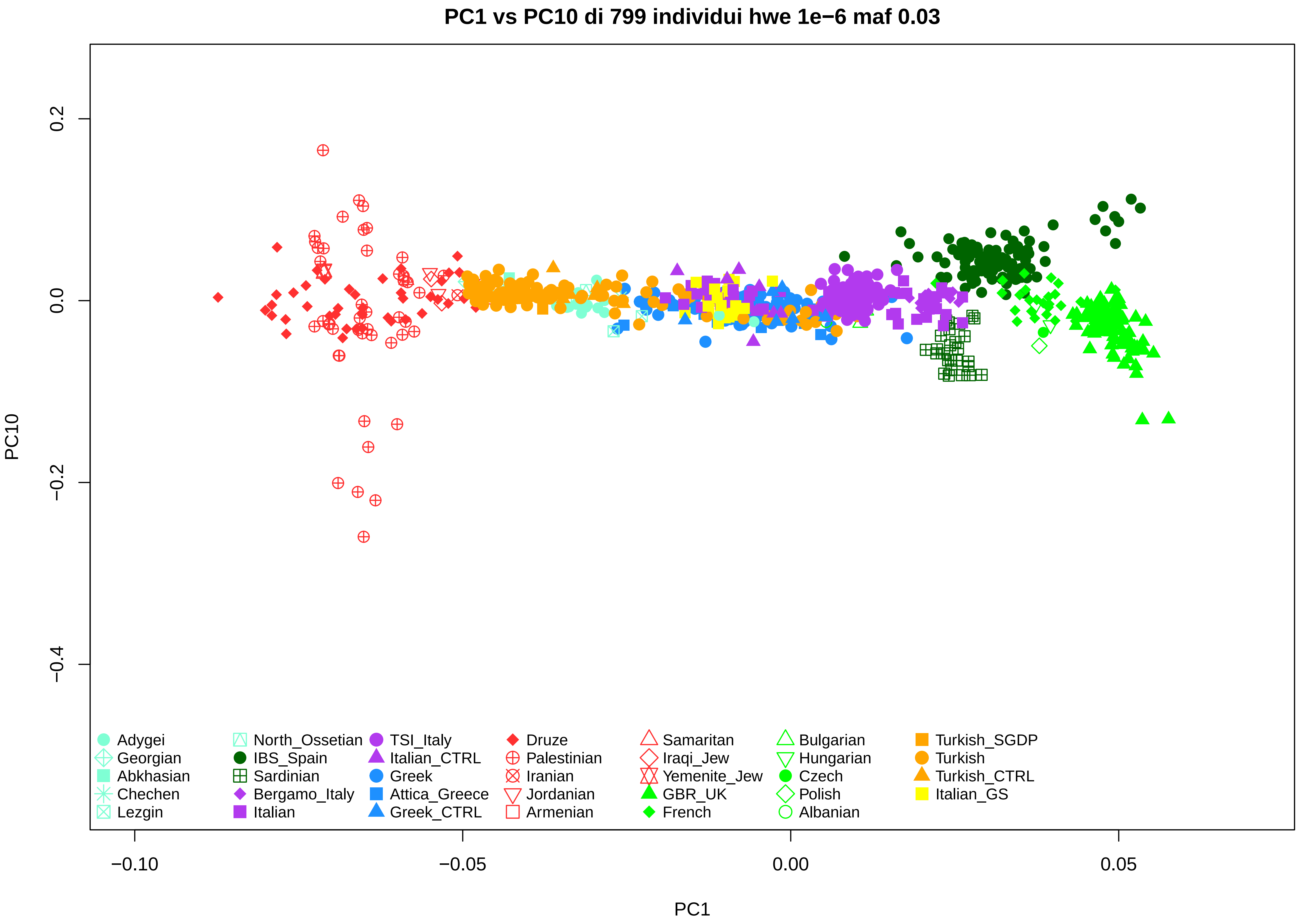
PC1 vs PC10 di 799 individui hwe 1e−6 maf 0.03
PC10
PC1
0.2
0.0
−0.2
−0.4
−0.10
−0.05
0.00
0.05
Adygei
Georgian
Abkhasian
Chechen
Lezgin
North_Ossetian
IBS_Spain
Sardinian
Bergamo_Italy
Italian
TSI_Italy
Italian_CTRL
Greek
Attica_Greece
Greek_CTRL
Druze
Palestinian
Iranian
Jordanian
Armenian
Samaritan
Iraqi_Jew
Yemenite_Jew
GBR_UK
French
Bulgarian
Hungarian
Czech
Polish
Albanian
Turkish_SGDP
Turkish
Turkish_CTRL
Italian_GS

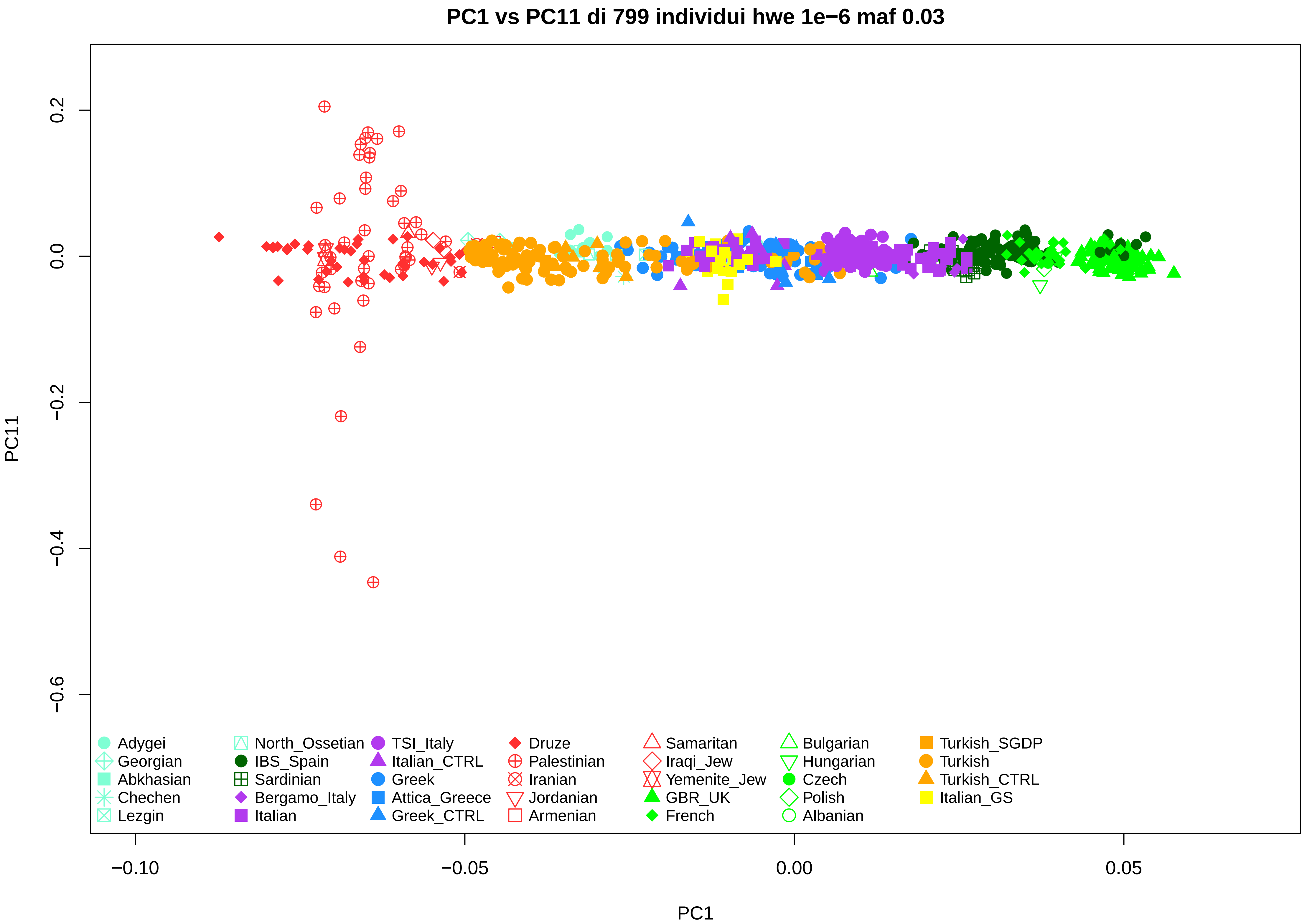
PC1 vs PC11 di 799 individui hwe 1e−6 maf 0.03
PC11
PC1
0.2
0.0
−0.2
−0.4
−0.6
−0.10
−0.05
0.00
0.05
Adygei
Georgian
Abkhasian
Chechen
Lezgin
North_Ossetian
IBS_Spain
Sardinian
Bergamo_Italy
Italian
TSI_Italy
Italian_CTRL
Greek
Attica_Greece
Greek_CTRL
Druze
Palestinian
Iranian
Jordanian
Armenian
Samaritan
Iraqi_Jew
Yemenite_Jew
GBR_UK
French
Bulgarian
Hungarian
Czech
Polish
Albanian
Turkish_SGDP
Turkish
Turkish_CTRL
Italian_GS

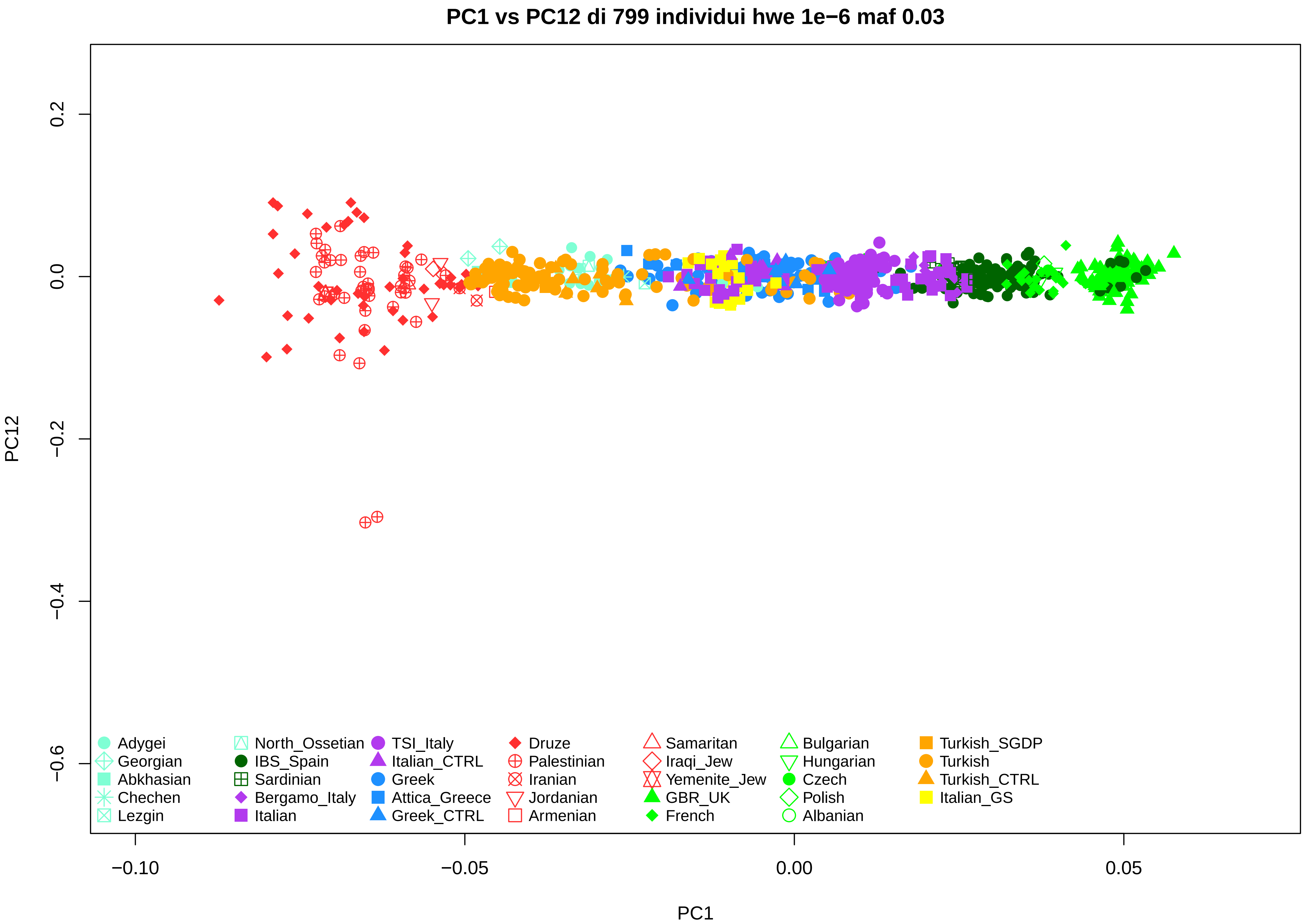
PC1 vs PC12 di 799 individui hwe 1e−6 maf 0.03
PC12
PC1
0.2
0.0
−0.2
−0.4
−0.6
−0.10
−0.05
0.00
0.05
Adygei
Georgian
Abkhasian
Chechen
Lezgin
North_Ossetian
IBS_Spain
Sardinian
Bergamo_Italy
Italian
TSI_Italy
Italian_CTRL
Greek
Attica_Greece
Greek_CTRL
Druze
Palestinian
Iranian
Jordanian
Armenian
Samaritan
Iraqi_Jew
Yemenite_Jew
GBR_UK
French
Bulgarian
Hungarian
Czech
Polish
Albanian
Turkish_SGDP
Turkish
Turkish_CTRL
Italian_GS

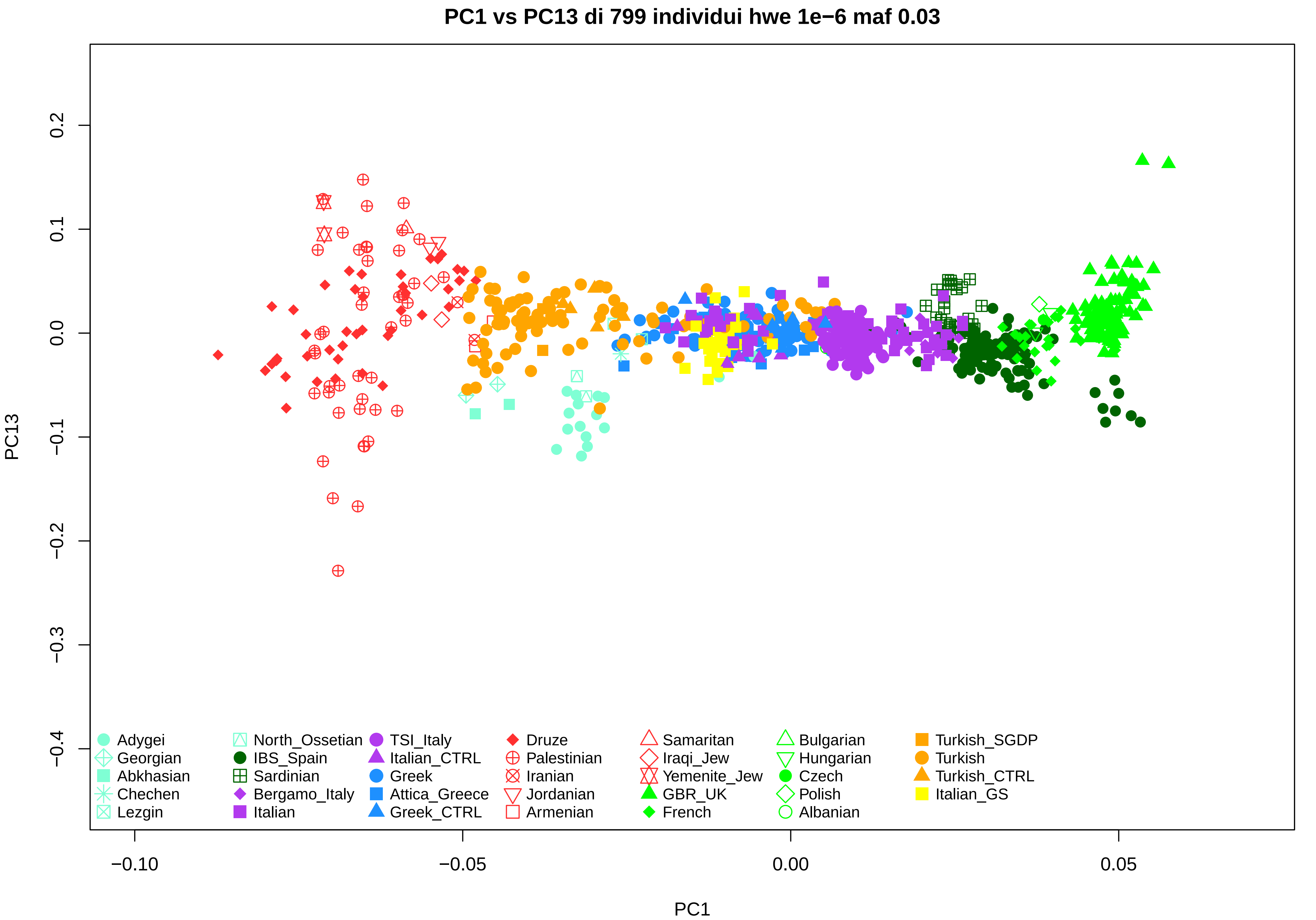
PC1 vs PC13 di 799 individui hwe 1e−6 maf 0.03
PC13
PC1
0.2
0.1
0.0
−0.1
−0.2
−0.3
−0.4
−0.10
−0.05
0.00
0.05
Adygei
Georgian
Abkhasian
Chechen
Lezgin
North_Ossetian
IBS_Spain
Sardinian
Bergamo_Italy
Italian
TSI_Italy
Italian_CTRL
Greek
Attica_Greece
Greek_CTRL
Druze
Palestinian
Iranian
Jordanian
Armenian
Samaritan
Iraqi_Jew
Yemenite_Jew
GBR_UK
French
Bulgarian
Hungarian
Czech
Polish
Albanian
Turkish_SGDP
Turkish
Turkish_CTRL
Italian_GS

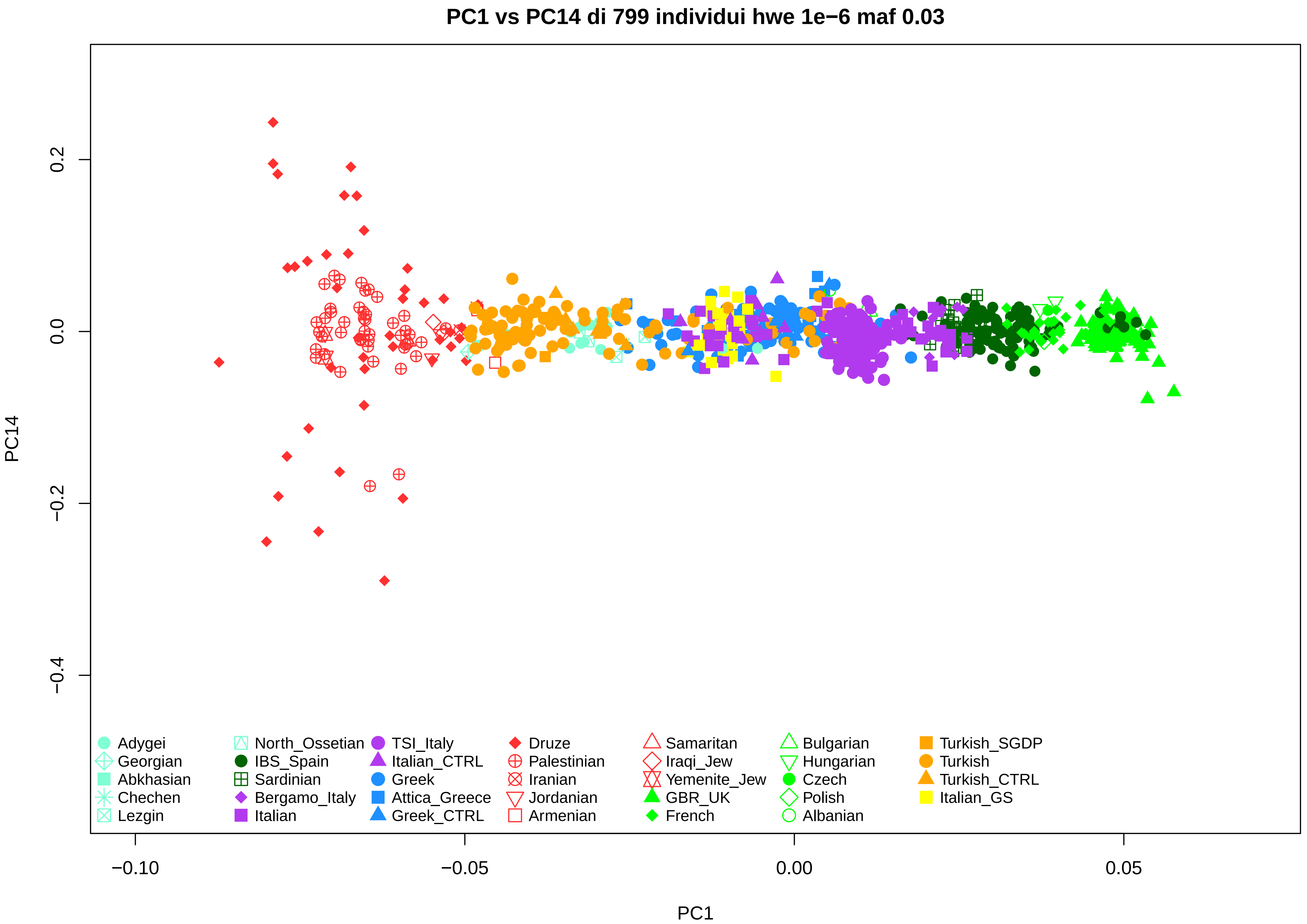
PC1 vs PC14 di 799 individui hwe 1e−6 maf 0.03
PC14
0.2
0.0
−0.2
−0.4
PC1
−0.10
−0.05
0.00
0.05
Adygei
Georgian
Abkhasian
Chechen
Lezgin
North_Ossetian
IBS_Spain
Sardinian
Bergamo_Italy
Italian
TSI_Italy
Italian_CTRL
Greek
Attica_Greece
Greek_CTRL
Druze
Palestinian
Iranian
Jordanian
Armenian
Samaritan
Iraqi_Jew
Yemenite_Jew
GBR_UK
French
Bulgarian
Hungarian
Czech
Polish
Albanian
Turkish_SGDP
Turkish
Turkish_CTRL
Italian_GS

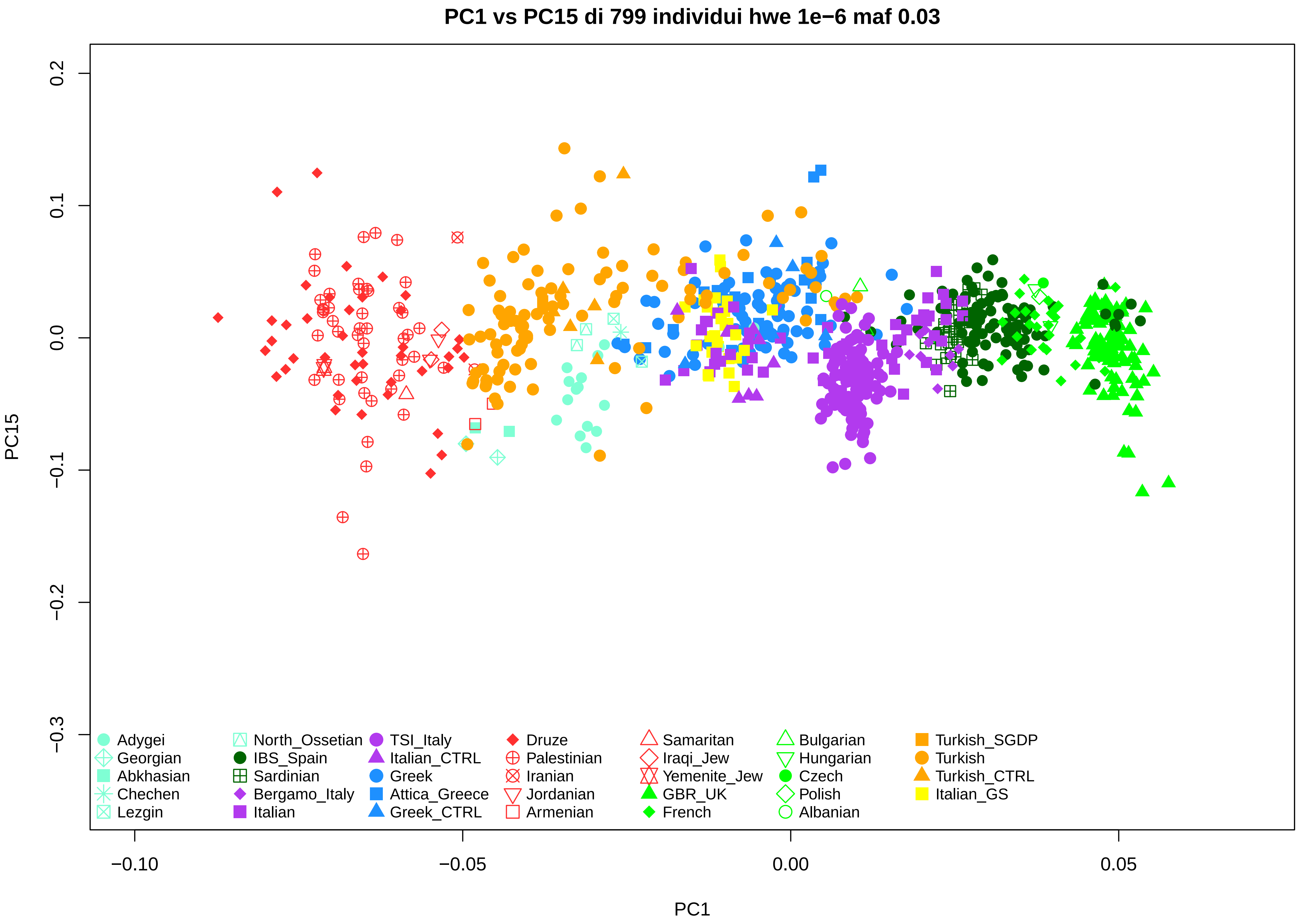

PC1 vs PC15 di 799 individui hwe 1e−6 maf 0.03
PC15
PC1
0.2
0.1
0.0
−0.1
−0.2
−0.3
−0.10
−0.05
0.00
0.05
Adygei
Georgian
Abkhasian
Chechen
Lezgin
North_Ossetian
IBS_Spain
Sardinian
Bergamo_Italy
Italian
TSI_Italy
Italian_CTRL
Greek
Attica_Greece
Greek_CTRL
Druze
Palestinian
Iranian
Jordanian
Armenian
Samaritan
Iraqi_Jew
Yemenite_Jew
GBR_UK
French
Bulgarian
Hungarian
Czech
Polish
Albanian
Turkish_SGDP
Turkish
Turkish_CTRL
Italian_GS

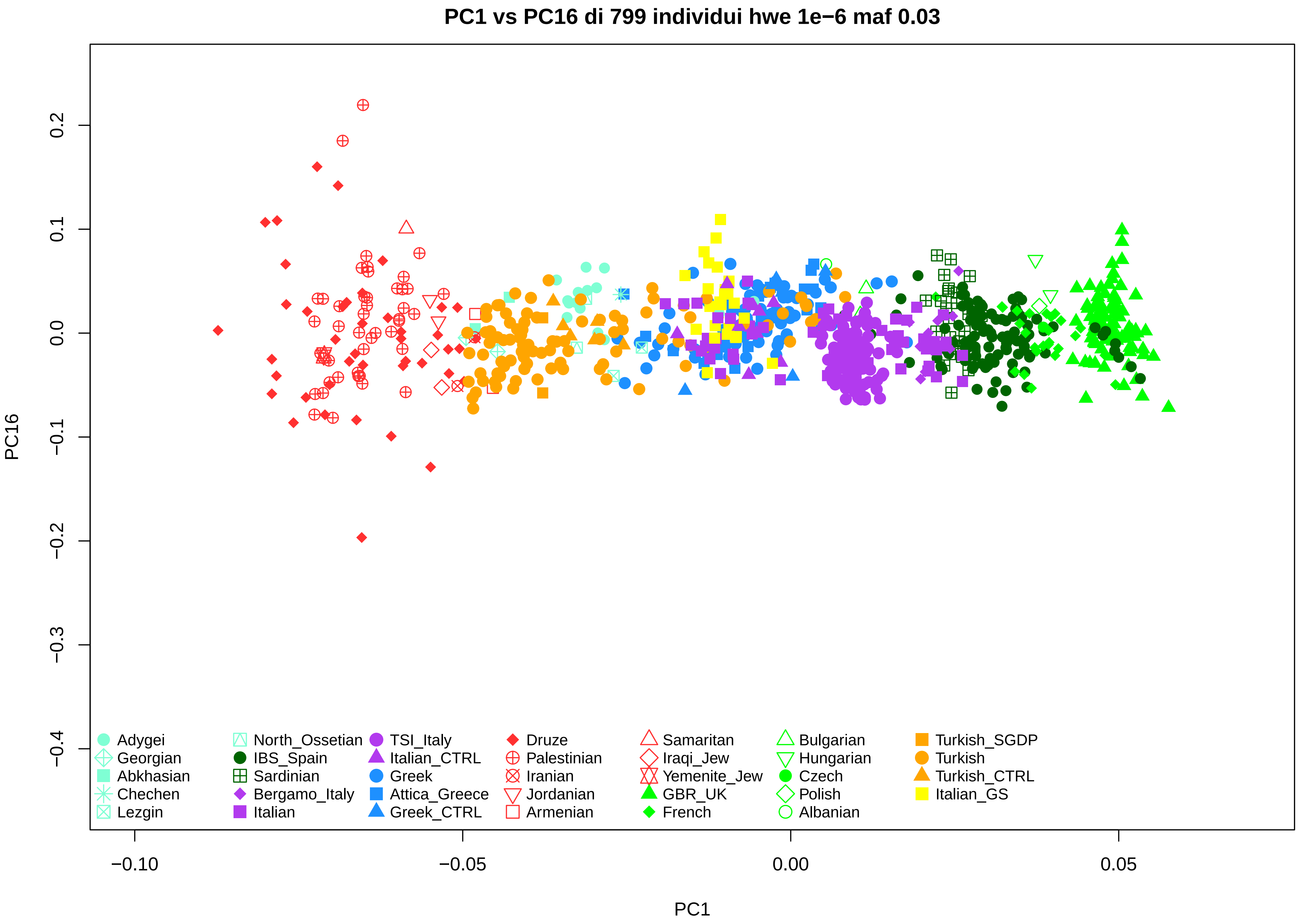
PC1 vs PC16 di 799 individui hwe 1e−6 maf 0.03
PC16
PC1
0.2
0.1
0.0
−0.1
−0.2
−0.3
−0.4
−0.10
−0.05
0.00
0.05
Adygei
Georgian
Abkhasian
Chechen
Lezgin
North_Ossetian
IBS_Spain
Sardinian
Bergamo_Italy
Italian
TSI_Italy
Italian_CTRL
Greek
Attica_Greece
Greek_CTRL
Druze
Palestinian
Iranian
Jordanian
Armenian
Samaritan
Iraqi_Jew
Yemenite_Jew
GBR_UK
French
Bulgarian
Hungarian
Czech
Polish
Albanian
Turkish_SGDP
Turkish
Turkish_CTRL
Italian_GS

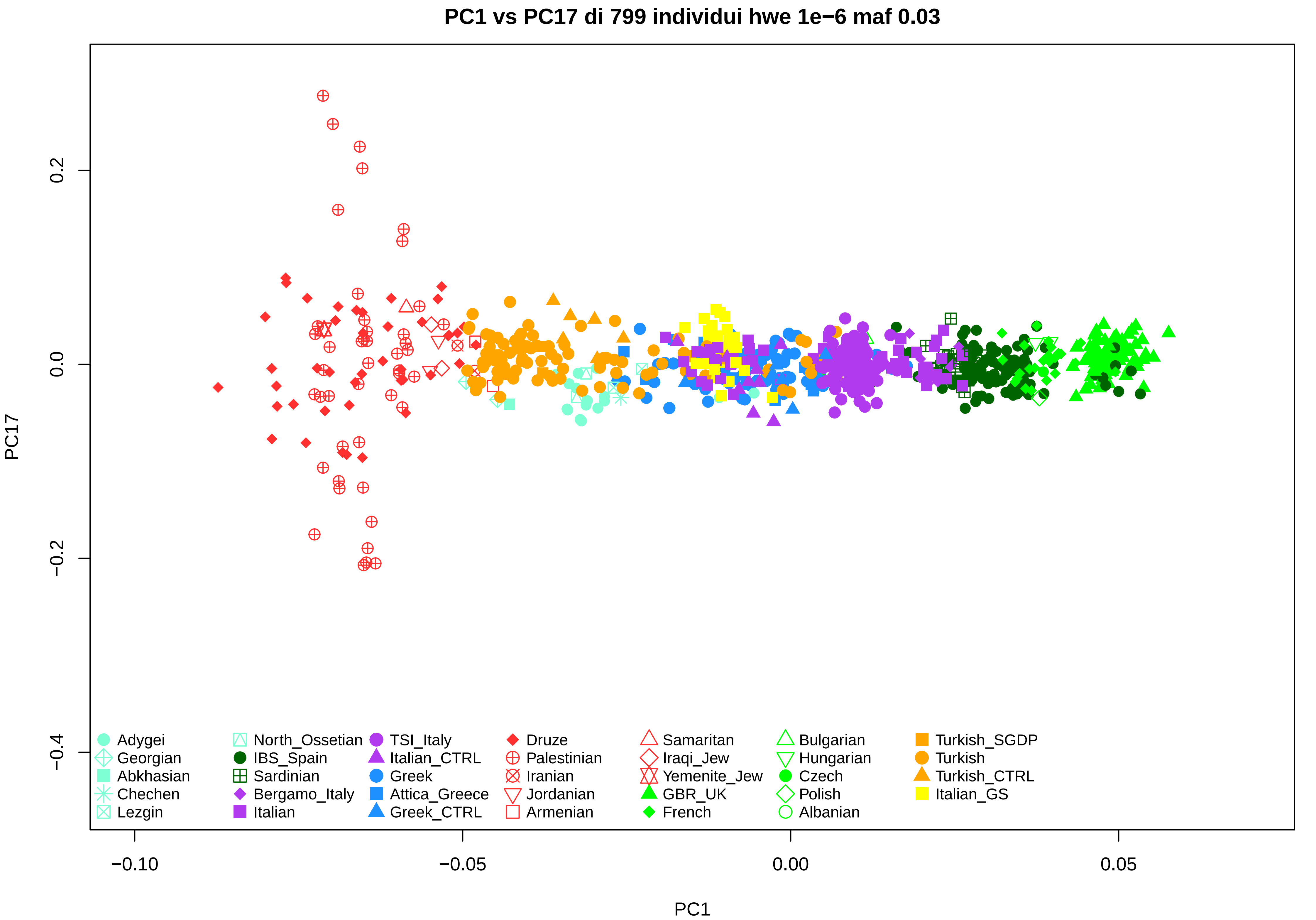
PC1 vs PC17 di 799 individui hwe 1e−6 maf 0.03
PC17
PC1
0.2
0.0
−0.2
−0.4
−0.10
−0.05
0.00
0.05
Adygei
Georgian
Abkhasian
Chechen
Lezgin
North_Ossetian
IBS_Spain
Sardinian
Bergamo_Italy
Italian
TSI_Italy
Italian_CTRL
Greek
Attica_Greece
Greek_CTRL
Druze
Palestinian
Iranian
Jordanian
Armenian
Samaritan
Iraqi_Jew
Yemenite_Jew
GBR_UK
French
Bulgarian
Hungarian
Czech
Polish
Albanian
Turkish_SGDP
Turkish
Turkish_CTRL
Italian_GS

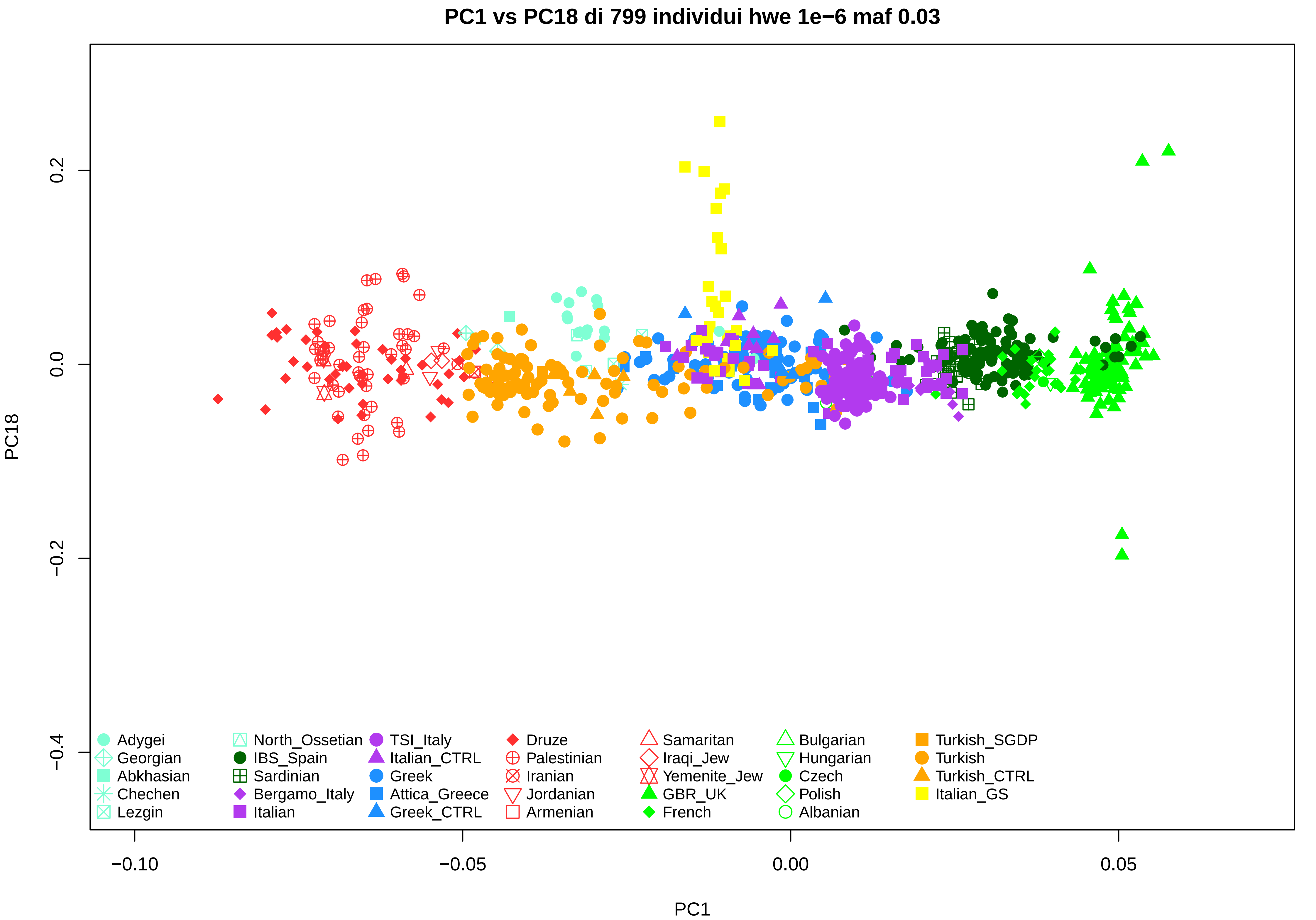
PC1 vs PC18 di 799 individui hwe 1e−6 maf 0.03
PC18
PC1
0.2
0.0
−0.2
−0.4
−0.10
−0.05
0.00
0.05
Adygei
Georgian
Abkhasian
Chechen
Lezgin
North_Ossetian
IBS_Spain
Sardinian
Bergamo_Italy
Italian
TSI_Italy
Italian_CTRL
Greek
Attica_Greece
Greek_CTRL
Druze
Palestinian
Iranian
Jordanian
Armenian
Samaritan
Iraqi_Jew
Yemenite_Jew
GBR_UK
French
Bulgarian
Hungarian
Czech
Polish
Albanian
Turkish_SGDP
Turkish
Turkish_CTRL
Italian_GS

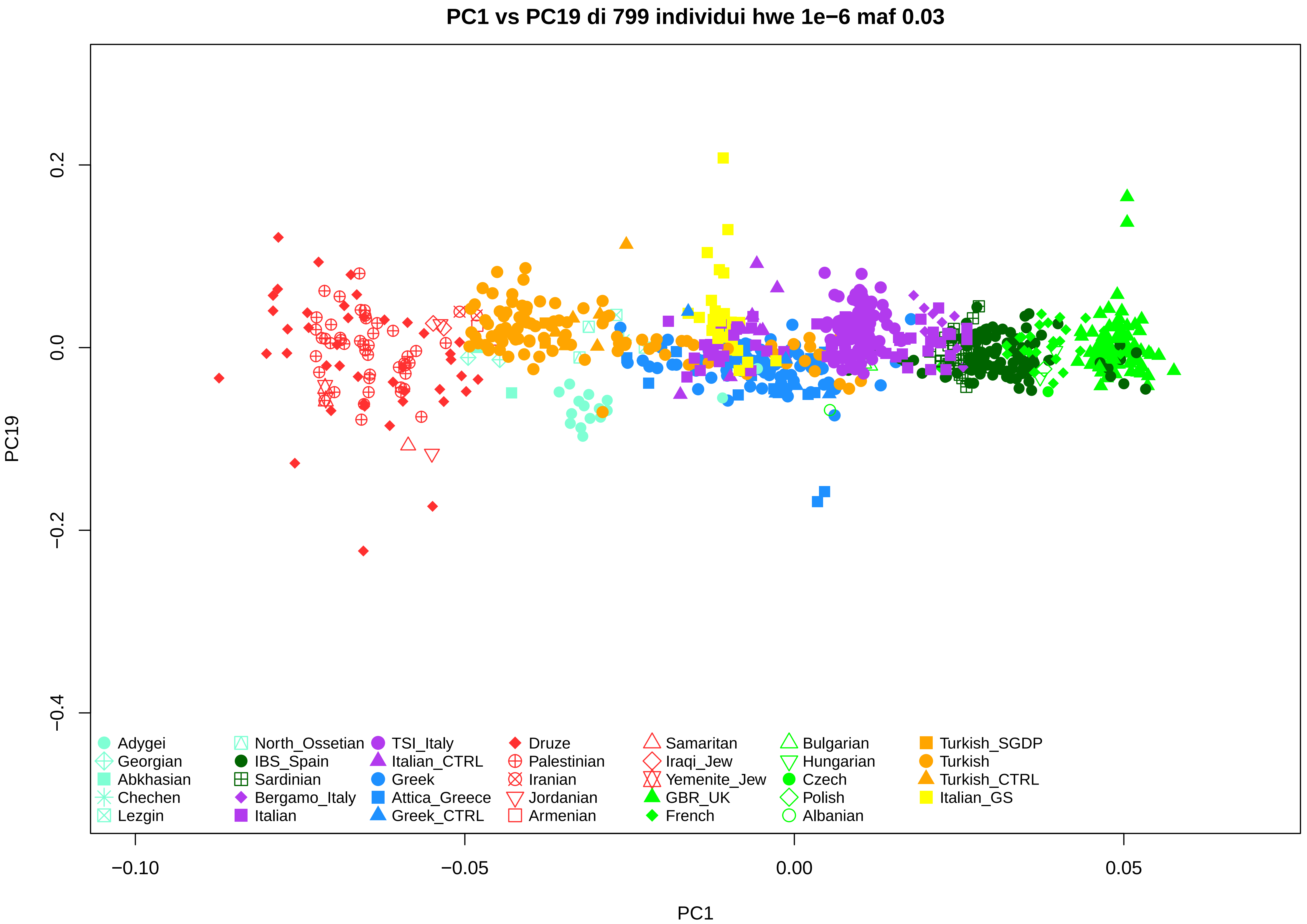
PC1 vs PC19 di 799 individui hwe 1e−6 maf 0.03
PC19
PC1
0.2
0.0
−0.2
−0.4
−0.10
−0.05
0.00
0.05
Adygei
Georgian
Abkhasian
Chechen
Lezgin
North_Ossetian
IBS_Spain
Sardinian
Bergamo_Italy
Italian
TSI_Italy
Italian_CTRL
Greek
Attica_Greece
Greek_CTRL
Druze
Palestinian
Iranian
Jordanian
Armenian
Samaritan
Iraqi_Jew
Yemenite_Jew
GBR_UK
French
Bulgarian
Hungarian
Czech
Polish
Albanian
Turkish_SGDP
Turkish
Turkish_CTRL
Italian_GS

### PC1 vs PC20 di 799 individui hwe 1e−6 maf 0.03

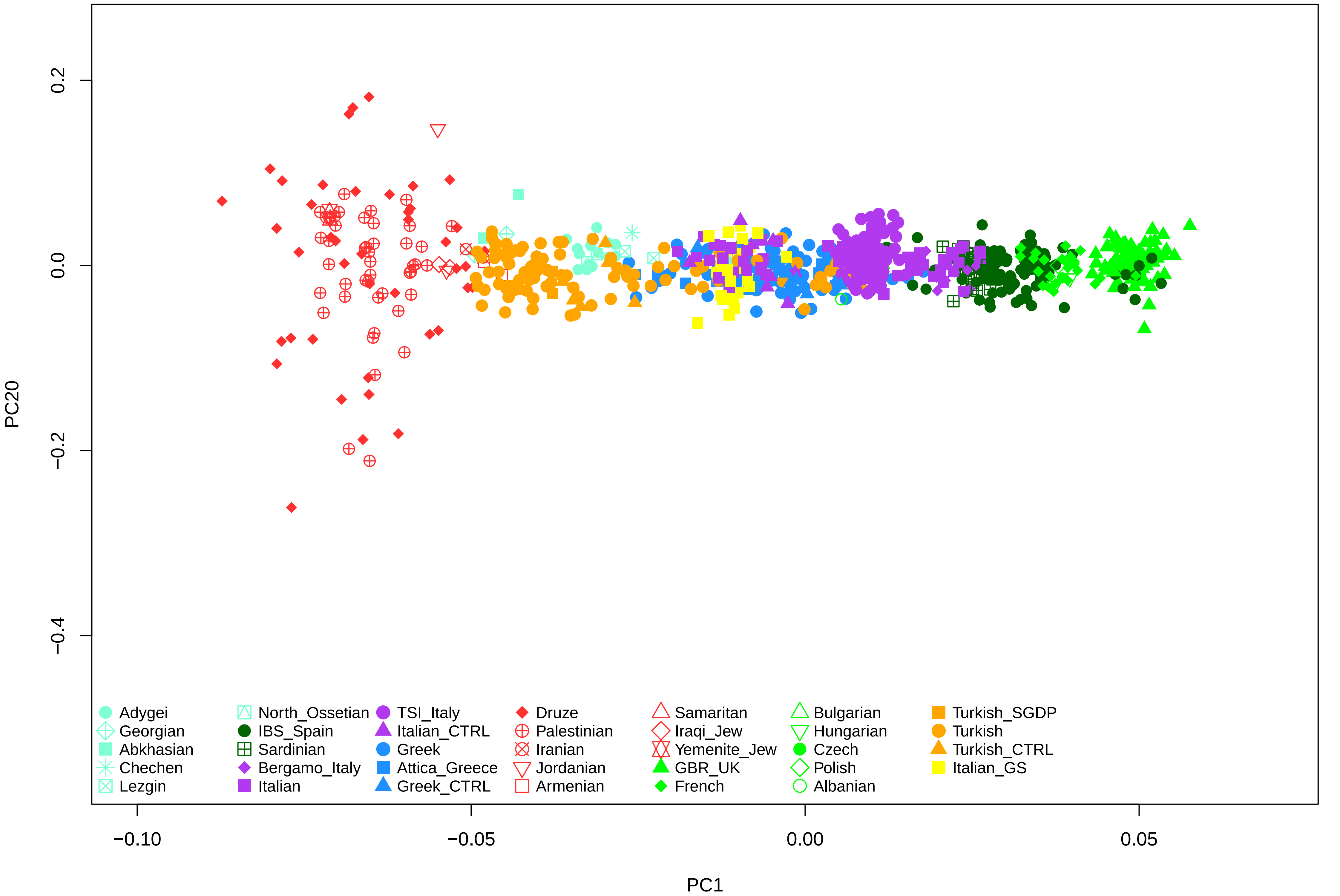
